# Novel heme-binding enables allosteric modulation in an ancient TIM-barrel glycosidase

**DOI:** 10.1101/2020.05.27.118968

**Authors:** Gloria Gamiz-Arco, Luis I. Gutierrez-Rus, Valeria A. Risso, Beatriz Ibarra-Molero, Yosuke Hoshino, Dušan Petrović, Adrian Romero-Rivera, Burckhard Seelig, Jose A. Gavira, Shina C.L. Kamerlin, Eric A. Gaucher, Jose M. Sanchez-Ruiz

**Author notes:** These authors contributed equally to this work (Gloria Gamiz-Arco, Luis I. Gutierrez-Rus).

## Abstract

Glycosidases are phylogenetically widely distributed enzymes that are crucial for the cleavage of glycosidic bonds. Here, we present the exceptional properties of a putative ancestor of bacterial and eukaryotic family-1 glycosidases. The ancestral protein shares the TIM-barrel fold with its modern descendants but displays large regions with greatly enhanced conformational flexibility. Yet, the barrel core remains comparatively rigid and the ancestral glycosidase activity is stable, with an optimum temperature within the experimental range for thermophilic family-1 glycosidases. None of the ~5500 reported crystallographic structures of ~1400 modern glycosidases show a bound porphyrin. Remarkably, the ancestral glycosidase binds heme tightly and stoichiometrically at a well-defined buried site. Heme binding rigidifies this TIM-barrel and allosterically enhances catalysis. Our work demonstrates the capability of ancestral protein reconstructions to reveal valuable but unexpected biomolecular features when sampling distant sequence space. The potential of the ancestral glycosidase as a scaffold for custom catalysis and biosensor engineering is discussed.

## INTRODUCTION

Pauling and Zuckerkandl proposed in 1963 that the sequences of modern protein homologs could be used to reconstruct the sequences of their ancestors^1^. While this was mostly only a theoretical possibility in the mid-twentieth century, ancestral sequence reconstruction has become a standard procedure in the twenty-first century, due to advances in bioinformatics and phylogenetics, together with the availability of increasingly large sequence databases. Indeed, in the last ~20 years, proteins encoded by reconstructed ancestral sequences (“resurrected” ancestral proteins, in the common jargon of the field) have been extensively used as tools to address important problems in molecular evolution^2,3^. In addition, a new and important implication of sequence reconstruction is currently emerging linked to the realization that resurrected ancestral proteins may display properties that are desirable in scaffolds for enzyme engineering^4–6^. For instance, high stability and substrate/catalytic promiscuity have been described in a number of ancestral resurrection studies^5,7^. These two features are known contributors to protein evolvability^8,9^, which points to the potential of resurrected ancestral proteins as scaffolds for the engineering of new functionalities^4,10^.

More generally, reconstruction studies that target ancient phylogenetic nodes typically predict extensive sequence differences with respect to their modern proteins. Consequently, proteins encoded by the reconstructed sequences may potentially display altered or unusual properties. Regardless of the possible evolutionary implications, it is of interest, therefore, to investigate which properties of putative ancestral proteins may differ from those of their modern counterparts and to explore whether and how these ancestral properties may lead to new possibilities in biotechnological applications. Here, we apply ancestral sequence reconstruction to a family of well-known and extensively characterized enzymes. Furthermore, these enzymes display 3D-structures based on the highly common and widely studied TIM-barrel fold, a fold which is both ubiquitous and highly evolvable^11–13^. Yet, we find upon ancestral resurrection a diversity of unusual and unexpected biomolecular properties that suggest new engineering possibilities that go beyond the typical applications of protein family being characterized.

Glycosidases catalyze the hydrolysis of glycosidic bonds in a wide diversity of molecules^14^. The process typically follows a Koshland mechanism based on two catalytic carboxylic acid residues and, with very few exceptions, does not involve cofactors. Glycosidic bonds are very stable and have an extremely low rate of spontaneous hydrolysis^15^. Glycosidases accelerate their hydrolysis up to ~17 orders of magnitude, being some of the most proficient enzymes functionally characterized^16^. Glycosidases are phylogenetically widely distributed enzymes. It has been estimated, for instance, that about 3% of the human genome encodes glycosidases^17^. They have been extensively studied, partly because of their many biotechnological applications^14^. Detailed information about glycosidases is collected in the public CAZy database (http://www.cazy.org)^18^ and the connected CAZypedia resource (http://www.cazypedia.org/)^19^. At the time of our study, glycosidases are classified into 167 families on the basis of sequence similarity. Since perturbations of protein structure during evolution typically occur more slowly than sequences change^20^, it is not surprising that the overall protein fold is conserved within each family. Forty eight of the currently described glycosidase families display a fold consistent with the TIM barrel architecture.

Here, we study family 1 glycosidases, which are of the classical TIM-barrel fold. Family 1 glycosidases (GH1) commonly function as β-glucosidases and β-galactosidases, although other activities are also found in the family^21^. We focus on a putative ancestor of modern bacterial and eukaryotic enzymes and find a number of highly unusual properties that clearly differentiate the ancestor from the properties of its modern descendants. The ancestral glycosidase thus displays much-enhanced conformational flexibility in large regions of its structure. This flexibility, however, does not compromise stability as shown by the ancestral optimum activity temperature which is within the typical range for family 1 glycosidases from thermophilic organisms. Unexpectedly, the ancestral glycosidase binds heme tightly at a well-defined site in the structure with concomitant allosteric increase in enzyme activity. Neither metallo-porphyrin binding nor allosteric modulation appear to have been reported for any modern glycosidases, despite of the fact that these enzymes have been extensively characterized. Overall, this work demonstrates the potential of ancestral reconstruction as a tool to explore sequence space to generate combinations of properties that are unusual or unexpected compared to the repertoire from modern proteins.

## RESULTS

### Ancestral sequence reconstruction

Ancestral sequence reconstruction (ASR) was performed based on a phylogenetic analysis of family 1 glycosidases (GH1) protein sequences (see the Methods for details). GH1 protein homologs are widely distributed in all three domains of life and representative sequences were collected from each domain, including characterized GH1 sequences obtained from CAZy as well as homologous sequences contained in GenBank. The phylogeny of GH1 homologs consists of four major clades (Figure 1A and Figure S1). One clade is composed mainly of archaea and bacteria from the recently proposed Candidate Phyla Radiation (CPR)^22^, while the other three clades include bacteria and eukaryotes. The archaeal/CPR clade largely contains uncharacterized proteins and was thus excluded from further analysis. In the bacterial/eukaryotic clades, eukaryotic homologs form a monophyletic clade within bacterial homologs. For our current study, the common ancestors of bacterial and eukaryotic homologs are selected for ASR analysis (N72, N73 and N125) because many homologs have been characterized and there is substrate diversity between the enzymes in the different clades.

**Figure 1.**
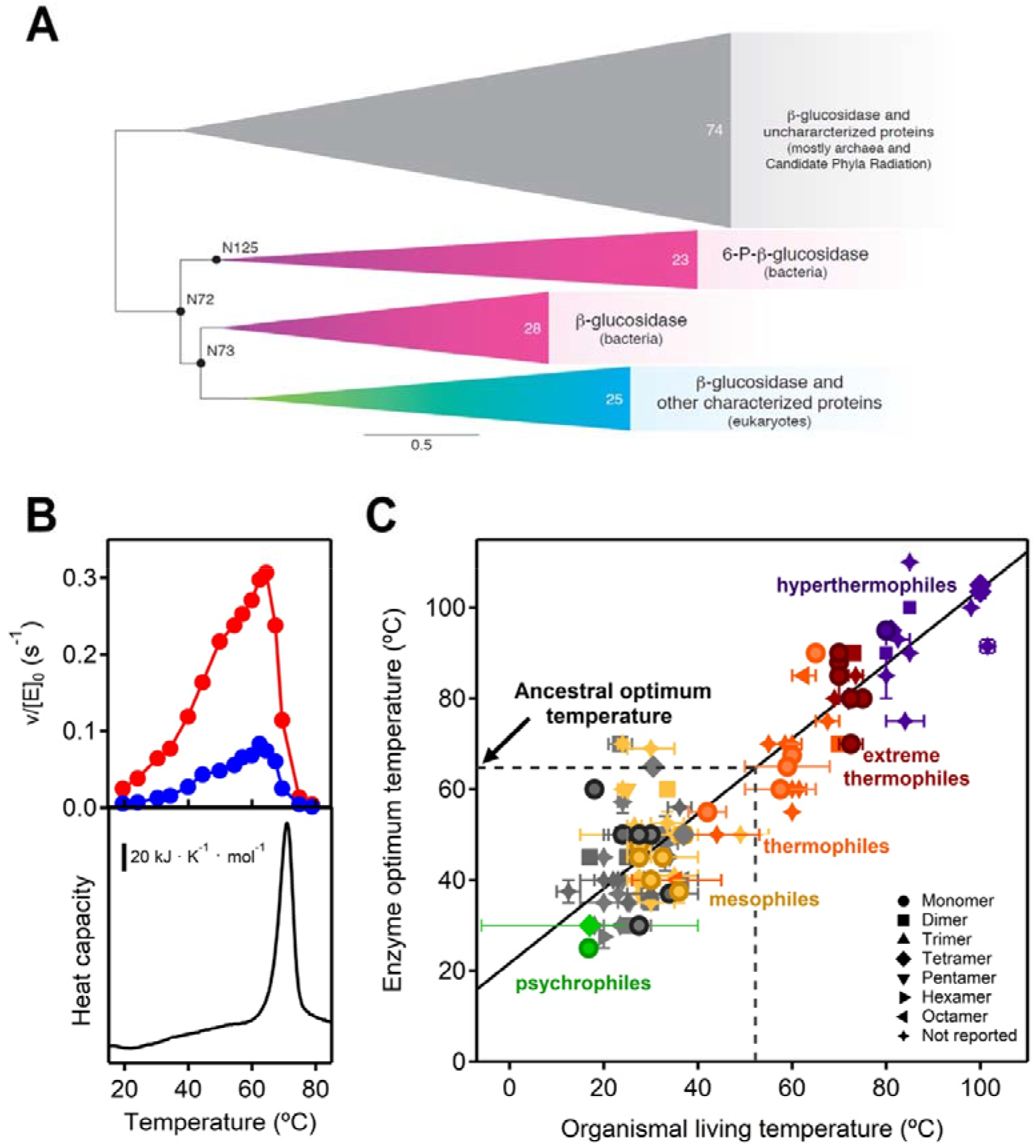
Ancestral sequence reconstruction of family 1 glycosidases (GH1) and assessment of ancestral stability. (A) Bayesian phylogenetic tree of GH1 protein sequences using 150 representative sequences. Triangles correspond to four major well-supported clades (see supplemental Figure S1 for nodal support) with common functions indicated. Numbers inside the triangles correspond to the number of sequences in each clade. Scale bar represents 0.5 amino acid replacements per site per unit evolutionary time. Reconstructed ancestral sequences were inferred at the labelled nodes and the protein at node 72 was exhaustively characterized. (B) Determination of the optimum temperature for the ancestral glycosidase (upper panel) using two different substrates 4-nitrophenyl-β-D-glucopyranoside (red) and 4-nitrophenyl-β-D-galactopyranoside (blue). The lower panel shows a differential scanning calorimetry profile for the ancestral glycosidase. Clearly, the activity drop observed at high temperature (upper panel) corresponds to the denaturation of the protein, as seen in the lower panel. (C) Plot of enzyme optimum temperature versus living temperature of the host organism for modern family 1 glycosidases. Data (Table S2) are derived from literature searches, as described in Methods. Color code denotes the organisms that published literature describes as hyperthermophiles, extreme thermophiles, thermophiles, mesophiles, psychrophiles; gray color is used for organisms that have not been thus classified (plants that live at moderate temperatures in most cases. The line is a linear-squares fit (T_OPT_=21.66+0.824T_LIVING_). Correlation coefficient is 0.90 and p~1.6·10^−45^ (probability that the correlation results from chance). An environmental temperature of about 52 °C can be estimated from the optimum temperature of the ancestral glycosidase.

### Selection of an ancestral glycosidase for experimental characterization

We prepared, synthesized and purified the proteins encoded by the most probabilistic sequences at nodes N72, N73 and N125. While the three proteins were active and stable, we found that those corresponding to N73 and N125 had a tendency to aggregate over time. We therefore selected the resurrected protein from node 72 for exhaustive biochemical and biophysical characterization. For the sake of simplicity, we will subsequently refer to this protein as the ancestral glycosidase in the current study.

It is important to note that the sequence of the ancestral glycosidase differs considerably from the sequences of modern proteins. The set of modern sequences used as a basis for ancestral reconstruction span a range of sequence identity (26-59%) with the ancestral glycosidase (Table S1). Also, using the ancestral sequence as the query of a BLAST search in several databases (non-redundant protein sequences, UniProtKB/Swiss-Prot, Protein Data Bank, Metagenomic proteins) yields a closest hit with only a 62% sequence identity to the ancestral glycosidase. These sequence differences translate into unexpected biomolecular properties.

### Stability

As it is customary in the glycosidase field, we assessed the stability of the ancestral glycosidase using profiles of activity versus temperature determined by incubation assays^23^. These profiles typically reveal a well-defined optimum activity temperature (Figure 1B) as a result of the concurrence of two effects. At low temperatures, the expected Arrhenius-like increase of activity with temperature is observed. At high temperatures, protein denaturation occurs and causes a sharp decrease in activity. For the ancestral glycosidase, this interpretation is supported by differential scanning calorimetry data (lower panel in Figure 1B) which show a denaturation transition that spans the temperature range in which the activity drops sharply.

The profiles of activity versus temperature (Figure 1B) show a sharp maximum at 65 °C for the optimum activity temperature of the ancestral glycosidase. In order to ascertain the implications of this value for the evaluation of the ancestral stability, we have searched the literature on family 1 glycosidases for reported optimum temperature values (see Methods for details and Table S2 for the results of the search). The values for ~130 modern enzymes show a good correlation with the environmental temperature of their respective host organisms (Figure 1C). Therefore, the enzyme optimum temperature is an appropriate reflection of stability in an environmental context for this protein family. The optimum temperature value for ancestral glycosidase is within the experimental range of optimum activity temperatures for family 1 glycosidases from thermophilic organisms and it is consistent with an ancestral environmental temperature of about 52 °C (Figure 1C).

### Conformational flexibility

Remarkably, despite its “thermophilic” stability, large regions in the structure of the ancestral glycosidase are flexible and/or unstructured, as demonstrated by both experiment and computation (Figure 2).

**Figure 2.**
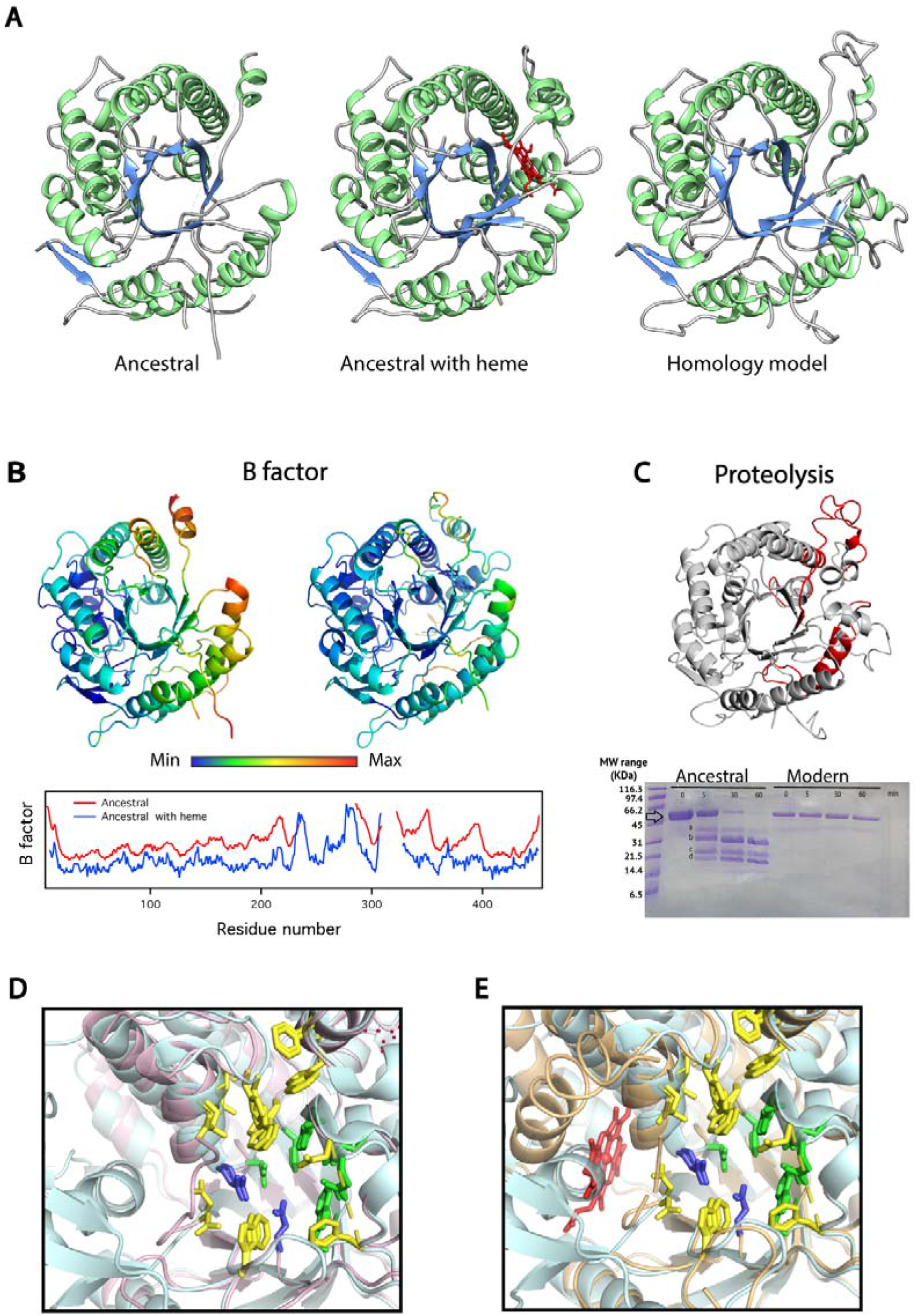
3D-Structure of the ancestral glycosidase as determined by X-ray crystallography. (A) Comparison between the ancestral structure determined in the absence (right) and presence (middle) of bound heme (red) and a homology model constructed as described in Supporting Information. The comparison visually reveals the missing sections in the electronic density of ancestral protein, mostly in the protein without heme bound. (B) 3D-structure of the ancestral protein without and with heme bound color-labeled according to normalized B-factor value and profiles of normalized B-factor versus residue number for the ancestral protein without (red) and with (blue) bound heme. Values are not shown for the sections that are missing in the experimental structures. (C) Proteolysis experiments with the ancestral glycosidase and the modern glycosidase from *Halothermothrix orenii*. The major fragments are labeled *a*, *b*, *c* and *d*. Mass spectrometry of the fragments predicts cleavage points within the red labeled sections in the shown structure. (D) Superposition of the structure of the ancestral glycosidase with that of the modern glycosidase from *Halothermothrix orenii* showing the critical active site residues. (E) Superposition of the structures of the ancestral glycosidase without and with heme bound showing the critical active site residues. In both D and E, the highlighted active-site residues include the catalytic carboxylic acid residues (blue) and the residues involved in binding of the glycone (yellow) and aglycone (green) parts of the substrate.

Proteolysis is known to provide a suitable probe of conformational diversity and the protein energy landscape^24^, since most cleavable sites are not exposed in folded compact protein states. The ancestral glycosidase is highly susceptible to proteolysis and degradation is already apparent after only a few minutes incubation at a low concentration of thermolysin (0.01 mg/mL, Figures 2C, and S2). Conversely, the modern glycosidase from the thermophilic *Halothermothrix orenii* remains essentially unaffected after several hours with the same concentration of the protease (Figure 2C) or even with a ten times larger protease concentration. These two glycosidases, modern thermophilic and putative ancestral, are monomeric (Figure S3) and display similar values for the optimum activity temperature (70 °C and 65 °C, respectively: Figure 1B and S4). Therefore, their enormously disparate susceptibilities to proteolysis can hardly be linked to differences in overall stability, but rather to enhanced conformational flexibility in the ancestral enzyme that exposes cleavable sites.

Furthermore, there is a large missing region in the electronic density map of the ancestral protein from X-ray crystallography (Figure 2A), despite the good resolution achieved (2.5 Å), while the rest of the model does agree with a homology model computed on the basis of known modern glycosidase structures (see Supplementary Methods for details). Missing regions are expected to correspond to regions of high flexibility but, in addition, flexibility is also suggested by the B-factor values in regions that are present in the ancestral structure (Figure 2B).

Lastly, molecular dynamics (MD) simulations (Figure 3) also indicate enhanced flexibility in specific regions as shown by cumulative 15 μs simulations of the substrate-free forms of the ancestral glycosidase (both with and without heme: see below) as well as the modern glycosidase from *Halothermothrix orenii* (PDB ID: 4PTV)^25^. Both ancestral and modern proteins have the same sequence length, and similar protein folds with a root mean square deviation (RMSD) difference of only 0.7Å between the structures. However, our molecular dynamics simulations indicate a clear difference in flexibility in the region spanning residues 227-334, which is highly disordered in the ancestral glycosidase but ordered and rigid in the modern glycosidase, with root mean square fluctuation (RMSF) values of <2Å (Figure 3). We also analysed the interactions formed between residues 227-334 and the rest of the protein by counting the total intramolecular hydrogen bonds formed along the MD simulations. We observe that on average, the modern glycosidase forms 115 ± 11 hydrogen bonding interactions during our simulations, whereas the ancestral glycosidase forms either 101 ± 12/101 ± 13 hydrogen bonding interactions (in the presence of heme and absence of heme, respectively). This suggests that the higher number of intramolecular hydrogen bonds formed between residues 227-334 and the protein can contribute to the reduced conformational flexibility observed in the case of the modern glycosidase, compared to the ancestral glycosidase.

**Figure 3.**
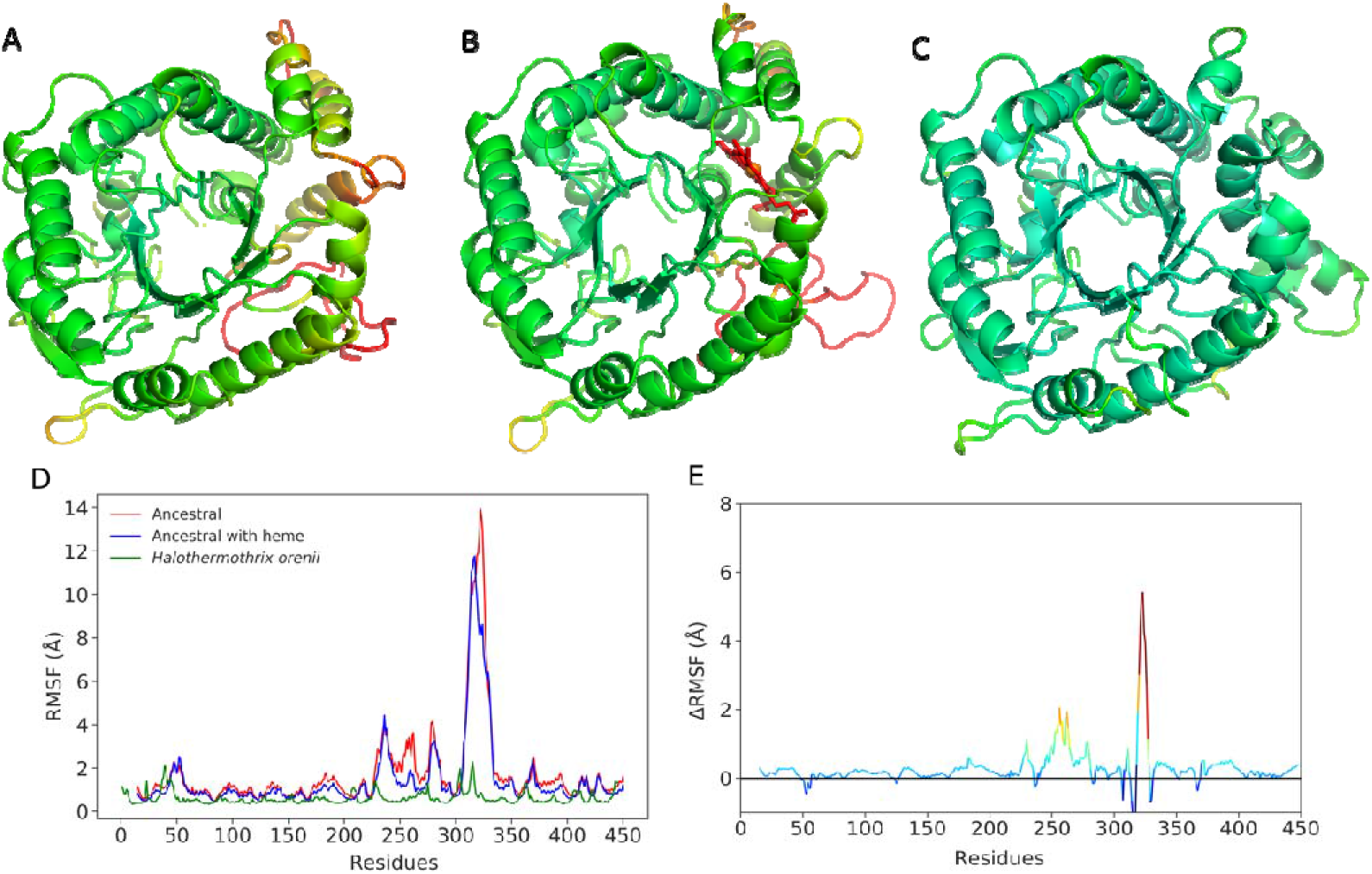
Representative snapshots from molecular dynamics simulations of ancestral and modern glycosidases, showing the ancestral glycosidase both (A) without and (B) in complex with heme, as well as (C) the corresponding modern protein from *Halothermotrix orenii*. Structures were extracted from our simulations based on [Adrian describe clustering algorithm and how cluster was selected]. All protein structures are colored by calculated root mean square fluctuations (RMSF) over the course of simulations of each system. Shown are also (D) absolute and (E) relative RMSF (Å) for each system, in the latter case relative to the heme bound structure.

It is important to note that there is a clear structural congruence between the results of the experimental and computational studies described above. That is, the missing regions in the X-ray structure (Figure 2A) match the high-flexibility regions in the molecular dynamics simulations (Figure 3) and include the proteolysis cleavage sites determined by mass spectrometry (Figure 2C). Overall, regions encompassing two alpha helices and several loops appear to be highly flexible or even unstructured in the ancestral glycosidase. The barrel core, however, remains structured and shows comparatively low conformational flexibility, which may explain the high thermal stability of the protein (see Discussion).

### Catalysis

We determined the Michaelis-Menten parameters for the ancestral enzyme with the substrates typically used to test the standard β-glucosidase and β-galactosidase activities of family 1 glycosidases (4-nitrophenyl-β-D-glucopyranoside and 4-nitrophenyl-β-D-galactopyranoside) (Figure 4 and Table S3). We also compared the results with the catalytic parameters for four modern family 1 glycosidases, specifically those from *Halothermothrix orenii*, *Marinomonas sp.* (strain MWYL1), *Saccharophagus degradans* (strain 2-40T), and *Thermotoga maritima* (Figure S5 and Table S3). Modern glycosidases are highly proficient enzymes accelerating the rate of glycoside bond hydrolysis up to about 17 orders of magnitude^16^. The ancestral enzyme appears to be less efficient and shows a turnover number about two orders of magnitude below the values for the modern glycosidases studied here (Figure 4). Still, the catalytic carboxylic acid residues as well as the residues known to be responsible for the interaction with the glycone moiety of the substrate (Marana, 2006) are present in the ancestral enzyme and appear in the static X-ray structure in a configuration similar to that observed in the modern proteins (Figure 2D). There are a few differences in the identity of the residues responsible for binding of the aglycone moiety of the substrate^26^, but these differences occur in positions that are variable in modern family 1 glycosidases (Figure S6 and Table S4). Overall, the comparatively low activity of the ancestral protein is likely linked to its conformational flexibility. That is, the protein in solution is sampling a diversity of conformations of which only a few are active towards the common substrates. From an evolutionary point of view, the comparatively low ancestral activity may reflect an early stage in the evolution of family 1 glycosidases before selection favored greater turnover (see Discussion).

**Figure 4.**
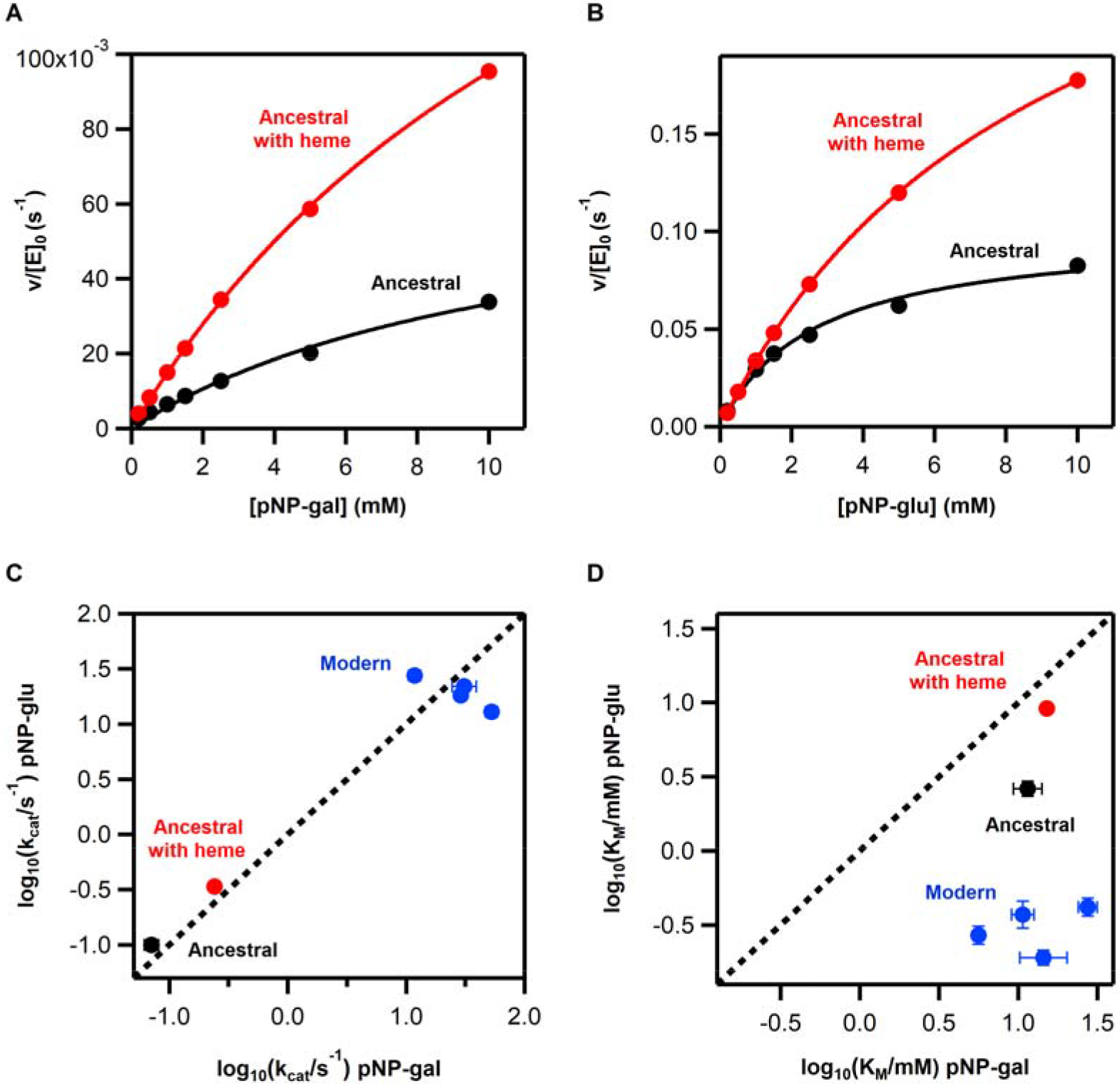
Ancestral versus modern catalysis in family 1 glycosidases. (A) and (B) Michaelis plots of rate versus substrate concentration at pH 7 and 25 °C for hydrolysis of 4-nitrophenyl-β-D-galactopyranoside (A) and 4-nitrophenyl-β-D-glucopyranoside (B) for the ancestral protein with and without heme bound. The lines are the best fits of the Michaelis-Menten equation. Michaelis plots for the 4 modern proteins studied in this work can be found in Figure S5. The values for the catalytic parameters derived from these fits are collected in Table S3. (C) Logarithm of the turnover number for a glucopyranoside substrate versus a galactopyranoside substrate. For both, the modern and the ancestral proteins, the turnover numbers for the two substrates are similar. (D) Logarithm of the Michaelis constant for a glucopyranoside substrate versus a galactopyranoside substrate. Binding is stronger for the glucopyranoside substrate in the modern proteins. pNP-glu and pNP-gal stand, respectively, for 4-nitrophenyl-β-D-glucopyranoside and 4-nitrophenyl-β-D-galactopyranoside.

Also, it is interesting to note that, although both β-glucosidase and β-galactosidase activities are typically described for family 1 glycosidases, these enzymes are commonly specialized as β-glucosidases^21^. This specialization does not occur, however, at the level of the turnover number, which is typically similar for both kinds of substrates. Instead, specialization occurs at the level of the substrate affinity, as reflected in lower values of the Michaelis constant (K_M_) for β-glucopyranoside substrates as compared to β-galactopyranoside substrates^21^. This pattern is indeed observed in the modern enzymes we have studied (Figure 4), which are described in the literature as β-glucosidases. On the other hand, this kind of specialization is not observed in the ancestral glycosidase, which shows similar *K*_*M*_*’s* for the β-glucopyranoside and the β-galactopyranoside substrates. This lack of specialization may again reflect an early stage in the evolution of family 1 glycosidases, an interpretation which would seem generally consistent with the presumed generalist nature of the most ancient enzymes^27^.

### Heme binding and allosteric modulation

Overall, it appears reasonable that our resurrected ancestral enzyme reflects an early stage in the evolution of family 1 glycosidases, perhaps following a fragment fusion event (see Discussion), at which catalysis was not yet optimized and substrate specialization had not yet evolved. The presence of a large unstructured and/or flexible regions in the ancestral structure could perhaps reflect the absence of a small molecule that binds within that region. While these proposals are speculative, the experimental results described in detail below, show that the ancestral glycosidase does bind heme tightly and stoichiometrically at a site in the flexible regions. This was a completely unexpected observation given the large number of modern glycosidases that have been characterized in the absence of any porphyrin rings.

We curiously noticed that most preparations of the ancestral glycosidase showed a light-reddish color after elution from an affinity column. UV-Vis spectra revealed the pattern of bands expected for a heme group^28^, including the Soret band at about 400 nm and, in some cases, even the weaker α and β bands (i.e., the Q bands) in the 500-600 nm region (Figure 5). From the intensity of the Soret band, a very low heme:protein ratio of about 0.02 was estimated for standard enzyme preparations, indicating that all the experiments described above were performed with essentially heme-free protein. However, the amount of bound heme in protein preparations was substantially enhanced by including hemin in the culture medium (heme with iron in the +3 oxidation state) or 5-aminolevulinic acid, the metabolic precursor of heme. Heme:protein rations of about 0.10 and 0.18, respectively, were then obtained (Figure 5A). These results suggest that the ancestral glycosidase does have the capability to bind heme, but also that, as is commonly the case with modern heme-binding proteins^29^, the limited amount of heme available in the expression host, combined with the high protein overexpression levels used, leads to low heme:protein ratios. The capability of the ancestral enzyme to bind heme was first confirmed by the *in vitro* experiments described next, and then confirmed via X-ray crystallography as subsequently described.

**Figure 5.**
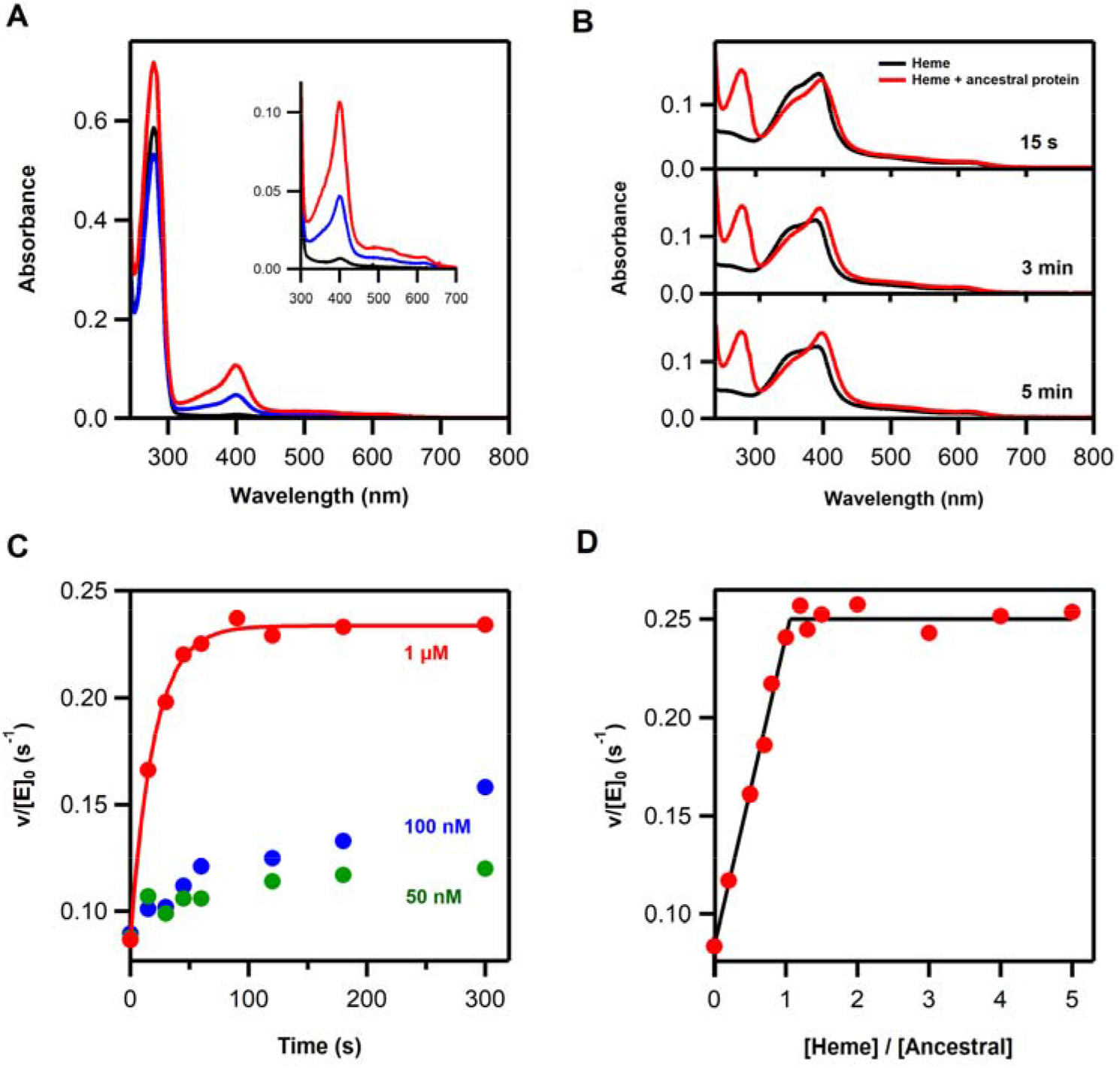
Heme binding to the ancestral glycosidase. (A) UV-VIS spectra for preparations of the ancestral glycosidase showing the protein absorption band at about 280 nm and the absorption bands due to the heme (the Soret band at about 400 nm and the Q bands at higher wavelengths). Black color is used for the protein obtained using the original purification procedure without the addition of hemin or hemin precursor. Blue and red are used to refer, respectively, to preparations in which hemin and 5-aminolevulinic acid (the metabolic precursor of heme) were added to the culture medium. (B) Binding of heme to the ancestral glycosidase *in vitro* as followed by changes in VIS spectrum. Spectra of a heme solution 1 μM in the absence (black) or presence (red) of a similar concentration of ancestral protein. The “flat” Soret band of free heme is linked to its self-association in solution, while the bound heme is monomeric and produces a sharper Soret band. (C) Kinetics of binding of heme to the ancestral glycosidase as followed by the increase in enzyme activity (rate of hydrolysis of 4-nitrophenyl-β-glucopyranoside; see Methods for details). In the three experiments shown a heme to protein molar ratio of 1.2 was used. The protein concentration in each experiment is shown. Note that activity increase is detected even with concentrations of 50 nm, indicating that binding is strong. (D) Plot of enzyme activity versus [heme]/[protein] ratio in solution for a protein concentration of 1 μM. Activity was determined after a 5 minutes incubation and the plot supports a 1:1 binding stoichiometry.

Heme has a tendency to associate in aqueous solution at neutral pH, a process that is reflected in a time-dependent decrease in the intensity of the Soret band, which becomes “flatter” upon formation of dimers and higher associations^30^. However, the process is reversed upon addition of the essentially heme-free ancestral glycosidase (Figure 5B), indicating that the protein binds heme and shifts the association equilibria towards the monomeric state. Remarkably, heme binding is also reflected in a several-fold increase in enzymatic activity which occurs on the seconds time scale when the heme and enzyme concentration are at a ~micromolar concentration (Figure 5C). Determination of activity after suitable incubation times for different heme:protein ratios in solution yielded a plot with an abrupt change of slope at a stoichiometric ratio of about 1:1 (Figure 5D). These experiments were carried out with ~micromolar heme and protein concentrations, indicating therefore a tight, sub-micromolar binding. Indeed, increases in activity upon heme addition to a protein solution were observed (Figure 5C) even with concentrations of ~50 nanomolar. The 1:1 stoichiometry of heme/protein was confirmed by experiments in which the protein was incubated with an excess of heme and free heme was removed through exclusion chromatography (2 passages through PD10 columns). The protein was then quantified by the bicinchoninic acid method^31^ with the Pierce™ BCA Protein Assay Kit while the amount of heme was determined using the pyridine hemochrome spectrum^32^ after transfer to concentrated sodium hydroxide (see Methods for details). This resulted in a heme/protein stoichiometry of 1.03±0.03 from five independent assays.

The experiments described above allowed us to set up a procedure for the preparation of the ancestral protein saturated with heme and to use this preparation for activity determinations and crystallization experiments. The procedure (see Methods for details) involved *in vitro* reconstitution using hemin but did not include any chemical system capable of performing a reduction. It is therefore safe to assume that our heme-bound ancestral glycosidase contains iron in the +3 oxidation state. Activity determinations with the heme-saturated ancestral enzyme corroborated that heme binding increases activity by ~3 fold (see Michaelis plots in Figure 4). X-ray crystallography confirmed the presence of one heme per protein molecule (Figures 6A, 6B and S7), which is located at the same site, with the same orientation and involved largely in the same molecular interactions in the three protein molecules (A, B, C) observed in the crystallographic unit cell. Besides interactions with several hydrophobic residues, the bound heme interacts (Figure 6A) with Tyr264 of α-helix 8, Tyr350 of α-helix 13, Arg345 of β-strand B and, directly via a water molecule, with Lys 261 of β-strand B, although this latter interaction is only observed in chain A. The bound heme shows B-factor values similar to those of the surrounding residues (Figure 6B), it is well-packed and 95% buried (Figure 6C). Indeed, the accessible surface area of the bound heme is only 43 Å^2^ (see Table S5 for atom detail), compared to the ~800 Å^2^ accessible surface area for a free heme^33^. The amino acid residues that interact with the heme include Tyr264 as the axial ligand while Arg345 and Tyr350 interact with one of the propionate moieties of the heme group (Figures 6A, 6B and S7). Interestingly, similar interactions are found is some modern heme proteins, including tyrosine as the axial ligand in some cases^33^.

**Figure 6.**
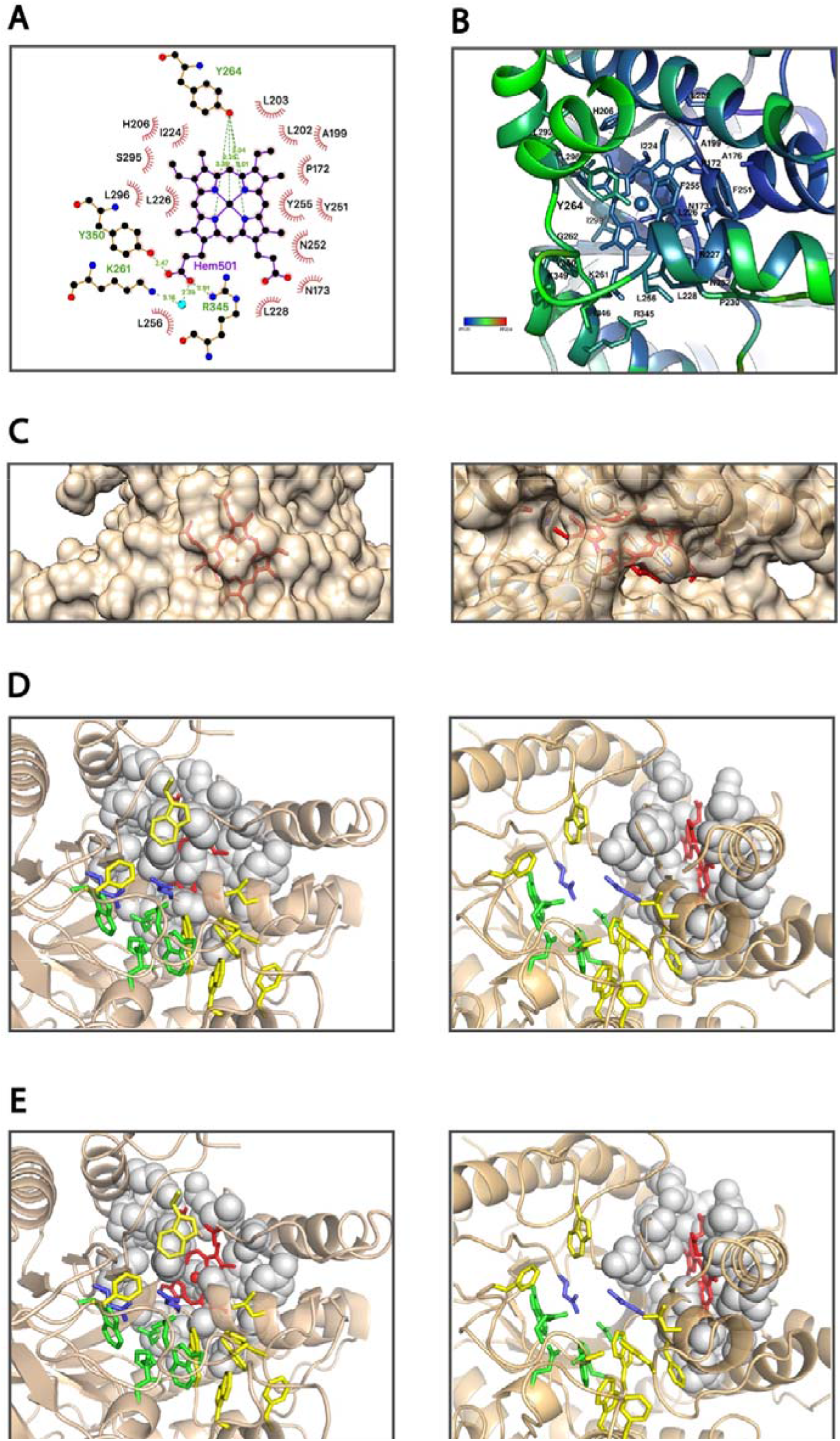
Local molecular environment of the bound heme in the ancestral protein. (A) Schematic representation of the heme molecule and the neighbor residues in the 3D-structure. (B) Heme group (red) and residues directly interacting with the heme colored by B value. (C) Van der Waals surface of the ancestral protein shown in translucent brown, so that it becomes visually apparent that the heme (shown in red) is mostly buried. (D) View of the active site of the ancestral protein showing the catalytic carboxylic acid residues (blue) and the residues involved in binding of the glycone (yellow) and aglycone (green) parts of the substrate molecule. The residues that block the connection of the heme group with the active site are shown with van der Waals spheres and colored in grey. (E) Same as in C, but the residues blocking the connection of the heme with the active site have been computationally mutated to alanine, in such a way that now the iron of the heme group can be seen at the bottom of the active site in the chosen view.

Heme binding clearly rigidifies the ancestral protein, as shown by fewer missing regions in the electronic density map, in contrast to the structure of the heme-free protein (see Figures 2A, 2B and 2C). This is also confirmed by molecular dynamics simulations of the ancestral glycosidase both with and without heme bound (Figure 3). That is, the MD simulations performed without the heme bound show that most of the protein has higher flexibility (Figure 3E, with ΔRMSF values greater than 0 in most of the sequence), particularly in the regions where the B-factors also indicate high flexibility (Figure 2B). This is noteworthy, as the only difference in starting structure between the two sets of simulations is the presence or absence of the heme, the starting structures are otherwise identical. The MD simulations show that removing the heme from the heme-bound structure has a clear effect on the flexibility of the whole enzyme, increasing it relative to the heme bound structure (Figure 3E), again also indicated by the B-factors (Figure 3). There are two regions where this difference is particularly pronounced. The first spans residues 252-265, which is located where the heme Fe(III) atom forms an interaction with the Tyr264 side chain as an axial ligand. Removing the heme removes this interaction, thus inducing greater flexibility in this region. The second region with increased flexibility spans 319-327, where again we observe that removing the heme increases the flexibility of this region.

Lastly, we note that the heme is located near the enzyme active site (at about 8 Å from the catalytic glutamate at position 171) but does not have direct access to this site as revealed in the structure (Figure 6D). Therefore, the increase in activity observed upon heme binding is an allosteric effect likely linked to dynamics (see Discussion) since heme binding does not substantially alter the position/conformation of the catalytic carboxylic acids nor the residues involved in substrate binding according to the static X-ray structures (Figure 2E).

## DISCUSSION

The TIM-barrel is the most common protein fold, accounting for ~10% of known enzyme structures and providing a scaffold for an enormous diversity of biomolecular functions^11–13^. It is composed of eight parallel (β/α) units linked by hydrogen bonds forming a cylindrical core (“the barrel”) with secondary structure elements connected by loops. The high capability of the fold to accommodate a wide diversity of different natural functions is likely linked to its modular architecture, with the barrel (and the αβ loops) providing stability and allowing a substantial degree of flexibility, variability and, therefore, evolvability for the βα loops. That is, the barrel provides a stable platform that can accommodate loops of different sequences and conformations at the so-called catalytic face.

Remarkably, the differences in conformational flexibility between different parts of the molecule appear to be even more pronounced in our ancestral TIM-barrel glycosidase. Stability is still guaranteed by a rigid barrel core, but flexibility is greatly enhanced and extends to large parts of the structure, as shown by a combination of computational and experimental results. Conformational flexibility implies that the protein in solution is sampling a diversity of conformations. On the one hand, this may prevent the enzyme from reaching the highest levels of catalysis for a given natural reaction since the protein ensemble may not be shifted towards the most active conformations. Indeed, while modern glycosidases approach catalysis levels up to 17 orders of magnitude above the rate of spontaneous glycoside bond hydrolysis^16^, the ancestral glycosidase displays turnover numbers about two orders of magnitude below the modern glycosidases studied here (Figure 4 and Table S3). On the other hand, flexibility is key to the emergence of new functions and contributes to evolvability, since minor conformations that catalyze alternative reactions may be enriched by subsequent evolution^34–37^. Therefore, the ancestral TIM-barrel described here holds promise as a scaffold for the generation of *de novo* catalysts, an important and largely unsolved problem in enzyme engineering. We have recently shown^38^ that completely new enzyme functions can be generated through a single mutation that generates both a cavity and a catalytic residue, provided that conformational flexibility around the mutation site allows for substrate and transition-state binding^10,36,37^. The combination of a rigid core that provides stability with high flexibility in specific regions makes the ancestral protein studied here an excellent scaffold to develop this minimalist approach to *de novo* catalysis (work in progress).

Catalytic features of the ancestral glycosidase, such as diminished activity levels and lack of specialization for glucopyranoside substrates, would seem consistent with an early stage in the evolution of family 1 glycosidases. It has been proposed that TIM-barrel proteins originated through fusions of smaller fragments^39^. The high conformational flexibility in some regions of the ancestral glycosidase structure would then also seem consistent with an early evolutionary stage, since fragment fusion is not expected to immediately lead to efficient packing and conformational rigidity in all parts of the generated structure. On the other hand, the capability of the reconstructed ancestral glycosidase to bind heme tightly and stoichiometrically at a well-defined site is rather surprising. None of the ~5500 X-ray structures for the ~1400 glycosidases currently reported in CAZy shows a porphyrin ring. It is certainly possible that heme binding to the ancestral glycosidase is simply an accidental byproduct of the high conformational flexibility at certain regions of the structure, although the tightness of the binding and the specificity of the molecular interactions involved argue against this possibility. In any case, this is an issue that can be investigated by studying modern glycosidases. If heme binding is a functional ancestral feature (a product of selection), we may expect that at least some modern glycosidases show some inefficient, vestigial capability to bind heme, in keeping with the general principle that features that become less functional undergo evolutionary degradation^40^. No mention of heme binding to modern family 1 glycosidases can be found in the CAZypedia resource^21^, but, of course, there is no reason why researchers in the glycosidase community should have tested heme-binding capabilities. Albeit limited, our results suggest that some modern family 1 glycosidases may retain a vestigial capability to bind heme. This suggests a complex evolutionary history for this family of enzymes involving perhaps a fortuitous (i.e., contingent) early fusion event with a heme-containing domain. One possibility in this context is that heme had a functional role in the isolated heme-containing domain, which was no longer required when the domain was fused with the larger glycosidase scaffold, thus enabling the subsequent degradation of the heme binding capability. Here, we focus on the relevant protein-engineering implications, which are independent of any evolutionary interpretations. In this context, heme-binding to the ancestral glycosidases opens up new possibilities that are briefly described below.

Metalloporphyrins are essential parts of many natural enzymes involved in redox and rearrangement catalysis and can be engineered for the catalysis of non-natural reactions^41^. Remarkably, however, the combination of the highly-evolvable TIM-barrel scaffold and the catalytically versatile metalloporphyrins is exceedingly rare among known modern proteins. A porphyrin ring is found in only 13 out of the 7637 PDB entries that are assigned the TIM-barrel fold according to the CATH classification^42^. These 13 entries correspond to just two proteins. One of them, uroporphyrinogen decarboxylase, is an enzyme involved in heme biosynthesis, while in the other identified case, flavocytochrome B2, the bound heme is far from the active site. By contrast, heme appears at about 8 Å from the catalytic Glu171 in our ancestral glycosidase. While its connection with the natural active site is blocked by several side chains in the determined structure (Figure 6D), it would appear feasible to use protein engineering to establish a conduit. As a simple illustration of this possibility, mutating Pro172, Asn173, Ile224, Leu226, Asn227 and Pro 272 to alanine *in silico* (Figure 6E) increases the accessible surface area of heme from barely 43 Å^2^ to about 300 Å^2^ and exposes the side of the heme facing the active site. The possibility thus arises that the engineering of metalloporphyrin catalysis, through rational design and/or laboratory evolution, would benefit from the evolutionary possibilities afforded by the flexible βα loops at a TIM-barrel catalytic face.

More immediate engineering possibilities arise from the allosteric modulation of catalysis in the ancestral protein, a phenomenon that, to our knowledge, has never been reported for modern glycosidases. Heme binding rigidifies the ancestral glycosidase and causes a several-fold activity enhancement. Heme is not expected to catalyze glycoside hydrolysis and, in any case, the bound heme does not have access to the active site in the experimentally determined ancestral structure. The activity enhancement upon heme binding is therefore an allosteric effect likely linked to dynamics, as it is the case with other allosteric effects reported in the literature^43^. Regardless of whether this feature is truly ancestral or just a byproduct of the enhanced conformational flexibility of the putative ancestral glycosidase, it is clear that it can provide a basis for biosensor engineering. For instance, computational design and laboratory directed evolution could be used to repurpose the heme binding site for the binding of a targeted substance of interest and to achieve a large concomitant change in glycosidase activity. The development of this application should be facilitated by the availability of a wide diversity of synthetic chemical probes for the sensitive detection of glycosidase activity^17^. In total, we anticipate that novel combinations of protein features will generate new possibilities in protein biotechnology and engineering.

## ACKNOWLEDGMENTS

This work was supported by Human Frontier Science Program Grant RGP0041 (J.M.S.-R., E.A.G., B.S. and S.C.L.K.), NIH grant R01AR069137 (E.A.G.), Department of Defense grant MURI W911NF-16-1-0372 (E.A.G.), the Swedish Research Council (2019-03499) (S.C.L.K.), the Knut and Alice Wallenberg Foundation (2018.0140 and 2019.0431) (S.C.L.K.), Spanish Ministry of Economy and Competitiveness/FEDER Funds Grants BIO2015-66426-R (J.M.S.-R.) RTI2018-097142-B-100 (J.M.S.-R.) and BIO2016-74875-P (J.A.G.). The simulations were enabled by resources provided by the Swedish National Infrastructure for Computing (SNIC) at UPPMAX partially funded by the Swedish Research Council through grant agreement no. 2016-07213. We acknowledge the Spanish Synchrotron Radiation Facility (ALBA, Barcelona) for provision of synchrotron radiation facilities and the staff at XALOC beam line for their invaluable support. We are also grateful to Victoria Longobardo Polanco (Proteomic Unit, Institute of Parasitology and Biomedicine “López-Neyra”) for help with mass spectrometry experiments and data analyses,

## METHODS

### Ancestral sequence reconstruction

#### Dataset construction

Characterized GH1 protein sequences were retrieved from the Carbohydrate-active enzyme database (CAZy)^18^, including β-glucosidase (accession numbers: ACI19973.1 and AAL80566.1), 6-P-β-glucosidase (AIY91871.1), β-mannosidase (AAL81332.1) and myrosinase (AAK32833.1). These characterized protein sequences were utilized as seeds to identify additional homologous sequences. GH1 homologs were retrieved for all three domains of life from GenBank (http://www.ncbi.nlm.nih.gov/) using BLASTp, with the cutoff threshold of <1×10^−5^. Sequences with the minimum length of 300 amino acids were included in the dataset. Taxonomically redundant sequences were excluded. A total number of 150 sequences were collected for further analysis.

#### Phylogenetic analysis

Sequences were aligned using T-Coffee. Initial non-bootstrapped phylogenetic trees were constructed using RAxML (ver. 8.2.11) to identify major clusters and to eliminate spurious sequences^44^. The RAxML analysis was performed with the hill-climbing mode using the gamma substitution model. These initial trees were used as starting point for more thorough Bayesian analysis. MrBayes (ver. 3.2.6) was conducted using the WAG amino acid replacement model with a gamma distribution and invariable sites model for at least 1,000,000 generations, with samplings at 100 intervals, and two runs with four chains per run in order to monitor convergence^45^. Twenty-five percent of sampled points were discarded as burnin. The tree topology was broadly identical between the RAxML and MrBayes analyses.

#### Ancestral sequence reconstruction

Ancestral sequences were reconstructed using FastML (ver. 3.1) with the WAG amino acid replacement model with a gamma distribution for variable replacement rates across sites^46^.

### Database searches

We searched the CAZy database in order to ascertain the presence of porphyrin rings in reported glycosidase structures. We systematically went through all 167 glycoside hydrolase families of the database in March 2020. For each family, we checked the structure section and we individually examined all the links provided to the protein data bank. Overall, we examined 5565 PDB files corresponding to 1435 different glycosidase enzymes. We did not find a single example of a reported structure with a bound porphyrin ring.

We also used the CAZy database as a starting point of an extensive literature search for optimum temperature values of family 1 glycosidases. We examined the section of characterized enzymes for family 1 glycosidases, the references included in the corresponding GenBank links, as well as the publications that cite those references in a Google Scholar search. Several hundred published articles were examined for experimental activity *versus* temperature profiles and reported values of the optimum temperature. We found such data for 127 different family 1 glycosidases. In many cases, the oligomerization state of the enzymes was also provided in the original references. The environmental temperatures (optimum growth temperatures) of the corresponding host organisms could be found in most cases in the “Bergey’s Manual of Systematic Bacteriology” although, in some cases, literature searches were performed to find the optimum temperatures. Most organisms were classified as hyperthermophilic, extreme thermophilic, thermophilic, mesophilic or psychrophilic in the Bergey’s manual or the relevant literature references. We have used this classification to color-code Figure 1C, since it leads to clear and intuitive data clusters. All the information related to the values of the optimum temperature for activity and the organismal living temperature is collected in Table S2.

In order to find examples of proteins with the TIM-barrel fold and a bound porphyrin ring in the reported structures, we checked (March-2020) all entries in the protein data bank that are classified as TIM-barrels in the CATH database. We examined a total of 7637 PDB files and found a porphyrin ring in only 13 of them. These 13 structures correspond to two proteins: flavocytochrome B2, a multi-domain protein in which the porphyrin ring is located in the non-TIM-barrel domain, and uroporphyrinogen decarboxylase, which is an enzyme involved in heme biosynthesis.

### Protein expression and purification

The different proteins studied in this work were purified following procedures we have previously described. Briefly, genes for the His-tagged proteins in a pET24 vector with kanamycin resistance were cloned into *E. coli* BL21 (DE3) cells, and the proteins were purified by Ni-NTA affinity chromatography in HEPES buffer. The His tag was placed at the C-terminus, i.e., at a position that is well removed from the catalytic face of the barrel, the regions of enhanced conformational flexibility in the ancestral protein and the heme binding site. Since the ancestral protein is susceptible to proteolysis, we included protease inhibitors in all steps of the purification (cOmplete^®^ EDTA-free Protease Inhibitor Cocktail from Roche, ref. 11873580001). Protein solutions were prepared by exhaustive dialysis against the desired buffer, typically 50 mM HEPES pH 7.

### Preparation of the ancestral protein with bound heme and determination of the heme to protein ratio

Stock solutions of hemin (heme with iron in the +3 oxidation state) were prepared daily in 1.4 M sodium hydroxide. Prior to use, the stock solution was diluted (typically 1:100) into HEPES buffer 50 mM, pH 7 and this solution was immediately used.

The ancestral protein with bound heme was prepared by incubating the protein with a 5-fold excess of heme for about one hour, followed by passage through a PD10 column and a Superdex-200 column to eliminate the non-bound heme. The heme to protein ratio in the resulting protein samples could be roughly estimated from the absorbance of the Soret band and the protein band at 280 nm in UV-VIS spectra. This procedure is not exact because the Soret band may depend on the interactions of the bound heme and, also, heme can show some absorption at 280 nm. For more accurate characterization protein concentration was determined by the bicinchoninic acid method^31^ with the Pierce™ BCA Protein Assay Kit (ThermoFisher Scientific). A method based on pyridine hemochrome spectra^32^ was used to determine the amount of heme. Briefly, 25 μl of 0.1 M potassium ferricyanide were added to a mixture of 2 mL or pyridine, 2 mL 0.1 M NaOH and 2 mL water. This solution was mixed with the protein solution in a 1:1 volume ratio and an excess of sodium dithionate was added. Lastly, the amount of heme was calculated from the absorbance of the pyridine hemochrome at 556 nm after correction for the absorbance of a blank. Using this approach, a heme to protein stoichiometry of 1.03±0.03 was determined from 5 independent measurements.

### Activity determinations

Glucosidase and galactosidase activities were tested following the absorbance of p-nitrophenol at 405 nm upon the hydrolysis of 4-nitrophenyl-β-D-glucopyranoside and 4-nitrophenyl-β-D-galactopyranoside^25^. Rates were calculated from the initial absorbance vs. time slope and the known extinction coefficient of p-nitrophenol at pH 7. Experiments at different substrate concentrations were carried out to arrive at Michaelis plots for the ancestral and several modern glycosidases studied in this work (Figure 4 and Figure S5).

As it is common in the literature, values of the optimum activity temperatures were determined from the profiles of activity versus temperature derived from measurements performed after several-minute incubations at each temperature^23^. Briefly, the protein was incubated at the desired temperature with 1 mM substrate in HEPES buffer 50 mM pH 7 and, after 10 minutes, the reaction was stopped by adding sodium carbonate to a concentration of 0.5 M. The amount of substrate hydrolyzed was determined from the absorbance of p-nitrophenol at 405 nm. We confirmed that the 10-minute incubation only hydrolyzed a fraction of the substrate present and, therefore, that the amount of substrate hydrolyzed after a 10-minute incubation is a suitable metric of enzyme activity. Profiles of activity versus temperature were determined using both 4-nitrophenyl-β-D-glucopyranoside and 4-nitrophenyl-β-D-galactopyranoside. The profiles for the ancestral glycosidase are shown in Figure 1B and those for the modern glycosidase from *Halothermothrix orenii* are given in Figure S4. In all cases, the profiles show a sharp maximum from which an unambiguous determination of the optimum temperature is possible. Note also that there is good agreement between the optimum temperature values derived using the two different substrates. Differential scanning calorimetry experiments were performed as we have previously described in detail^9^.

### Proteolysis experiments

For proteolysis experiments, the ancestral glycosidase and the modern glycosidase from *Halothermothrix orenii* at a concentration of 1 mg/mL were incubated at 25 °C with thermolysin for different times in HEPES buffer 50 mM pH 7 containing 10 mM calcium chloride. Stock solutions of thermolysin were prepared fresh in the same solvent at a concentration of 1 mg/mL and were diluted 1:10 when added to the protein solution. The reaction was stopped by addition of EDTA to a final concentration of 12.5 mM and aliquots were loaded into 15% (w/v) SDS-PAGE gels for electrophoresis. In some experiments, fragments separated by electrophoresis were extracted, desalted and subjected to LC-MS/MS analysis for mass determination. Fragment masses were determined by MALDI and their sequences were investigated using peptide mapping finger-printing and MALDI-TOF/TOF (Figure S2). This allowed us to locate approximately the cleavage sites, as shown in Figure 2C.

### Crystallization and structure determination

The ancestral glycosidase, dissolved in 150 mM NaCl, 50 mM HEPES pH 7.0, was concentrated to 35 mg/mL and to 70 mg/mL for the vapor-diffusion (VD) and counter-diffusion crystallization experiments, respectively. We checked by SDS electrophoresis that the concentrated protein used for crystallization was not proteolyzed. Hanging-drops VD experiments were prepared by mixing 1 μL of protein solution with the reservoir, in a 1:1 ratio, and equilibrated against 500 μL of each precipitant cocktail HR-I (Crystal Screen 1, Hampton Research). Capillary counter-diffusion experiments were set up in capillaries of 0.3 mm inner diameter using the CSK-24, AS-49 and PEG448-49 screening kits^47^. A similar procedure was followed for the crystallization of the ancestral glycosidase-heme complex, using two fixed concentrations at 75 and 30 mg/ml for the counter-diffusion and VD experiments. Experiments were performed at 293 K.

Crystals of the ancestral glycosidase were obtained only in condition #41 of HR-I, whilst the GH1N72-Heme complex crystallized in conditions #6 and #9 of HR-I and PPP8 of the mix of PEG counter-diffusion screen. Crystals were extracted either from the capillary or fished directly from the drop and subsequently cryo-protected by equilibration with 15 % (v/v) glycerol prepared in the mother liquid, flash-cooled in liquid nitrogen and stored until data collection. Crystals were diffracted at the XALOC beamline of the Spanish synchrotron light radiation source (ALBA, Barcelona). Indexed data were scaled and reduced using the CCP4 program suite^48^.

Initial data sets were obtained for the ancestral glycosidase crystals diffracting the X-ray to 2.5 Å. The clean (without water, ligands, etc.) 3D model of the β-glucosidase from *Thermotoga maritima* (PDB ID. 2J78) was used as search model for molecular replacement^48^. Two monomers were found in the asymmetric unit as expected from the Matthews coefficient for the P2(1) space group. Refinement, including Titration-Libration-Screw (TLS) parametrization, water pick and model validation was carried out with PHENIX suite^49^. Unidentifiable amino acids in the highly disordered region have been assigned as poly-UKN chains C and D corresponding to the 18 and 14 (poly-Alanine) of chains A and B, respectively.

Crystals of the ancestral glycosidase-heme complex belong to the same space group than the ancestral glycosidase but were not isomorphous. The determined unit cell was bigger accommodating three polypeptide chains in the asymmetric as determined from the Matthews coefficient. A similar protocol was followed to place the three monomers in the unit cell by molecular replacement and to refine the structure. After a first refinement round the presence of one protoporphyrin ring in each polypeptide chain was determined. It was also clear that disordered regions of the ancestral glycosidase model were visible in the heme complex model.

The summary of data collection, refinement statistics and quality indicators are collected in Table S6. The coordinates and the experimental structure factors have been deposited in the Protein Data Bank with ID 6Z1M and 6Z1H for the ancestral glycosidase with and without bound heme, respectively.

### Molecular dynamics simulations

Molecular simulations were performed on both ancestral glycosidases (this work, PDB ID: 6Z1M, 6Z1H) and the modern glycosidase from *Halothermothrix orennii* (PDB ID: 4PTV). The structure of the heme-free ancestral glycosidase was obtained by manually deleting the heme coordinates from the corresponding heme-bound crystal structure. The missing regions of the ancestral glycosidases were reconstructed using MODELLER. Histidine protonation states were selected based on empirical pK_a_ estimates performed using PROPKA 3.1 and visual inspection. All other residues were placed in their standard protonation states at physiological pH. The heme group was described using a bonded model, creating a bond between the Tyr264 side chain and the Fe(III) atom of the heme. We used MCPB.py as implemented in AMBER19^50^ to obtain the necessary parameters for creating the bonding pattern between the Fe(III) atom and the 4 nitrogen atoms of the heme and the tyrosine side chain oxygen. The resulting structure was then optimized, and frequency calculations were performed at the ωB97X-D/6-31G* level of theory followed by the Seminario method^51^ to obtain the force constants from the Hessian of the frequency calculation. Partial charges were obtained for the heme and for the Tyr264 side chain at the HF/6-31G* level of theory, using the restrained electrostatic potential (RESP) approach, following the MCPB.py protocol, and performing the calculations using Gaussian 09 Rev. E.01. All systems were solvated in an octahedral box filled with TIP3P water molecules^52^, with 10Å from the solute to the box edges in all directions. Na^+^ and Cl^−^ counter ions were added to the system to neutralize each enzyme. The protein was described using the AMBER ff14SB force field^53^, and the heme was described using the General AMBER Force Field (GAFF)^54^.

Following system preparation, the LEaP module of AMBER19 was used to generate the topology and coordinate files for the MD simulations, which were performed using the CUDA version of the PMEMD module of the AMBER19 simulation package. The solvated system was first subjected to a 5000 step steepest descent minimization, followed by a 5000 step conjugate gradient minimization with positional restraints on all heavy atoms of the solute, using a 5 kcal mol^−1^ Å^−2^ harmonic potential. The minimized system was then heated up to 300 K using the Berendsen thermostat^55^, with a time constant of 1 ps for the coupling, and 5 kcal mol^−1^ Å^−2^ positional restraints (again a harmonic potential) applied during the heating process. The positional restraints were then gradually decreased to 1 kcal mol^−1^ Å^−2^ over five 500 ps steps of NPT equilibration, using the Berendsen thermostat and barostat to keep the system at 300 K and 1 atm. For the production runs, each system was subjected to either 500 ns of sampling in an NPT ensemble at constant temperature (300 K) and constant pressure (1 atm), controlled by the Langevin thermostat, with a collision frequency of 2.0 ps^−1^, and the Berendsen barostat with a coupling constant of 1.0 ps. A 2 fs time step was used for all simulations, and snapshots were saved from the simulation every 5 ps. The SHAKE algorithm^56^ was applied to constrain all bonds involving hydrogen atoms. A 10 Å cutoff was applied to all nonbonded interactions, with the electrostatic interactions being treated with the particle mesh Ewald (PME) approach^57^. 10 independent simulations were performed for each starting structure during 500 ns (for RMSD convergence, see Figure S8). All the subsequent analyses were performed with the CPPTRAJ toolkit from Ambertools19^50^. Parameters used to describe the heme, input files, snapshots from our simulations, and simulation trajectories (with water molecules and ions removed to save files size) are available for download from Zenodo at DOI: 10.5281/zenodo.3857791.

