## Supporting Information for "Novel heme-binding enables allosteric modulation in an ancient TIM-barrel glycosidase"

^1^Departamento de Quimica Fisica. Facultad de Ciencias, Unidad de Excelencia de Quimica Aplicada a Biomedicina y Medioambiente (UEQ), Universidad de Granada, 18071 Granada, Spain.

^2^Department of Biology, Georgia State University, Atlanta, GA 30306 U.S.A.

^3^Science for Life Laboratory, Department of Chemistry-BMC, Uppsala University, BMC Box 576, S-751 23 Uppsala, Sweden.

^4^Department of Biochemistry, Molecular Biology, and Biophysics, University of Minnesota, Minneapolis, Minnesota, United States of America, & BioTechnology Institute, University of Minnesota, St. Paul, Minnesota, United States of America.

^5^Laboratorio de Estudios Cristalograficos, Instituto Andaluz de Ciencias de la Tierra, CSIC, Unidad de Excelencia de Quimica Aplicada a Biomedicina y Medioambiente (UEQ), Universidad de Granada, Avenida de las Palmeras 4, Granada 18100 Armilla, Spain.

^6^These authors contributed equally to this work

^7^Current address: Hit Discovery, Discovery Sciences, Biopharmaceutical R&D, AstraZeneca, 431 50 Gothenburg, Sweden.

*

**Supplementary Methods**

**Homology model preparation**

For the reconstructed ancestral sequence for node 72, 3D structures were initially prepared using homology modelling. In searching for the suitable templates, several criteria were imposed, i.e., high sequence identity (>= 50%) and query sequence coverage (>= 90%), as determined by BLAST and HHBlits (Altschul et al., 1997; Remmert et al., 2011), as well as the oligomerization state of a monomer and belonging to the GH1 family in the CAZY database. The following PDB structures fulfilled the criteria, and were used to create homology models with SWISS-MODEL (Arnold et al., 2006; Benkert et al., 2011; Biasini et al., 2014): family 1 β-glucosidase from Thermotoga maritima (1W3J), β-glucosidase A from Clostridium cellulovorans (3AHX), engineered β-glucosidase from soil metagenome (3CMJ), GH1 β-glucosidase Td2F2 (3WH5), and β-glucosidase 1A from Thermotoga neapolitana (5IDI). The quality of the five homology models was assessed using the Swiss-Model provided scores and the structural parameters estimated with MolProbity (Williams et al., 2018), and the models from 1W3J, 3CMJ, and 5IDI templates were selected for the further work. While the protein core (the (βα)_8_ barrel) was generally modelled very well, major differences could be observed between the homology models in the region of the catalytic loops.

Altschul SF, Madden TL, Schaäffer AA, Zhang J, Zhang Z, Miller W, Lipman DJ (1997) Gapped BLAST and PSI-BLAST: a new generation of protein database search programs. Nucleic Acid Res 25:3389-3402.

Arnold K, Bordoli L, Kopp J, Schwede T (2006) The SWISS-MODEL workspace: a web-based environment for protein structure homology modelling. Bioinformatics 22:195-201.

Remmert M, Biegert A, Hauser A, Söding J (2011) HHblits: lightning-fast iterative protein sequence searching by HMM-HMM alignment. Nat Methods 9:173-175.

Benkert P, Biasini M, Schwede T (2011) Toward the estimation of the absolute quality of individual protein structure models. Bioinformatics 27:343-350.

Biasini M, Bienert S, Waterhouse A, Arnold K, Studer G, Schimdt T, Kiefer F, Gallo Cassarino T, Bertoni M, Bordoli L, Schwede T (2014) SWISS-MODEL: modelling protein tertiary and quaternary structure using evolutionary information. Nucleic Acid Res 42:W252-W258.

Williams CJ, Headd JJ, Moriarty NW, Prisant MG, Videau LL, Deis LN, Verma V, Keedy DA, Hintze BJ, Chen VB, Jain S, Lewis SM, Arendall WB 3^rd^, Snoeyink J, Adams PD, Lovell SC, Richardson JS, Richardson DC (2018) MolProbity: More and better reference data for improved all-atom structure validation. Protein Sci 27:293-315.

**Supplementary Tables**

**Table S1.** Sequence identity of the ancestral glycosidase with the modern glycosidases in the set used as starting point for ancestral sequence reconstruction. The sequence identity values with the ancestral protein span the 0.26-0.59 range.

| Ancestral | 1,000 |
| --- | --- |
| Euk-XP_002676353.1.Naegleria.gruberi.NEG-M.Oth | 0,483 |
| Euk-OLQ05614.1.Symbiodinium.microadriaticum.SAR | 0,463 |
| Euk-OCF22674.1.Kwoniella.bestiolae.CBS.10118.Fg | 0,439 |
| Euk-KXS17974.1.Gonapodya.prolifera.JEL478.Fg.435 | 0,366 |
| Euk-KOO32978.1.Chrysochromulina.CCMP291.Oth.458 | 0,431 |
| Euk-KNC76787.1.Sphaeroforma.arctica.JP610.Met.475 | 0,389 |
| Euk-GAX29444.1.Fistulifera.solaris.SAR.446 | 0,519 |
| Euk-GAX16517.1.Fistulifera.solaris.SAR.450 | 0,500 |
| Euk-ETP15771.1.Phytophthora.parasitica.CJ01A1.SAR | 0,364 |
| Euk-ETP11111.1.Phytophthora.parasitica.CJ01A1.SAR | 0,516 |
| Euk-ETP02644.1.Phytophthora.parasitica.CJ01A1.SAR | 0,366 |
| Euk-EOD40853.1.Emiliania.huxleyi.CCMP1516.Oth.401 | 0,249 |
| Euk-EOD13795.1.Emiliania.huxleyi.CCMP1516.Oth.457 | 0,432 |
| Euk-EIE19776.1.Coccomyxa.subellipsoidea.C-169.Arpl | 0,271 |
| Euk-CHR-CAC34952.1.Piromyces.E2.Fg.463 | 0,393 |
| Euk-CHR-CAC08178.1.Homo.sapiens.Met.452 | 0,419 |
| Euk-CHR-BAO04178.1.Delphinium.grandiflorum.Arpl | 0,418 |
| Euk-CHR-BAE87009.1.Phanerochaete.chrysosporium.Fg | 0,464 |
| Euk-CHR-BAE63197.1.Aspergillus.oryzae.RIB40.Fg.460 | 0,440 |
| Euk-CHR-AGS32242.1.Coptotermes.gestroi.Met.460 | 0,424 |
| Euk-CHR-AAP13852.1.Bombyx.mori.Met.462 | 0,398 |
| Euk-CHR-AAL92115.1.Glycine.max.Arpl.456 | 0,377 |
| Euk-CHR-AAL37719.1.Solanum.lycopersicum.Arpl.461 | 0,435 |
| Euk-CHR-AAL25999.1.Brevicoryne.brassicae.Met.453 | 0,419 |
| Euk-CHR-AAK62412.1.Arabidopsis.thaliana.Arpl.465 | 0,404 |
| Euk-CHR-AAK32833.1.Arabidopsis.thaliana.Arpl.462 | 0,392 |
| Euk-CBJ30694.1.Ectocarpus.siliculosus.SAR.450 | 0,489 |
| Euk-AAG46031.1.Leishmania.infantum.Exc.456 | 0,371 |
| CPR-OYX53756.1.Sacchari.32-50-13.Oth.420 | 0,303 |
| CPR-OHA99561.1.Zambryski.R-O2_12_FULL_43_12b.Pc | 0,345 |
| CPR-OHA32072.1.Taylor.R-O2_01_FULL_45_15b.Pc | 0,340 |
| CPR-OHA13338.1.Taga.R-O2_01_FULL_39_11.Pc.405 | 0,317 |
| CPR-OHA09246.1.Sung.R-O2_01_FULL_59_16.Pc.399 | 0,311 |
| CPR-OGZ46118.1.Ryan.R-O2_01_FULL_48_27.Pc.416 | 0,330 |
| CPR-OGZ07021.1.Lloyd.R-O2_02_FULL_50_13.Pc.385 | 0,328 |
| CPR-OGZ06701.1.Lloyd.R-O2_02_FULL_50_11.Pc.387 | 0,332 |
| CPR-OGZ02593.1.Lipton.R-O2_01_FULL_53_13.Pc.396 | 0,315 |
| CPR-OGY99744.1.Lipton.R-O2_01_FULL_52_25.Pc.404 | 0,320 |
| CPR-OGY99381.1.Lipton.R-O2_01_FULL_52_25.Pc.410 | 0,305 |
| CPR-OGY71081.1.Jackson.R-O2_01_FULL_44_13.Pc.395 | 0,300 |
| CPR-OGY40781.1.Brenner.RIFOXYD1_FULL_41_16.Pc.389 | 0,312 |
| CPR-OGM99997.1.Yanofsky.R-O2_01_FULL_41_53.Pc.416 | 0,322 |
| CPR-OGM98994.1.Yanofsky.R-O2_01_FULL_41_26.Pc.386 | 0,314 |
| CPR-OGM90535.1.Wolfe.R-O2_01_FULL_38_11.Pc.412 | 0,336 |
| CPR-OGM00912.1.Uhr.RIFOXYC2_FULL_47_19.Pc.406 | 0,334 |
| CPR-OGL39085.1.Sacchari.R-O2_02_FULL_46_7.Oth | 0,311 |
| CPR-OGL30033.1.Sacchari.R-O2_02_FULL_47_12.Oth | 0,308 |
| CPR-OGK55604.1.Roizman.R-O2_01_FULL_45_11.Mc | 0,325 |
| CPR-OGJ70814.1.Peri.R-O2_12_FULL_53_10.Oth.390 | 0,318 |
| CPR-OGG92310.1.Kuenen.R-O2_12_FULL_42_13.Pc.406 | 0,316 |
| CPR-OGG87996.1.Kaiser.RIFOXYD1_FULL_42_15.Pc.382 | 0,312 |
| CPR-OGG55752.1.Kaiser.R-O2_01_FULL_55_37.Pc.387 | 0,317 |
| CPR-OGG06799.1.Gottesman.R-O2_01_FULL_42_12.Mc.399 | 0,326 |
| CPR-OGF90942.1.Giovannoni.R-O2_02_FULL_45_14.Pc | 0,351 |
| CPR-OGE98567.1.Doudna.R-O2_12_FULL_42_9.Pc.394 | 0,348 |
| CPR-OGE83632.1.Doudna.R-O2_02_FULL_43_13b.Pc.393 | 0,322 |
| CPR-OGE24950.1.Davies.R-O2_02_FULL_39_12.Mc.398 | 0,316 |
| CPR-OGE15085.1.Davies.GWA1_38_6.Mc.403 | 0,336 |
| CPR-KUK76511.1.WS6.34_10.Oth.396 | 0,338 |
| CPR-KKS47395.1.Giovannoni.GW2011_GWF2_42_19.Pc.397 | 0,351 |
| CPR-KKS25793.1.Jorgensen.GW2011_GWF2_41_8.Pc.408 | 0,332 |
| CPR-KKQ84990.1.Woese.GW2011_GWB1_38_8.Mc.399 | 0,347 |
| CPR-KKQ36955.1.Woese.GW2011_GWA1_37_7.Mc.396 | 0,353 |
| CPR-KKQ29214.1.Nomura.GW2011_GWA1_37_20.Pc.379 | 0,365 |
| CPR-KKQ12362.1.Moran.GW2011_GWF1_36_78.Pc.403 | 0,333 |
| Bac-WP_081838630.1.Thermogem.carboxidivorans.Clf | 0,511 |
| Bac-WP_053225736.1.Solirubrobacter.soli.Ac.440 | 0,483 |
| Bac-WP_033274371.1.Actinospica.acidiphila.Ac.393 | 0,313 |
| Bac-WP_028863797.1.Psychromonas.aquimarina.gP.448 | 0,415 |
| Bac-WP_026735568.1.Fischerella.PCC.9605.Cy.448 | 0,467 |
| Bac-WP_026389483.1.Acholeplasma.multilocale.Tn.438 | 0,336 |
| Bac-WP_016873414.1.Chlorogloeopsis.fritschii.Cy | 0,470 |
| Bac-WP_010471747.1.Acaryochloris.CCMEE.5410.Cy.441 | 0,488 |
| Bac-SEK57728.1.Rhodococcus.maanshanensis.Ac.387 | 0,300 |
| Bac-OUQ12786.1.Massiliomicrobiota.An142.Fm.454 | 0,449 |
| Bac-OUQ09514.1.Massiliomicrobiota.An142.Fm.466 | 0,340 |
| Bac-OIO58018.1.CG1_02_48_14.Mn.415 | 0,349 |
| Bac-OGS21660.1.RIFOXYA2_FULL_39_19.El.449 | 0,500 |
| Bac-OGF51021.1.RIFOXYA2_FULL_40_8.Frs.449 | 0,457 |
| Bac-OAA30738.1.Kosmotoga.arenicorallina.S304.Tg | 0,300 |
| Bac-KPM53585.1.Frankia.R43.Ac.388 | 0,322 |
| Bac-KIX85561.1.JCVI.TM6SC1.Dp.402 | 0,294 |
| Bac-GAT31492.1.Terrimicrobium.sacchariphilum.V.449 | 0,503 |
| Bac-EGF89970.1.Asticcacaulis.biprosthecum.C19.aP | 0,293 |
| Bac-EFH82146.1.Ktedonobacter.racemifer.Clf | 0,503 |
| Bac-CRX38655.1.Estrella.lausannensis.Chl.421 | 0,269 |
| Bac-CRH90323.1.Chlamydia.trachomatis.Chl.449 | 0,401 |
| Bac-CHR-CAC47470.1.Sinorhizobium.meliloti.1021.aP | 0,483 |
| Bac-CHR-BAC96154.1.Vibrio.vulnificus.YJ016.gP.445 | 0,481 |
| Bac-CHR-BAB37196.1.Escherichia.coli.Sakai.gP.464 | 0,366 |
| Bac-CHR-BAB36995.1.Escherichia.coli.Sakai.gP.462 | 0,346 |
| Bac-CHR-AIY95585.1.Bacillus.subtilis.Fm | 0,353 |
| Bac-CHR-AIY95329.1.Bacillus.subtilis.Fm | 0,365 |
| Bac-CHR-AIY91871.1.Bacillus.subtilis.Fm | 0,437 |
| Bac-CHR-AIY91623.1.Bacillus.subtilis.Fm | 0,473 |
| Bac-CHR-AGA60135.1.Microbacterium.Gsoil167.Ac.398 | 0,288 |
| Bac-CHR-AEI42200.1.Paenibacillus.mucilaginosus.Fm | 0,296 |
| Bac-CHR-ADD27066.1.Meiothermus.ruber.DSM.1279.DT | 0,524 |
| Bac-CHR-ADD01617.1.Thermoanaerobacter.italicus.Fm | 0,453 |
| Bac-CHR-ACV58907.1.Alicyclobaci.acidocaldarius.Fm | 0,551 |
| Bac-CHR-ACO44852.1.Deinococcus.deserti.VCD115.DT | 0,497 |
| Bac-CHR-ACM06095.1.Thermomicrobium.roseum.Clf | 0,521 |
| Bac-CHR-ACL70277.1.Halothermothrix.orenii.H.168.Fm | 0,557 |
| Bac-CHR-ACI19973.1.Dictyoglomus.thermophilum.Dg | 0,581 |
| Bac-CHR-ABU56651.1.Roseiflexus.castenholzii.Clf | 0,541 |
| Bac-CHR-ABR73190.1.Marinomonas.MWYL1.gP.444 | 0,449 |
| Bac-CHR-ABJ60960.1.Lactobacillus.gasseri.Fm | 0,340 |
| Bac-CHR-ABJ59900.1.Lactobacillus.gasseri.Fm | 0,494 |
| Bac-CHR-ABJ59596.1.Lactobacillus.gasseri.Fm | 0,362 |
| Bac-CHR-ABD82858.1.Saccharophagus.degradans.gP | 0,497 |
| Bac-CHR-ABD80656.1.Saccharophagus.degradans.gP | 0,443 |
| Bac-CHR-AAN58797.1.Streptococcus.mutans.UA159.Fm | 0,346 |
| Bac-CHR-AAK78365.1.Clostridium.acetobutylicum.Fm | 0,509 |
| Bac-CHR-AAG59862.1.Sphingomonas.paucimobilis.aP | 0,261 |
| Bac-CHR-AAB49339.1.Fusobacterium.mortiferum.Fs.442 | 0,525 |
| Bac-CHR-AAA24815.1.Dickeya.chrysanthemi.gP.456 | 0,365 |
| Bac-CCB88561.1.Simkania.negevensis.Z.Chl.422 | 0,270 |
| Bac-CAB95278.1.Streptomyces.coelicolor.A3_2.Ac.444 | 0,526 |
| Bac-BAY28878.1.Nostoc.carneum.NIES-2107.Cy.452 | 0,471 |
| Bac-BAM04503.1.Phycisphaera.mikurensis.Pl | 0,464 |
| Bac-BAL98072.1.Caldilinea.aerophila.DSM.14535.Clf | 0,522 |
| Bac-APF18099.1.Caldithrix.abyssi.DSM.13497.Cld.414 | 0,313 |
| Bac-AFG36462.1.Spirochaeta.africana.DSM.8902.Sp | 0,331 |
| Bac-AEP25088.1.Thermotoga.maritima.MSB8.Tg.443 | 0,593 |
| Bac-ADD01635.1.Thermoanaerobacter.italicus.Ab9.Fm | 0,578 |
| Bac-ADB52696.1.Conexibacter.woesei.DSM.14684.Ac | 0,496 |
| Bac-ACZ10042.1.Sebaldella.termitidis.ATCC.33386.Fs | 0,390 |
| Bac-ACZ09962.1.Sebaldella.termitidis.ATCC.33386.Fs | 0,399 |
| Bac-ACL69240.1.Halothermothrix.orenii.H.168.Fm.417 | 0,339 |
| Bac-ACI21065.1.Thermodesulfovib.yellowstonii.Nsr | 0,336 |
| Bac-ABJ83756.1.Solibacter.usitatus.Ellin6076.Ad | 0,305 |
| Bac-ABG04991.1.Rubrobacter.xylanophilus.Ac | 0,524 |
| Bac-AAK79373.1.Clostridium.acetobutylicum.Fm | 0,359 |
| Arc-SMD30897.1.Picrophilus.oshimae.Ery-Thp | 0,256 |
| Arc-Seed-BAA29440.1.Pyrococcus.horikoshii.Ery-Thc | 0,377 |
| Arc-OYT35865.1.Archaeoglobales.ex4484_92.Ery-Ach | 0,334 |
| Arc-OWP55065.1.Cuniculiplasma.C_DKE.Ery-Thp.470 | 0,263 |
| Arc-KYH41278.1.Bathyarchaeota.B26-2.Ery-Unc.477 | 0,278 |
| Arc-KJE49247.1.Acidiplasma.MBA-1.Ery-Thp.464 | 0,237 |
| Arc-CHR-ADL19795.1.Acidilo.saccharovorans.TK-Thp | 0,311 |
| Arc-CHR-ABW01492.1.Caldivir.maquilingensis.TK-Thp | 0,295 |
| Arc-CHR-AAL81332.1.Pyrococcus.furiosus.Ery-Thc | 0,294 |
| Arc-CHR-AAL80566.1.Pyrococcus.furiosus.Ery-Thc | 0,392 |
| Arc-CHR-AAL80197.1.Pyrococcus.furiosus.Ery-Thc | 0,275 |
| Arc-CHR-AAK43121.1.Sulfolobus.solfataricus.TK-Thp | 0,290 |
| Arc-CHR-AAD43138.1.Thermosphaera.aggregans.TK-Thp | 0,272 |
| Arc-BAB59827.1.Thermoplasma.volcanium.GSS1.Ery-Thp | 0,265 |
| Arc-AJB42198.1.Thermofilum.carboxyditrophus.TK-Thp | 0,285 |
| Arc-AJB41496.1.Thermofilum.carboxyditrophus.TK-Thp | 0,277 |
| Arc-AIF16120.1.marine.thaumarchaeote.TK-Thm | 0,345 |

**Table S2.** Literature values of the optimum activity temperatures and host living temperatures for modern family 1 glycosidases. The relevant references are appended at the end of the table. See Methods in the main text for details on the performed literature search. Organismal living temperature is defined as the optimum growth temperature. For some organisms, optimum intervals or lving temperature intervals are reported in the literature. Likewise, for some enzymes an interval of optimum activity temperature is provided in the literature. Organisms are classified as hyperthermophiles, extreme thermophiles, thermophiles, mesophiles and psycrophiles according to the descriptions reported in the literature. No classification is reported when such a description is not specifically provided in the literature, although most of the non-classified organisms are obviously mesophiles.

| Domain of life | **Enzyme description** | **Species** | **Organismal living temperature (ºC)** | **Enzyme optimum temperature (ºC)** | **Association state** | **Association state determination** | **Classification by growth temperature** | **Relevant**  **references** |
| --- | --- | --- | --- | --- | --- | --- | --- | --- |
| Archaea | β-glycosidase  (Tkβgly) | *Thermococcus kodakarensis KOD1* | 85 | 100 | Dimer | Native PAGE | Hyperthermophile | (Hwa et al., 2014)(Hwa et al., 2015)  (Ezaki et al., 1999)  (Atomi et al., 2004) |
| Archaea | β-mannosidase | *Pyrococcus furiosus* | 100 | 105 | Tetramer | Treatment with a cross-linker (dimethyl suberimidate) | Hyperthermophile | (Bauer et al., 1996)  (Fiala and Stetter, 1986)  (Robb et al., 2001) |
| Archaea | β-glucosidase | *Pyrococcus furiosus* | 100 | 102 - 105 | Tetramer | Native PAGE | Hyperthermophile | (Kengen et al., 1993)  (Fiala and Stetter, 1986)  (Robb et al., 2001) |
| Archaea | β-glycosidase  (rSSG) | *Sulfolobus shibatae* | 81 | 95 | Tetramer | Gel filtration chromatography | Hyperthermophile | (Park et al., 2007)  (Huber and Stetter, 2015) |
| Archaea | β-glucosidase  (Tpa-glu) | *Thermococcus pacificus P-4* | 80 - 88 | 75 | Not reported |  | Hyperthermophile | (Kim et al., 2015)  (Kobayashi, 2015) |
| Archaea | β-galactosidase | *Pyrococcus* *woesei* | 100 - 103 | 90 - 93 | Not reported |  | Hyperthermophile | (Da̧browski et al., 1998)  (Da̧browski et al., 2000)  (Stetter and Huber, 2015) |
| Archaea | β‐glycosidase (BGPh) | *Pyrococcus horikoshii* | 98 | Over 100 | Not reported |  | Hyperthermophile | (Matsui et al., 2000)  (Stetter and Huber, 2015) |
| Archaea | β-galactosidase (LacS) | *Sulfolobus solfataricus P2* | 85 | 90 | Not reported |  | Hyperthermophile | (Wu et al., 2013)  (Huber and Stetter, 2015a)  (Aguilar et al., 1997) |
| Archaea | β -glycosidase | *Acidilobus saccharovorans 345-15^T^* | 80 - 85 | 93 | Not reported |  | Hyperthermophilic | (Gumerov et al., 2015)  (Prokofeva et al., 2009) |
| Archaea | Glycosidehydrolase (CMbg0408) | *Caldivirga maquilingensis IC-167* | 85 | 110 | Not reported |  | Hyperthermophile | (Letsididi et al., 2017)  (Itoh et al., 2015) |
| Bacteria | β-glucosidase (BglB) | *Microbispora bispora NRRL 15568* | 50 - 65 | 60 | Monomer | Native PAGE and zymogram | Thermophile | (Wright et al., 1992)  (Kim, 2015) |
| Bacteria | β-D-glucosidase A (BglA) | *Bacillus sp. GL1* | 25 - 40 | 45 | Monomer | Native PAGE | Mesophile | (Hashimoto et al., 1998)  (Logan and Vos, 2015) |
| Bacteria | Phospho-β-glucosidase | *Fusobacterium mortiferum* | 35 - 37 | 35 - 40 | Monomer | Gel filtration chromatography | Mesophile | (Thompson et al., 1997)  (Olsen, 2014) |
| Bacteria | β-glucosidase (BglU) | *Micrococcus antarcticus* | 16.8 | 25 | Monomer | Gel filtration chromatography, native PAGE and zymogram | Psychrophile | (Fan et al., 2011)  (Busse, 2015)  (Liu et al., 2000) |
| Bacteria | β-D-glucosidase | *Paenibacillus sp. strain HC1* | 28 - 40 | 37 | Monomer | Gel filtration chromatography |  | (Harada et al., 2005)  (Fergus G. Priest, 2015) |
| Bacteria | β-glucosidase (bgl3) | *Streptomyces sp. QM-B814* | 25 - 35 | 50 | Monomer | Gel filtration chromatography | Mesophile | (Perez‐Pons et al., 1994)  (Kämpfer, 2015) |
| Bacteria | β-glycosidase (Ttbgly) | *Thermus thermophilus*  *HB 27* | 70 | 88 | Monomer | Gel filtration chromatography | Extreme thermophile | (Dion et al., 1999)  (da Costa et al., 2015)  (Henne et al., 2004) |
| Bacteria | β-glycosidase (KNOUC202βgly) | *Thermus thermophilus KNOUC202* | 70 | 90 | Monomer | Native PAGE and zymogram | Extreme thermophile | (Nam et al., 2010)  (da Costa et al., 2015)  (Henne et al., 2004) |
| Bacteria | β-glycosidase (TtβGly) | *Thermus thermophilus HJ6* | 70 | 90 | Monomer | Gel filtration chromatography | Extreme thermophile | (Gu et al., 2009)  (da Costa et al., 2015)  (Henne et al., 2004) |
| Bacteria | β-Glycosidase (SdBgl1B) | *Saccharophagus degradans strain 2–40* | 30 | 50 | Monomer | DLS and SAXS measurements  Gel filtration chromatography (this work) |  | (Brognaro et al., 2016)  (Ekborg et al., 2005) |
| Bacteria | β-glycosidase | *Thermus nonproteolyticus HG102* | 65 | 90 | Monomer | Gel filtration chromatography | Thermophile | (Xiangyuan et al., 2001)  (Wang et al., 2003) |
| Bacteria | β-glycosidases (bglA) | *Thermus*  *sp. IB-21* | 70 - 75 | 70 | Monomer | Native PAGE and zymogram | Extreme thermophile | (Kang et al., 2005)  (Kristjansson et al., 1986) |
| Bacteria | β-glycosidases (bglB) | *Thermus*  *sp. IB-21* | 70 - 75 | 80 | Monomer | Native PAGE and zymogram | Extreme  thermophile | (Kang et al., 2005)  (Kristjansson et al., 1986) |
| Bacteria | β-glycosidase (Tat β-gly) | [*Thermus flavus  AT-62*](https://www.sciencedirect.com/topics/biochemistry-genetics-and-molecular-biology/thermus) | 70 | 80 - 90 | Monomer | Native PAGE | Extreme  thermophile | (Sang et al., 2005)  (SAIKI et al., 1972) |
| Bacteria | β-glucan glucohydrolase (GghA) | *Thermotoga neapolitana* | 80 | 95 | Monomer | Gel filtration chromatography | Hyperthermophile | (Yernool et al., 2000)  (Nesbø et al., 2015) |
| Bacteria | β-glycosidase (tfi β-gly) | *Thermus filiformis*  *Wai33 A1* | 70 | 80 - 90 | Monomer | Native PAGE and zymogram | Thermophile | (Kang et al., 2004)  (da Costa et al., 2015)  (Mandelli et al., 2017) |
| Bacteria | β-glucosidase (HoBGLA) | *Halothermothrix orenii* | 60 | 65 -70 | Monomer | Gel filtration chromatography (this work) | Thermophile | (Hassan et al., 2015)  (Cayol et al., 2015) |
| Bacteria | β-1,4 glucosidase (*Ht*Bgl) | *Hungateiclostridium thermocellum* | 50 - 68 | 65 | Monomer | SAXS analysis | Thermophile | (Sharma et al., 2019)  (Akinosho et al., 2014) |
| Bacteria | β-glucosidase | *Sinorhizobium meliloti* | 25 - 30 | 45 | Monomer | Gel filtration chromatography, native PAGE and zymogram | Mesophile | (Kim et al., 2010)  (Kuykendall et al., 2015)  (Galardini et al., 2013) |
| Bacteria | Glycoside hydrolase (CoGH1A) | *Caldicellulosiruptor owensensis* | 75 | 75 - 85 | Monomer and trimer | Gel filtration chromatography | Extreme   thermophile | (Peng et al., 2016)  (Huang et al., 1998) |
| Bacteria | β-glucan glucohydrolase (BglA) | *Clostridium cellulovorans* | 37 | 50 | Dimer | Native PAGE and zymogram | Mesophile | (Kosugi et al., 2006)  (Rainey et al., 2015)  (Tamaru et al., 2011) |
| Bacteria | β-glucosidase (DtGH) | *Dictyoglomus thermophilum* | 73 | 90 | Dimer | Gel filtration chromatography | Extreme thermophile | (Zou et al., 2012)  (Patel, 2015)  (Ding et al., 2000) |
| Bacteria | β-glucosidase (Tm-BglA) | *Thermatoga maritima* | 80 | 90 | Dimer | Gel filtration chromatography (this work) | Hyperthermophile | (Song et al., 2011)(Xue et al., 2015)  (Nesbø et al., 2015) |
| Bacteria | β-glucosidase | *Caldicellulosiruptor saccharolyticus* | 70 | 70 | Dimer | Gel filtration chromatography | Thermophilic | (Hong et al., 2009) |
| Bacteria | β-galactosidase from (CcGH1) | *Clostridium cellulolyticum*  *H10* | 32 - 35 | 60 | Dimer and tetramer | Native PAGE and zymogram | Mesophile | (Liu et al., 2010)  (Rainey et al., 2015)  (Petitdemange et al., 1984) |
| Bacteria | β-glucosidase (reBglM1) | *Marinomonas MWYL1* | 20-40 | 40 | Trimer | Gel filtration chromatography (this work) |  | (Zhao et al., 2012)(John G. Holt et al., 1994) |
| Bacteria | β-glucosidase (DturβGlu) | *Dictyoglomus turgidum* | 72 | 80 | Tetramer | Gel filtration chromatography | Hyperthermophile | (Fusco et al., 2018)  (Patel, 2015)  (Fusco et al., 2018) |
| Bacteria | β-glucosidase (EaBglA) | *Exiguobacterium antarcticum B7* | -6 - 40 | 30 | Tetramer | Gel filtration chromatography | Psychrophilic | (Zanphorlin et al., 2016)  (Vishnivetskaya et al., 2007)  (Yoo et al., 2019) |
| Bacteria | β-glucosidase | *Sphingopyxis alaskensis* | ≈ 37 | 50 | Tetramer | Gel filtration chromatography |  | (Shin and Oh, 2014)  (Yabuuchi and Kosako, 2015) |
| Bacteria | β-glycosidase (Aaβ-gly) | *Alicyclobacillus acidocaldarius* | 60 - 65 | 85 | Octamer | Gel filtration chromatography | Thermophile | (Lauro et al., 2006)  (Darland and Brock, 1971) |
| Bacteria | β -glucosidase (CglT) | *Thermoanaerobacter brockii* | 65 - 70 | 75 | Not reported |  | Thermophile | (Breves et al., 1997)  (Onyenwoke and Wiegel, 2015a) |
| Bacteria | β-Glycoside Hydrolases (CfBgl1) | *Cellulomonas fimi* | ≈30 | 40 | Not reported |  | Mesophile | (Gao and Wakarchuk, 2014)  (Stackebrandt and Schumann, 2015)  (Clarke et al., 1996) |
| Bacteria | β-glucosidase from (FiBgl1A) | *Fervidobacterium islandicum* | 65 | 90 | Not reported |  | Extreme thermophile | (Jabbour et al., 2012)  (Huber and Stetter, 2015b)  (Huber et al., 1990) |
| Bacteria | β -glucosidase (Bgl1) | *Sphingomonas paucimobilis* | 25 - 30 | 50 | Not reported |  | Mesophile | (Marques et al., 2003)  (Yabuuchi and Kosako, 2015)  (Balkwill D.L., J.K. Fredrickson, 2006) |
| Bacteria | β-glucosidase (*bglSp*) | *Sphingomonas*  *sp. 2F2* | 25 - 30 | 37 | Not reported |  | Mesophile | (Wang et al., 2011)  (Yabuuchi and Kosako, 2015)  (Balkwill D.L., J.K. Fredrickson, 2006) |
| Bacteria | β-glucosidase (*bglX*) | *Lactococcus sp. FSJ4* | 28 - 30 | 40 | Not reported |  | Mesophile | (Fang et al., 2014)  (Teuber, 2015) |
| Bacteria | β-glucosidase from (BglPm) | *Paenibacillus*  *mucilaginosus*  *KCTC 3870T* | 28 - 40 | 45 | Not reported |  |  | (Cui et al., 2014)  (Fergus G. Priest, 2015) |
| Bacteria | β-glucosidase (AsBG1) | *Alicyclobacillus*  *sp. A4* | 60 | 55 | Not reported |  | Thermophile | (Cao et al., 2018)  (Bai et al., 2010)  (Gordon, 2017) |
| Bacteria | β-glucosidase (Te-BglA) | *Thermoanaerobacter ethanolicus*  *JW200* | 69 | 80 | Not reported |  | Extreme thermophile | (Song et al., 2011)  (Onyenwoke and Wiegel, 2015a)  (Wiegel and Ljungdahl, 1981) |
| Bacteria | β-glucosidase (Bglp) | *Anoxybacillus flavithermus subsp. yunnanensis E13T* | 60 | 60 | Not reported |  | Thermophile | (Liu et al., 2017)  (Pikuta, 2015) |
| Bacteria | β-Glucosidase (Bgl) | *Bacillus circulans subsp. Alkalophilus* | 30 - 37 | 50 - 55 | Not reported |  | Mesophile | (Paavilainen et al., 1993)  (Logan and Vos, 2015) |
| Bacteria | β-glucosidase (Bhbgl) | *Bacillus halodurans*  *C-125* | 15-55 | 50 | Not reported |  | Mesophile | (Xu et al., 2011)  (Logan and Vos, 2015)  (Lee et al., 2005) |
| Bacteria | β-glucosidase (bgl) | *Thermoanaerobacterium thermosaccharolyticum DSM 571* | 55 - 62 | 70 | Not reported |  | Thermophile | (Pei et al., 2012)  (Onyenwoke and Wiegel, 2015b) |
| Bacteria | β-glucosidase (BglC) | *Thermobifida fusca* | 35 - 53 | 50 | Not reported |  | Thermophile | (Spiridonov and Wilson, 2001)  (Trujillo and Goodfellow, 2015) |
| Bacteria | β-1,4-glucosidase (*Bgl*A) | *Thermotoga petrophila RKU-1* | 80 | 80 - 90 | Not reported |  | Hyperthermophile | (Haq et al., 2012)  (Nesbø et al., 2015) |
| Bacteria | β-glucosidase (DT-Bgl) | *Anoxybacillus*  *sp. DT3-1* | 55 | 70 | Not reported |  | Thermophile | (Chan et al., 2016)  (Chai et al., 2012) |
| Bacteria | β-glucosidase (BGL) | *Thermoanaerobacterium aotearoense P8G3#4* | 60 - 63 | 60 | Not reported |  | Thermophile | (Yang et al., 2015)  (Onyenwoke and Wiegel, 2015b) |
| Bacteria | β-glucosidase (Tt-bgl) | *Thermotoga thermarum*  *DSM 5069T* | 70 | 90 | Not reported |  | Hyperthermophile | (Zhao et al., 2013a)  (Windberger et al., 1989) |
| Bacteria | β-glucosidase (BglA) | *Exiguobacterium sp. DAU5* | 27 - 30 | 45 | Not reported |  |  | (Chang et al., 2011)  (Chang et al., 2013) |
| Bacteria | β-glucosidase (bglH) | *Bacillus licheniformis* | 49 | 50 | Not reported |  | Mesophile | (Zahoor et al., 2011)  (Lavermicocca et al., 2016) |
| Bacteria | β-Glucosidase (SC1059) | *Streptomyces Coelicolor A3 (2)* | 25 - 35 | 35 | Not reported |  | Mesophile | (Gu et al., 2013)  (Kämpfer, 2015)  (Ge et al., 2010) |
| Bacteria | β-glucosidase (SC7558) | *Streptomyces Coelicolor A3 (2)* | 25 - 35 | 35 | Not reported |  | Mesophile | (Gu et al., 2013)  (Kämpfer, 2015)  (Ge et al., 2010) |
| Bacteria | 6-phospho-β-glucosidase (Asb) | *Pectobacterium carotovorum subsp. carotovorum LY34* | 27 - 30 | 40 | Not reported |  |  | (An et al., 2005)  (Hauben et al., 2015) |
| Bacteria | β-glucosidase (CasB) | *Pectobacterium carotovorum subsp. carotovorum LY34* | 27 - 30 | 40 | Not reported |  |  | (Kim et al., 2013)  (Hauben et al., 2015) |
| Bacteria | β-glucosidase (BglY) | *Pectobacterium carotovorum subsp. carotovorum LY34* | 27 - 30 | 40 | Not reported |  |  | (An et al., 2012)  (Hauben et al., 2015) |
| Bacteria | β-glucosidase (celG) | *Pectobacterium carotovorum subsp. carotovorum LY34* | 27 - 30 | 40 | Not reported |  |  | (Hong et al., 2007)  (Hauben et al., 2015) |
| Bacteria | 6-phospho-β-glucosidase (Ascb) | *Pectobacterium carotovorum subsp. carotovorum LY34* | 27 - 30 | 40 | Not reported |  |  | (An et al., 2005)  (Hauben et al., 2015) |
| Bacteria | β-glucosidase (AcBg) | *Acidothermus cellulolyticus 11B* | 55 | 70 | Not reported |  | Thermophile | (Li et al., 2018)  (Barabote et al., 2009) |
| Bacteria | β-glucosidase (H0HC94) | *Agrobacterium tumefaciens 5A* | 25 -28 | 52 | Not reported |  | Mesophile | (Goswami et al., 2016)  (Goswami et al., 2017)  (Young et al., 2015) |
| Bacteria | β-glucosidase (BglZ) | *Bacillus amyloliquefaciens ABBD* | 30 - 40 | 25 | Not reported |  | Mesophile | (Kurniasih et al., 2014)  (Priest et al., 1987)  (Logan and Vos, 2015) |
| Bacteria | β-glucosidases  (BglC) | *Bacillus sp. SJ-10* | 30 - 40 | 40 | Not reported |  | Mesophile | (Lee et al., 2015)  (Jang et al., 2018)  (Logan and Vos, 2015) |
| Bacteria | β-glucosidase (*Bgl*B) | *Bacillus subtilis RA10* | 28 - 30 | 50 | Not reported |  | Mesophile | (Tiwari et al., 2017)  (Logan and Vos, 2015) |
| Bacteria | β-glucosidase (CbBgl1A) | *Caldicellulosiruptor bescii* | 72 - 75 | 85 | Not reported |  | Extreme thermophilic | (Bai et al., 2013)  (Yang et al., 2010) |
| Bacteria | β-galactosidase (GkGal1A) | *Geobacillus kaustophilus HTA426* | 60 | 70 | Not reported |  | Thermophile | (Shanshan et al., 2014)  (Takami et al., 2004) |
| Bacteria | 6-phospho-β-galactosidase (LacG) | *Lactobacillus casei strain BL23* | 30 - 40 | 41 | Not reported |  | Mesophile | (Bidart et al., 2018)  (Hammes and Hertel, 2015) |
| Bacteria | Phospho-β-galactosidase (LacG1) | *Lactobacillus gasseri* | 30 - 40 | 40 | Not reported |  | Mesophile | (Honda et al., 2012)  (Hammes and Hertel, 2015) |
| Bacteria | Phospho-β-galactosidase (rLacG2) | *Lactobacillus gasseri* | 30 - 40 | 50 | Not reported |  | Mesophile | (Honda et al., 2012)  (Hammes and Hertel, 2015) |
| Bacteria | Phospho-β-glucosidase (CelD) | *Oenococcus oeni* | 22 | 40 | Not reported |  |  | (Capaldo et al., 2011)  (Dicks and Holzapfel, 2015) |
| Bacteria | β-glucosidase (bglsG) | *Streptomyces griseus subsp. griseus* | 25 - 35 | 69 | Not reported |  | Mesophile | (Kumar et al., 2017)  (Kämpfer et al., 2014)  (Lacey, 1978) |
| Eukaryota | Β-glucosidase (hCBG) | [*Homo sapiens*](http://www.cazy.org/e356.html) | 37 | 50 | Monomer | Native PAGE and gel filtration chromatography | Mesophile | (Daniels et al., 1981)  (Berrin et al., 2002)  (Victor E. Del Bene., 1990)  (De Farias and Bonato, 2002) |
| Eukaryota | β-glucosidase (MbmgBG1) | *Macrotermes barneyi* | 20 - 28 | 50 | Monomer | Native PAGE and zymogram |  | (Wu et al., 2012)  (Wang Z. Y. et al., 2009) |
| Eukaryota | Β-glucosidase (bgl4) | *Humicola grisea var. thermoidea IF09854* | 38 - 46 | 55 | Monomer | Gel filtration chromatography | Thermophile | (Takashima et al., 1996)  (Takashima et al., 1999)  (Madan and Thind, 1998) |
| Eukaryota | β-glucosidase from (rThBgl) | *Trichoderma harzianum* | 28 - 32 | 40 | Monomer | Gel filtration chromatography | Mesophile | (Santos et al., 2016)  (Radhakrishna et al., 1988) |
| Eukaryota | β-glucosidase | *Aspergillus oryzae* | 30 | 50 | Monomer | Gel filtration chromatography |  | (Riou et al., 1998)  (Sakurai et al. 2014) |
| Eukaryota | β-glucosidase (OsTAGG1) | Oryza sativa *Japonica Group* | 25 - 30 | 50 | Monomer | Gel filtration chromatography |  | (Wakuta et al. 2010)  (Fageria et al., 2011) |
| Eukaryota | β-glucosidase (Os3BGlu6) | Rice *(Oryza sativa)* | 25 - 30 | 40 – 55 | Monomer | Crystal structure | Mesophile | (Seshadri et al., 2009)  (Fageria et al., 2011)  (Banerjee et al., 2019) |
| Eukaryota | β-glucosidase (BGQ60) | Seeds of barley *(Hordeum vulgare L.)* | 18 | 60 | Monomer | Gel filtration chromatography |  | (Leah et al., 1995)  (Fageria et al., 2011) |
| Eukaryota | β-glucosidase (Os3bglu7) | *Oryza sativa Japonica Group* | 25 - 30 | 30 | Monomer | Gel filtration chromatography |  | (Opassiri et al., 2013) (Kuntothom et al. 2009)  (Fageria et al., 2011) |
| Eukaryota | β-glucosidase | *Aspergillus niger* | 30 - 35 | 50 | Dimer | Gel filtration chromatography |  | (Watanabe et al., 1992)  (Alborch et al. 2011) |
| Eukaryota | β-thioglucoside hydrolase (TGG1) | *Arabidopsis thaliana* | 23 - 25 | 50 | Dimer | Gel filtration chromatography | Mesophile | (Zhou et al., 2012)  (Rivero-Lepinckas et al., 2006)  (Banerjee et al., 2019) |
| Eukaryota | β-thioglucoside hydrolase (TGG2) | *Arabidopsis thaliana* | 23 - 25 | 50 | Dimer | Gel filtration chromatography | Mesophile | (Zhou et al., 2012)  (Rivero-Lepinckas et al., 2006)  (Banerjee et al., 2019) |
| Eukaryota | β-galactosidase-like enzyme (BglA) | *Sporobolomyces singularis* | 25 | 45 | Dimer | Gel filtration chromatography |  | (Ishikawa et al., 2005)  (Hamamoto et al., 2011) |
| Eukaryota | β-glucosidase (Glu1) | Maize (*Zea mays L.*) | 21 - 27 | 50 | Dimer | Cation exchange chromatography, crystallographic data and gel filtration chromatography |  | (Esen, 1992)(Chang et al., 2013)(Czjzek et al., 2001)(Greaves, 1996) |
| Eukaryota | β-glucosidase | *Candida wickerhamii* | 30 | 35 | Dimer | Gel filtration chromatography |  | (Skory and Freer, 1994) (Freer, 1985)  (Kilian et al., 1983) |
| Eukaryota | Isoflavone conjugate-specific β-glucosidase (GmICHG) | *Glycine max* | 20 - 30 | 30 | Dimer | Gel filtration chromatography |  | (Hsieh and Graham, 2001)  (Hofstra, 1972) (Hesketh et al. 1973) |
| Eukaryota | β-galactosidase/β-glucosidase | *Hamamotoa singularis ATCC 24193* | 25 | 45 | Dimer | Gel filtration chromatography |  | (Ishikawa et al. 2005)  (Rodrigues de Miranda, 1984) |
| Eukaryota | β-glucosidase | *Malus domestica* | 21 - 26 | 70 | Dimer | Gel filtration chromatography |  | (Yu et al. 2007)  (Calderon-Zavala et al. 2004) |
| Eukaryota | β-glucosidase (ICHG) | *Cyamopsis tetragonoloba* (guar) | 25 - 35 | 37 | Trimer | Gel filtration chromatography |  | (Asati and Sharma, 2019)  (Gresta et al., 2018) |
| Eukaryota | β -glucosidase gene (bgl2) | *Trichoderma reesei* | 25 - 30 | 40 | Tetramer | Gel filtration chromatography | Mesophile | (Takashima et al., 1996)  (Takashima et al., 1999)  (Singh et al., 2014)  (Adav et al., 2012) |
| Eukaryota | Isoflavonoid 7-O-β-apiosyl-β-glucose β-glycosidase | *Dalbergia nigrescens KURZ* | 30.5 | 65 | Tetramer | Gel filtration chromatography |  | (Chuankhayan et al., 2005)  (Ferraz-Grande and Takaki, 2001) |
| Eukaryota | β-1,4-glucosidase (BGL) | *Fusarium oxysporum* | 25 | 60 | Pentamer | Gel filtration chromatography | Mesophile | (Zhao et al., 2013b)  (Hibar et al., 2006)  (Liu et al., 2006) |
| Eukaryota | β-glucosidase | *Secale cereale* | 20 | 25-30 | Hexamer | Gel filtration chromatography |  | (Sue et al. 2000) (Sue et al. 2006)  (White et al. 1991) |
| Eukaryota | β-glucosidase  (NfBGL595) | *Neosartorya fischeri NRRL181* | 26 - 45 | 40 | Octamer | Gel filtration chromatography | Thermophile | (Ramachandran et al., 2012)  (Nielsen et al., 1988)  (Järv et al., 2004) |
| Eukaryota | β-glucosidase | *Aspergillus fischeri* | 37 | 40 | Octamer | Gel filtration chromatography |  | (Ramachandran et al. 2012)  (Lamoth, 2016) |
| Eukaryota | *Strictosidine β-glucosidase* | *Catharanthus roseus* | 20 - 27 | 30 | Multimeric | Native PAGE |  | (Geerlings et al., 2000) (Hemscheidt and Zenk, 1980)  (Pandey, 2017) |
| Eukaryota | Myrosinase (TGG4) | *Arabidopsis thaliana* | 23 - 25 | 70 | Not reported |  | Mesophile | (Andersson et al., 2009)  (Rivero-Lepinckas et al., 2006)  (Banerjee et al., 2019) |
| Eukaryota | Myrosinase (TGG5) | *Arabidopsis thaliana* | 23 - 25 | 60 | Not reported |  | Mesophile | (Andersson et al., 2009)  (Rivero-Lepinckas et al., 2006)  (Banerjee et al., 2019) |
| Eukaryota | β-glucosidase  (Cel1B) | Trichoderma reesei | 25 - 30 | 40 | Not reported |  | Mesophile | (Guo et al., 2016)  (Singh et al., 2014)  (Adav et al., 2012) |
| Eukaryota | β -glucosidase (*Cg*BG1) | *Coptotermes gestroi* | 33.6 - 38.6 | 56 | Not reported |  |  | (Franco Cairo et al., 2013)  (Cao and Su, 2016) |
| Eukaryota | β-glucosidase | *Bombyx mori* | 23 - 28 | 35 | Not reported |  |  | (Byeon et al., 2005)  (Rahmathulla, 2012) |
| Eukaryota | β-primeverosidase | *Camellia sinensis* | 20 | 45 | Not reported |  |  | (Ijima et al., 1998)  (Lu et al., 2019) |
| Eukaryota | Ipecoside β-glucosidase | *Carapichea ipecacuanha* | 23 - 25 | 55 - 60 | Not reported |  |  | (Nomura et al., 2008)  (Trujillo et al. 2017) |
| Eukaryota | Myrosinase (TGG1) | *Carica papaya* | 22 - 26 | 40 | Not reported |  |  | (Nong et al. 2010)  (Allan and Jager, 1978) |
| Eukaryota | Myrosinase (TGG2) | *Carica papaya* | 22 - 26 | 40 | Not reported |  |  | (Wang et al., 2009)  (Allan and Jager, 1978) |
| Eukaryota | β-glucosidase | *Coptotermes formosanus* | 30.2 - 35.7 | 42 - 54 | Not reported |  |  | (Zhang et al. 2010)  (Cao and Su, 2016) |
| Eukaryota | Acyl-glucose-dependent anthocyanin β-glucosyltransferase (DgAA7BG-GT1) | *Delphinium grandiflorum* | 15 - 25 | 35 | Not reported |  |  | (Nishizaki et al. 2013)  (Sasaki et al.) |
| Eukaryota | Acyl-glucose-dependent anthocyanin β-glucosyltransferase (DgAA7BG-GT2) | *Delphinium grandiflorum* | 15 - 25 | 35 | Not reported |  |  | (Nishizaki et al. 2013)  (Sasaki et al. 2007) |
| Eukaryota | Acyl-glucose–dependent anthocyanin 5(7)-O-glucosyltransferase (Dg AA7GT) | *Delphinium grandiflorum* | 15 - 25 | 40 | Not reported |  |  | (Matsuba et al., 2010)  (Sasaki et al.,2007) |
| Eukaryota | Rutinase | *Fagopyrum tataricum* | 30 | 50 | Not reported |  |  | (Jia et al., 2019)  (Xoxiong et al. 2011) |
| Eukaryota | β-mannosidase / β-glucosidase (HvBII) | *Hordeum vulgare subsp. vulgare* | 18 | 30 | Not reported |  |  | (Kuntothom et al. 2009)  (Fageria et al., 2011) |
| Eukaryota | β-glucosidase (Os3bglu8) | *Oryza sativa Japonica Group* | 25 - 30 | 30 | Not reported |  |  | (Kuntothom et al. 2009)  (Fageria et al., 2011) |
| Eukaryota | β-glucosidase (Os7bglu26) | *Oryza sativa Japonica Group* | 25 - 30 | 40 | Not reported |  |  | (Kuntothom et al. 2009)  (Fageria et al., 2011) |
| Eukaryota | β-glucosidase | *Phialophora sp. G5 / CGMCC 3328* | 27 - 30 | 50 | Not reported |  |  | (Li et al. 2013)  (Li et al. 2017) |
| Eukaryota | Myrosinase | *Raphanus sativus* | 23 | 37 | Not reported |  |  | (Jwanny et al. 1995)  (Nieuwhof, 1976) |
| Eukaryota | β-glucosidase (*Llbglu1*) | *Leucaena leucocephala (subabul)* | 25 - 30 | 45 | Not reported |  |  | (Shaik et al., 2013)  (Shelton and Brewbaker, 1994) |
| Eukaryota | Acyl-glucose–dependent anthocyanin 5(7)-O-glucosyltransferase (Dc AA5GT) | *Dianthus caryophyllus* | 10 -15 | 35 – 40 | Not reported |  |  | (Matsuba et al., 2010)  (Wan et al., 2015) |
| Eukaryota | β -mannosidase | *Lycopersicon esculentum Mill* | 18 - 28 | 40 | Not reported |  |  | (Mo and Bewley, 2002)  (Saeed et al., 2007) |

Adav, S.S., Chao, L.T., and Sze, S.K. (2012). Quantitative secretomic analysis of Trichoderma reesei strains reveals enzymatic composition for lignocellulosic biomass degradation. Mol. Cell. Proteomics 11, M111.012419.

Aguilar, C.F., Sanderson, I., Moracci, M., Ciaramella, M., Nucci, R., Rossi, M., and Pearl, L.H. (1997). Crystal structure of the β-glycosidase from the hyperthermophilic archeon sulfolobus solfataricus: Resilience as a key factor in thermostability. J. Mol. Biol. 271, 789–802.

Akinosho, H., Yee, K., Close, D., and Ragauskas, A. (2014). The emergence of Clostridium thermocellum as a high utility candidate for consolidated bioprocessing applications. Front. Chem. 2, 66.

Alborch, L., Bragulat, M.R., Abarca, M.L., Cabañes, F.J. (2011). Effect of water activity, temperature and incubation time on growth and ochratoxin A production by Aspergillus niger and Aspergillus carbonarius on maize kernels. Int J Food Microbiol 147, 53-57

Allan, P., Jager, J. (1978). Net photosynthesis in macadamia and papaw and the possible alleviation of heat stress. Crop Prod 7, 125-128

An, C.L., Kim, M.K., Kang, T.H., Kim, J., Kim, H., and Yun, H.D. (2012). Cloning and biochemical analysis of β-glucoside utilization (bgl) operon without phosphotransferase system in Pectobacterium carotovorum subsp. carotovorum LY34. Microbiol. Res. 167, 461–469.

An, C.L., Lim, W.J., Hong, S.Y., Shin, E.C., Kim, M.K., Lee, J.R., Park, S.R., Woo, J.G., Lim, Y.P., and Yun, H.D. (2005). Structural and biochemical analysis of the asc operon encoding 6-phospho-β-glucosidase in Pectobacterium carotovorum subsp. carotovorum LY34. Res. Microbiol. 156, 145–153.

Andersson, D., Chakrabarty, R., Bejai, S., Zhang, J., Rask, L., and Meijer, J. (2009). Myrosinases from root and leaves of Arabidopsis thaliana have different catalytic properties. Phytochemistry 70, 1345–1354.

Asati, V., and Sharma, P.K. (2019). Purification and characterization of an isoflavones conjugate hydrolyzing β-glucosidase (ICHG) from Cyamopsis tetragonoloba (guar). Biochem. Biophys. Reports 20, 100669.

Atomi, H., Fukui, T., Kanai, T., Morikawa, M., and Imanaka, T. (2004). Description of Thermococcus kodakaraensis sp. nov., a well studied hyperthermophilic archaeon previously reported as Pyrococcus sp. KOD1. Archaea 1, 263–267.

Bai, A., Zhao, X., Jin, Y., Yang, G., and Feng, Y. (2013). A novel thermophilic β-glucosidase from Caldicellulosiruptor bescii: Characterization and its synergistic catalysis with other cellulases. J. Mol. Catal. B Enzym. 85–86, 248–256.

Bai, Y., Wang, J., Zhang, Z., Yang, P., Shi, P., Luo, H., Meng, K., Huang, H., and Yao, B. (2010). A new xylanase from thermoacidophilic Alicyclobacillus sp. A4 with broad-range pH activity and pH stability. J. Ind. Microbiol. Biotechnol. 37, 187–194.

Balkwill D.L., J.K. Fredrickson, and M.F.R. (2006). Sphingomonas and Related Genera. In The Prokaryotes: An Evolving Electronic Resource for the Microbiological Community.

Banerjee, R., Kumar, G.V., and Kumar, S.P.J. (2019). Omics-based approaches in plant biotechnology.

Barabote, R.D., Xie, G., Leu, D.H., Normand, P., Necsulea, A., Daubin, V., Médigue, C., Adney, W.S., Xin, C.X., Lapidus, A., et al. (2009). Complete genome of the cellulolytic thermophile Acidothermus cellulolyticus IIB provides insights into its ecophysiological and evolutionary adaptations. Genome Res. 19, 1033–1042.

Bauer, M.W., Bylina, E.J., Swanson, R. V., and Kelly, R.M. (1996). Comparison of a β-glucosidase and a β-mannosidase from the hyperthermophilic archaeon Pyrococcus furiosus. Purification, characterization, gene cloning, and sequence analysis. J. Biol. Chem. 271, 23749–23755.

Berrin, J., Mclauchlan, W.R., Needs, P., Williamson, G., Puigserver, A., Kroon, P.A., and Juge, N. (2002). Functional expression of human liver cytosolic b -glucosidase in Pichia pastoris. Insights into its role in the metabolism of dietary glucosides. 258, 249–258.

Bidart, G.N., Rodríguez-Díaz, J., Pérez-Martínez, G., and Yebra, M.J. (2018). The lactose operon from Lactobacillus casei is involved in the transport and metabolism of the human milk oligosaccharide core-2 N-acetyllactosamine. Sci. Rep. 8, 7152.

Breves, R., Bronnenmeier, K., Wild, N., Lottspeich, F., Staudenbauer, W.L., Ju¨, J., and Hofemeister, J. (1997). Genes Encoding Two Different-Glucosidases of Thermoanaerobacter brockii Are Clustered in a Common Operon. Appl. Environ. Microbiol. 63, 3902–3910.

Brognaro, H., Almeida, V.M., de Araujo, E.A., Piyadov, V., Santos, M.A.M., Marana, S.R., and Polikarpov, I. (2016). Biochemical Characterization and Low-Resolution SAXS Molecular Envelope of GH1 β-Glycosidase from Saccharophagus degradans. Mol. Biotechnol. 58, 777–788.

Busse, H.-J. (2015). Micrococcus. In Bergey’s Manual of Systematics of Archaea and Bacteria.

Byeon, G.M., Lee, K.S., Gui, Z.Z., Kim, I., Kang, P.D., Lee, S.M., Sohn, H.D. and Jin B.R. (2005). A digestive beta-glucosidase from the silkworm, Bombyx mori: cDNA cloning, expression and enzymatic characterization. Comp Biochem Physiol B Biochem Mol Biol 141: 418-427

Cairo, J.P.L.F., Oliveira, L.C., Uchima, C.A., Alvarez, T.M., Citadini, A.S., Cota, J., Leonardo, F.C., Costa-Leonardo, A.M., Carazzolle, M.F., Costa, F.F., Pereira, G.A.G. and Squina, F.M. (2013). Deciphering the synergism of endogenous glycoside hydrolase families 1 and 9 from Coptotermes gestroi. Insect Biochem Molec 43, 970-981

Calderon-Zavala, G., Lakso, A.N. and Piccioni R.M. (2004). Temperature effects on fruit and shoot growth in the apple (Malus domestica) early in the season. Acta Hortic 636, 447-453

Cao, H., Zhang, Y., Shi, P., Ma, R., Yang, H., Xia, W., Cui, Y., Luo, H., Bai, Y., and Yao, B. (2018). A highly glucose-tolerant GH1 β-glucosidase with greater conversion rate of soybean isoflavones in monogastric animals. J. Ind. Microbiol. Biotechnol. 45, 369–378.

Cao, R. and Su, N.Y. (2016). Temperature preferences of four subterranean termite species (Isoptera: Rhinotermitidae) and temperature dependent survivorship and wood-consumption rate. Ann Entomol Soc Am 109, 64-71

Capaldo, A., Walker, M.E., Ford, C.M., and Jiranek, V. (2011). β-Glucoside metabolism in Oenococcus oeni: Cloning and characterization of the phospho-β-glucosidase CelD. J. Mol. Catal. B Enzym. 69, 27–34.

Cayol, J.-L., Ollivier, B., and Garcia, J.-L. (2015). Halothermothrix. In Bergey’s Manual of Systematics of Archaea and Bacteria.

Chai, Y.Y., Kahar, U.M., Salleh, M.M., Illias, R.M., and Mau Goh, K. (2012). Isolation and characterization of pullulan-degrading Anoxybacillus species isolated from Malaysian hot springs. Environ. Technol. (United Kingdom) 33, 1231–1238.

Chan, C.S., Sin, L.L., Chan, K.G., Shamsir, M.S., Manan, F.A., Sani, R.K., and Goh, K.M. (2016). Characterization of a glucose-tolerant β-glucosidase from Anoxybacillus sp. DT3-1. Biotechnol. Biofuels 9, 174.

Chang, J., Lee, Y.S., Fang, S.J., Park, I.H., and Choi, Y.L. (2013). Recombinant expression and characterization of an organic-solvent-tolerant α-amylase from exiguobacterium sp. DAU5. Appl. Biochem. Biotechnol. 169, 1870–1883.

Chang, J., Park, I.H., Lee, Y.S., Ahn, S.C., Zhou, Y., and Choi, Y.L. (2011). Cloning, expression, and characterization of β-glucosidase from Exiguobacterium sp. DAU5 and transglycosylation activity. Biotechnol. Bioprocess Eng. 16, 97–106.

Chuankhayan, P., Hua, Y., Svasti, J., Sakdarat, S., Sullivan, P.A. and Cairns, J.R.K. (2005). Purification of an isoflavonoid 7-O-β-apiosyl-glucoside β-glycosidase and its substrates from Dalbergia nigrescens Kurz. Phytochemistry 66, 1880-1889

Clarke, J.H., Davidson, K., Gilbert, H.J., Fontes, C.M.G.A., and Hazlewood, G.P. (1996). A modular xylanase from mesophilic Cellulomonas fimi contains the same cellulose-binding and thermostabilizing domains as xylanases from thermophilic bacteria . FEMS Microbiol. Lett. 139, 27–35.

Cui, C.-H., Kim, J.-K., Kim, S.-C., and Im, W.-T. (2014). Characterization of a Ginsenoside-Transforming β-glucosidase from Paenibacillus mucilaginosus and Its Application for Enhanced Production of Minor Ginsenoside F2. PLoS One 9, e85727.

Czjzek, M., Cicek, M., Zamboni, R., Burmeister, W.P., Bevan, D.R., Henrissat, B., and Esen, A. (2001). Crystal structure of a monocotyledon (maize ZMGlu1) β-glucosidase and a model of its complex with p-nitrophenyl β-D-thioglucoside. Biochem. J 354, 37–46.

da Costa, M.S., Fernanda Nobre, M., and Rainey, F.A. (2015). Thermus. In Bergey’s Manual of Systematics of Archaea and Bacteria.

Da̧browski, S., Maciuńska, J., and Synowiecki, J. (1998). Cloning and nucleotide sequence of the thermostable β-galactosidase gene from Pyrococcus woesei in Escherichia coli and some properties of the isolated enzyme. Appl. Biochem. Biotechnol. - Part B Mol. Biotechnol. 10, 217–222.

Da̧browski, S.S., Sobiewska, G., Maciuńska, J., Synowiecki, J., and Kur, J. (2000). Cloning, expression, and purification of the His6-tagged thermostable β-galactosidase from Pyrococcus woesei in Escherichia coli and some properties of the isolated enzyme. Protein Expr. Purif. 19, 107–112.

Daniels, L.B., Coyle, P.J., Chiao, Y.B., Glew, R.H., and Labow, R.S. (1981). Purification and characterization of a cytosolic broad specificity β-glucosidase from human liver. J. Biol. Chem. 256, 13004–13013.

Darland, G., and Brock, T.D. (1971). Bacillus acidocaldarius sp.nov., an Acidophilic Thermophilic Spore-forming Bacterium. J. Gen. Microbiol. 67, 9–15.

De Farias, S.T., and Bonato, M.C.M. (2002). Preferred codons and amino acid couples in hyperthermophiles. Genome Biol. 3, preprint0006.1.

Dicks, L.M.T., and Holzapfel, W.H. (2015). Oenococcus. In Bergey’s Manual of Systematics of Archaea and Bacteria.

Ding, Y.H.R., Ronimus, R.S., and Morgan, H.W. (2000). Sequencing, cloning, and high-level expression of the pfp gene, encoding a PP(i)-dependent phosphofructokinase from the extremely thermophilic eubacterium Dictyoglomus thermophilum. J. Bacteriol. 182, 4661–4666.

Dion, M., Fourage, L., Hallet, J.N., and Colas, B. (1999). Cloning and expression of a beta-glycosidase gene from Thermus thermophilus. Sequence and biochemical characterization of the encoded enzyme. Glycoconj. J. 16, 27–37.

Ekborg, N.A., Gonzalez, J.M., Howard, M.B., Taylor, L.E., Hutcheson, S.W., and Weiner, R.M. (2005). Saccharophagus degradans gen. nov., sp. nov., a versatile marine degrader of complex polysaccharides. Int. J. Syst. Evol. Microbiol. 55, 1545–1549.

Esen, A. (1992). Purification and Partial Characterization of Maize (Zea mays L.) β-Glucosidase. Plant Physiol 98, 174–182.

Ezaki, S., Miyaoku, K., Nishi, K.I., Tanaka, T., Fujiwara, S., Takagi, M., Atomi, H., and Imanaka, T. (1999). Gene analysis and enzymatic properties of thermostable β-glycosidase from Pyrococcus kodakaraensis KOD1. J. Biosci. Bioeng. 88, 130–135.

Fageria, N.K., Baligar, V.C., and Jones, C.A. (2011). Growth and mineral nutrition of field crops.

Fan, H.X., Miao, L.L., Liu, Y., Liu, H.C., and Liu, Z.P. (2011). Gene cloning and characterization of a cold-adapted β-glucosidase belonging to glycosyl hydrolase family 1 from a psychrotolerant bacterium Micrococcus antarcticus. Enzyme Microb. Technol. 49, 94–99.

Fang, S., Chang, J., Lee, Y.S., Guo, W., Choi, Y.L., and Zhou, Y. (2014). Cloning and characterization of a new broadspecific β-glucosidase from Lactococcus sp. FSJ4. World J. Microbiol. Biotechnol. 30, 213–223.

Fergus G. Priest, H. (2015). Paenibacillus. In Bergey’s Manual of Systematics of Archaea and Bacteria.

Ferraz-Grande, F.G.A. and Takaki, M. (2001). Temperature dependent seed germination of Dalbergia nigra allem (Leguminosae). Braz Arch Biol Technol 44, Dec 2001

Fiala, G., and Stetter, K.O. (1986). Pyrococcus furiosus sp. nov. represents a novel genus of marine heterotrophic archaebacteria growing optimally at 100°C. Arch. Microbiol. 145, 56–61.

Franco Cairo, J.P.L., Oliveira, L.C., Uchima, C.A., Alvarez, T.M., Citadini, A.P. da S., Cota, J., Leonardo, F.C., Costa-Leonardo, A.M., Carazzolle, M.F., Costa, F.F., et al. (2013). Deciphering the synergism of endogenous glycoside hydrolase families 1 and 9 from Coptotermes gestroi. Insect Biochem. Mol. Biol. 43, 970–981.

Freer, S.N. (1985). Purification and characterization of the extracellular β-glucosidase produced by Candida wickerhamii. Arch Biochem Biophys 232, 512-522

Fusco, F.A., Fiorentino, G., Pedone, E., Contursi, P., Bartolucci, S., and Limauro, D. (2018). Biochemical characterization of a novel thermostable β-glucosidase from Dictyoglomus turgidum. Int. J. Biol. Macromol. 113, 783–791.

Galardini, M., Bazzicalupo, M., Biondi, E., Brambilla, E., Brilli, M., Bruce, D., Chain, P., Chen, A., Daligault, H., Davenport, K.W., et al. (2013). Permanent draft genome sequences of the symbiotic nitrogen fixing Ensifer meliloti strains BO21CC and AK58. Stand. Genomic Sci. 9, 325-333.

Gao, J., and Wakarchuk, W. (2014). Characterization of five β-glycoside hydrolases from Cellulomonas fimi ATCC 484. J. Bacteriol. 196, 4103–4110.

Ge, Y.D., Cao, Z.Y., Wang, Z. Da, Chen, L.L., Zhu, Y.M., and Zhu, G.P. (2010). Identification and biochemical characterization of a thermostable malate dehydrogenase from the mesophile Streptomyces coelicolor A3(2). Biosci. Biotechnol. Biochem. 74, 2194–2201.

Geerlings, A., Ibañez, M.M.L., Memelink, J., van der Heijden, R. and Verpoorte, R. (2000). Molecular cloning and analysis of strictosidine β-D-glucosidase, an enzyme in terpenoid indole alkaloid biosynthesis in Catharanthus roseus. J Biol Chem 275, 3051-3056

Gordon, A. (2017). Case study: Addressing the problem of Alicyclobacillus in tropical beverages. In Food Safety and Quality Systems in Developing Countries.

Goswami, S., Das, S., and Datta, S. (2017). Understanding the role of residues around the active site tunnel towards generating a glucose-tolerant β-glucosidase from Agrobacterium tumefaciens 5A. Protein Eng. Des. Sel. 30, 523–530.

Goswami, S., Gupta, N., and Datta, S. (2016). Using the β-glucosidase catalyzed reaction product glucose to improve the ionic liquid tolerance of β-glucosidases. Biotechnol. Biofuels 9, 72.

Greaves, J.A. (1996). Improving suboptimal temperature tolerance in maize-the search for variation. J. Exp. Bot. 47, 307–323.

Gresta, F., Cristaudo, A., Trostle, C., Anastasi, U., Guarnaccia, P., Catara, S., and Onofri, A. (2018). Germination of guar (Cyamopsis tetragonoloba (L.) Taub.) genotypes with reduced temperature requirements. Aust. J. Crop Sci. 12, 954–960.

Gu, M.Z., Wang, J.C., Liu, W.B., Zhou, Y., and Ye, B.C. (2013). Expression and displaying of β-glucosidase from Streptomyces coelicolor A3 in Escherichia coli. Appl. Biochem. Biotechnol. 170, 1713–1723.

Gu, N., Kim, J., Kim, H., You, D., Kim, H., Jeon, S., and Ioeng, J.B.I.B. (2009). Gene cloning and enzymatic properties of hyperthermostable β -glycosidase from Thermus thermophilus HJ6. JBIOSC 107, 21–26.

Gumerov, V.M., Rakitin, A.L., Mardanov, A. V., and Ravin, N. V. (2015). A Novel Highly Thermostable Multifunctional Beta-Glycosidase from Crenarchaeon Acidilobus saccharovorans. Archaea 2015, 978632.

Guo, B., Sato, N., Biely, P., Amano, Y., and Nozaki, K. (2016). Comparison of catalytic properties of multiple β-glucosidases of Trichoderma reesei. Appl. Microbiol. Biotechnol. 100, 4959–4968.

Hamamoto, M., Boekhout, T., and Nakase, T. (2011). Sporobolomyces Kluyver & van Niel (1924). In The Yeasts.

Hammes, W.P., and Hertel, C. (2015). Lactobacillus. In Bergey’s Manual of Systematics of Archaea and Bacteria.

Haq, I.U., Khan, M.A., Muneer, B., Hussain, Z., Afzal, S., Majeed, S., Rashid, N., Javed, M.M., and Ahmad, I. (2012). Cloning, characterization and molecular docking of a highly thermostable β-1,4-glucosidase from Thermotoga petrophila. Biotechnol. Lett. 34, 1703–1709.

Harada, K.M., Tanaka, K., Fukuda, Y., Hashimoto, W., and Murata, K. (2005). Degradation of rice bran hemicellulose by Paenibacillus sp. strain HC1: Gene cloning, characterization and function of β-D-glucosidase as an enzyme involved in degradation. Arch. Microbiol. 184, 215–224.

Hashimoto, W., Miki, H., Nankai, H., Sato, N., Kawai, S., and Murata, K. (1998). Molecular cloning of two genes for beta-D-glucosidase in Bacillus sp. GL1 and identification of one as a gellan-degrading enzyme. Arch. Biochem. Biophys. 360, 1–9.

Hassan, N., Nguyen, T.H., Intanon, M., Kori, L.D., Patel, B.K.C., Haltrich, D., Divne, C., and Tan, T.C. (2015). Biochemical and structural characterization of a thermostable β-glucosidase from Halothermothrix orenii for galacto-oligosaccharide synthesis. Appl. Microbiol. Biotechnol. 99, 1731–1744.

Hauben, L., Gijsegem, F. Van, and Swings, J. (2015). Pectobacterium. In Bergey’s Manual of Systematics of Archaea and Bacteria.

Hemscheidt, T. and Zenk, M.H. (1980). Glucosidases involved in indole alkaloid biosynthesis of Catharantus cell cultures. FEBS Lett 110, 187-191

Henne, A., Brüggemann, H., Raasch, C., Wiezer, A., Hartsch, T., Liesegang, H., Johann, A., Lienard, T., Gohl, O., Martinez-Arias, R., et al. (2004). The genome sequence of the extreme thermophile Thermus thermophilus. Nat. Biotechnol. 22, 547–553.

Hesketh, J.D., Myhre, D.L. and Willey, C.R. (1973). Temperature control of time intervals between vegetative and reproductive events in soybeans. Crop Sci 13, 250-254

Hibar, K., Daami-Remadi, M., Jabnoun-Khiareddine, H., and El Mahjoub, M. (2006). Temperature effect on mycelial growth and on disease incidence of Fusarium oxysporum f.sp. radicis-lycopersici. Plant Pathol. J. 5, 233–238.

Hofsta, G. (1972). Response of soybeans to temperature under high light intensities. Can J Plant Sci 52, 535-543

Honda, H., Nagaoka, S., Kawai, Y., Kemperman, R., Kok, J., Yamazaki, Y., Tateno, Y., Kitazawa, H., and Saito, T. (2012). Purification and characterization of two phospho-β-galactosidases, LacG1 and LacG2, from Lactobacillus gasseri ATCC33323(T). J. Gen. Appl. Microbiol. 58, 11–17.

Hong, M.R., Kim, Y.S., Park, C.S., Lee, J.K., Kim, Y.S., and Oh, D.K. (2009). Characterization of a recombinant β-glucosidase from the thermophilic bacterium Caldicellulosiruptor saccharolyticus. J. Biosci. Bioeng. 108, 36–40.

Hong, S.Y., Cho, K.M., Math, R.K., Kim, Y.H., Hong, S.J., Cho, Y.U., Kim, H., and Yun, H.D. (2007). Characterization of the recombinant cellobiase from celG gene in the beta-glucoside utilization gene operon of Pectobacterium carotovorum subsp. carotovorum LY34. J. Mol. Catal. B Enzym. 47, 91–98.

Hsieh, M.C. and Graham, T.L. (2001). Partial purification and characterization of a soybean beta-glucosidase with high specific activity towards isoflavone conjugates. Phytochemistry 58, 995-1005

Huang, C.Y., Patel, B.K., Mah, R.A., and Baresi, L. (1998). Caldicellulosiruptor owensensis sp. nov., an anaerobic, extremely thermophilic, xylanolytic bacterium. Int. J. Syst. Bacteriol. 48, 91–97.

Huber, H., and Stetter, K.O. (2015). Sulfolobus. In Bergey’s Manual of Systematics of Archaea and Bacteria.

Huber, R., and Stetter, K.O. (2015b). Fervidobacterium. In Bergey’s Manual of Systematics of Archaea and Bacteria.

Huber, R., Woese, C.R., Langworthy, T.A., Kristjansson, J.K., and Stetter, K.O. (1990). Fervidobacterium islandicum sp. nov., a new extremely thermophilic eubacterium belonging to the “Thermotogales.” Arch. Microbiol. 154, 105–111.

Hwa, K.Y., Subramani, B., Shen, S.T., and Lee, Y.M. (2014). An intermolecular disulfide bond is required for thermostability and thermoactivity of β-glycosidase from Thermococcus kodakarensis KOD1. Appl. Microbiol. Biotechnol. 98, 7825–7836.

Hwa, K.Y., Subramani, B., Shen, S.T., and Lee, Y.M. (2015). Exchange of active site residues alters substrate specificity in extremely thermostable β-glycosidase from Thermococcus kodakarensis KOD1. Enzyme Microb. Technol. 77, 14–20.

Ijima, Y., Ogawa, K., Watanabe, N., Usui, T., Ohnishi-Kameyama, M., Nagata, T. and Sakata, K. (1998). Characterization of a β-primeverosidase, being concerned with alcoholic aroma formation in tea leaves to be processed into black tea, and preliminary observations on its substrate specificity. J Agric Food Chem. 46: 1712-1718

Ishikawa, E., Sakai, T., Ikemura, H., Matsumoto, K. and Abe, H. (2005). Identification, cloning, and characterization of a Sporobolomyces singularis beta-galactosidase-like enzyme involved in galacto-oligosaccharide production. J Biosci Bioeng 99, 331-339

Itoh, T., Suzuki, K., Sanchez, P.C., and Nakase, T. (2015). Caldivirga. In Bergey’s Manual of Systematics of Archaea and Bacteria.

Jabbour, D., Klippel, B., and Antranikian, G. (2012). A novel thermostable and glucose-tolerant β-glucosidase from Fervidobacterium islandicum. Appl. Microbiol. Biotechnol. 93, 1947–1956.

Jang, W.J., Lee, J.M., Kim, Y.R., Hasan, M.T., and Kong, I.S. (2018). Complete Genome Sequence of Bacillus sp. SJ-10 (KCCM 90078) Producing 400-kDa Poly-γ-glutamic Acid. Curr. Microbiol. 75, 1378–1383.

Järv, H., Lehtmaa, J., Summerbell, R.C., Hoekstra, E.S., Samson, R.A., and Naaber, P. (2004). Isolation of Neosartorya pseudofischeri from Blood: First Hint of Pulmonary Aspergillosis. J. Clin. Microbiol. 42, 925–928.

Jia, P., Wang, Y., Niu, Y., Han, X., Zhu, Y., Xu, Q., Li, Y and Chen, P. (2019). loning and molecular characterization of rutin degrading enzyme from tartary buckwheat (Fagopyrum tataricum Gaertn.). Plan Physiol Bioch 143, 61-71

John G. Holt, Krieg, N.R., Sneath, P.H.A., Staley, J.T., and Williams, S.T. (1994). Bergey’s Manual of Determinative Bacteriology.

Jwanny, E.W., El-Sayed, S.T., Rashad, M.M., Mahmoud, A.E. and Abdallah, N.M. (1995). Myrosinase from roots of Raphanus sativus. Phytochemistry 6, 1301-1303

Kämpfer, P. (2015). Streptomyces. In Bergey’s Manual of Systematics of Archaea and Bacteria.

Kämpfer, P., Glaeser, S.P., Parkes, L., Keulen, G. van, and Dyson, P. (2014). The Family Streptomycetaceae. In The Prokaryotes: Actinobacteria, pp. 1–1061.

Kang, S.K., Cho, K.K., Ahn, J.K., Bok, J.D., Kang, S.H., Woo, J.H., Lee, H.G., You, S.K., and Choi, Y.J. (2005). Three forms of thermostable lactose-hydrolase from Thermus sp. IB-21: Cloning, expression, and enzyme characterization. J. Biotechnol. 117, 337–346.

Kang, S.K., Cho, K.K., Ahn, J.K., Kang, S.H., Han, K.H., Lee, H.G., and Choi, Y.J. (2004). Cloning and expression of thermostable β-glycosidase gene from Thermus filiformis Wai33 A1 in Escherichia coli and enzyme characterization. J. Microbiol. Biotechnol. 14, 584–592.

Kengen, S.W.M., Luesink, E.J., Stams, A.J.M., and Zehnder, A.J.B. (1993). Purification and characterization of an extremely thermostable β‐glucosidase from the hyperthermophilic archaeon Pyrococcus furiosus. Eur. J. Biochem. 213, 305–312.

Kilian, S.G., Prior, B.A., Pretorius, I.S., du Preez, J.C., Venter, J.J. and Potgieter, H.J. (1983). Nutritional, temperature, pH, and oxygen requirements of Candida wickerhamii. Eur J Appl Microbiol Biotechnol 17, 334-338

Kim, J., Lee, J., Shin, Y., and Kim, G. (2010). Characterization of an indican-hydrolyzing enzyme from Sinorhizobium meliloti. Process Biochem. 45, 892–896.

Kim, M.K., An, C.L., Kang, T.H., Kim, J., Kim, H., and Yun, H.D. (2013). Activation of a casB gene encoding β-glucosidase of Pectobacterium carotovorum subsp. carotovorum LY34. Microbiol. Res. 168, 138–146.

Kim, S.B. (2015). Thermobispora. In Bergey’s Manual of Systematics of Archaea and Bacteria.

Kim, Y.J., Lee, J.E., Lee, H.S., Kwon, K.K., Kang, S.G., and Lee, J.H. (2015). Novel substrate specificity of a thermostable β-glucosidase from the hyperthermophilic archaeon, Thermococcus pacificus P-4. Korean J. Microbiol. 51, 68–74.

Kobayashi, T. (2015). Thermococcus. In Bergey’s Manual of Systematics of Archaea and Bacteria.

Kosugi, A., Arai, T., and Doi, R.H. (2006). Degradation of cellulosome-produced cello-oligosaccharides by an extracellular non-cellulosomal β-glucan glucohydrolase, BglA, from Clostridium cellulovorans. Biochem. Biophys. Res. Commun. 349, 20–23.

Kristjansson, J.K., Hreggvidsson, G. 0, and Alfredsson, G.A. (1986). Isolation of Halotolerant Thermus spp. from Submarine Hot Springs in Iceland. Appl. Environ. Microbiol. 52, 1313–1316.

Kumar, P., Ryan, B., and Henehan, G.T.M. (2017). β-Glucosidase from Streptomyces griseus: Nanoparticle immobilisation and application to alkyl glucoside synthesis. Protein Expr. Purif. 132, 164–170.

Kuntothom, T., Luang, S., Harvey, A.J., Fincher, G.B., Opassiri, R., Hrmova, M., Ketudat-Cairns, J.R. (2009). Rice family GH1 glycoside hydrolases with beta-D-glucosidase and beta-D-mannosidase activities. Arch Biochem Biophys 491, 85-95

Kurniasih, S.D., Alfi, A., Natalia, D., Radjasa, O.K., and Nurachman, Z. (2014). Construction of individual, fused, and co-expressed proteins of endoglucanase and β-glucosidase for hydrolyzing sugarcane bagasse. Microbiol. Res. 169, 725–732.

Kuykendall, L.D., Hashem, F.M., and Wang, E.T. (2015). Sinorhizobium. In Bergey’s Manual of Systematics of Archaea and Bacteria.

Lacey, J. (1978). Thermophilic actinomycetes: Characteristics and identification. J. Allergy Clin. Immunol. 61, 231–232.

Lamoth, G. (2016). Aspergillus fumigatus-Related Species in Clinical Practice. Front Microbiol 7, 683

Lauro, B. Di, Rossi, M., and Moracci, M. (2006). Characterization of a β-glycosidase from the thermoacidophilic bacterium Alicyclobacillus acidocaldarius. Extremophiles 10, 301–310.

Lavermicocca, P., Valerio, F., De Bellis, P., Sisto, A., and Leguérinel, I. (2016). Sporeforming bacteria associated with bread production: Spoilage and toxigenic potential. In Food Hygiene and Toxicology in Ready-to-Eat Foods.

Leah, R., Kigel, J., Svendsen, I., and Mundy, J. (1995). Biochemical and molecular characterization of a barley seed β- glucosidase. J. Biol. Chem. 270, 15789–15797.

Lee, D.W., Choe, E.A., Kim, S.B., Eom, S.H., Hong, Y.H., Lee, S.J., Lee, H.S., Lee, D.Y., and Pyun, Y.R. (2005). Distinct metal dependence for catalytic and structural functions in the l-arabinose isomerases from the mesophilic Bacillus halodurans and the thermophilic Geobacillus stearothermophilus. Arch. Biochem. Biophys. 434, 333–343.

Lee, J.M., Kim, Y.R., Kim, J.K., Jeong, G.T., Ha, J.C., and Kong, I.S. (2015). Characterization of salt-tolerant β-glucosidase with increased thermostability under high salinity conditions from Bacillus sp. SJ-10 isolated from jeotgal, a traditional Korean fermented seafood. Bioprocess Biosyst. Eng. 38, 1335–1346.

Letsididi, R., Hassanin, H.A., Koko, M.Y., Ndayishimiye, J.B., Zhang, T., Jiang, B., Stressler, T., Fischer, L., and Mu, W. (2017). Characterization of a thermostable glycoside hydrolase (CMbg0408) from the hyperthermophilic archaeon Caldivirga maquilingensisIC-167. J. Sci. Food Agric. 97, 2132–2140.

Li, X., Zhao, J., Shi, P., Yang, P., Wang, Y., Luo, H. and Yao, B. (2013). Molecular cloning and expression of a novel β-glucosidase gene from Phialophora sp. G5. Appl Biochem Biotech 169, 941-949

Li, Y., Mingwei, B.U., Peng, C., Xiaohong, L.I., Changwu, C., Gui, G., Feng, Y., Weiwei, H., and Zuoming, Z. (2018). Characterization of a Thermophilic Monosaccharide Stimulated β-Glucosidase from Acidothermus cellulolyticus. Chem. Res. Chin. Univ 34, 212–220.

Li, Y., Xiao, J., de Hoog, G.S., Wang, X., Wan, Z., Yu, J., Liu, W. and Li, R. (2017). Biodiversity and human-pathogenicity of Phialophora verrucosa and relatives in Chaetothyriales. Persoonia 38, 1-19

Liu, H., Xu, Y., Ma, Y., and Zhou, P. (2000). Characterization of Micrococcus antarcticus sp. nov., a psychrophilic bacterium from Antarctica. Int. J. Syst. Evol. Microbiol. 50, 715–719.

Liu, S., Duan, X., Lu, X., and Gao, P. (2006). A novel thermophilic endoglucanase from a mesophilic fungus Fusarium oxysporum. Chinese Sci. Bull. 51, 191–197.

Liu, W., Bevan, D.R., and Zhang, Y.H.P. (2010). The family 1 glycoside hydrolase from Clostridium cellulolyticum H10 is a cellodextrin glucohydrolase. Appl. Biochem. Biotechnol. 161, 264–273.

Liu, Y., Li, R., Wang, J., Zhang, X., Jia, R., Gao, Y., and Peng, H. (2017). Increased enzymatic hydrolysis of sugarcane bagasse by a novel glucose- and xylose-stimulated β-glucosidase from Anoxybacillus flavithermus subsp. yunnanensis E13T. BMC Biochem. 18, 4.

Logan, N.A., and Vos, P. De (2015). Bacillus. In Bergey’s Manual of Systematics of Archaea and Bacteria.

Lu, Y.Z., Liu, P.F., Montazar, A., Paw U, K.T. and Hu, Y.G. (2019). Soil water infiltration model for sprinkler irrigation control strategy: a case for tea plantation in Yangtze river region. Agriculture 9, 206

Madan, M., and Thind, K.S. (1998). Physiology of fungi.

Mandelli, F., Couger, M.B., Paixão, D.A.A., Machado, C.B., Carnielli, C.M., Aricetti, J.A., Polikarpov, I., Prade, R., Caldana, C., Paes Leme, A.F., et al. (2017). Thermal adaptation strategies of the extremophile bacterium Thermus filiformis based on multi-omics analysis. Extremophiles 21, 775–788.

Marques, A.R., Coutinho, P.M., Videira, P., Fialho, A.M., and Sá-Correia, I. (2003). Sphingomonas paucimobilis β-glucosidase Bgl1: A member of a new bacterial subfamily in glycoside hydrolase family 1. Biochem. J. 370, 793–804.

Matsuba, Y., Sasaki, N., Tera, M., Okamura, M., Abe, Y., Okamoto, E., Nakamura, H., Funabashi, H., Takatsu, M., Saito, M., Matsuoka, H., Nagasawa, K. and Ozeki, Y. (2010). A novel glucosylation reaction on anthocyanins catalyzed by acyl-glucose–dependent glucosyltransferase in the petals of Carnation and Delphinium. Plant Cell 22: 3374–3389

Matsui, I., Sakai, Y., Matsui, E., Kikuchi, H., Kawarabayasi, Y., and Honda, K. (2000). Novel substrate specificity of a membrane-bound beta-glycosidase from the hyperthermophilic archaeon Pyrococcus horikoshii. FEBS Lett. 467, 195–200.

Minami, Y., Kanafuji, T. and Miura, K. (1996). Purification and characterization of a β-glucosidase from Polygonum tinctorium, which catalyzes preferentially the hydrolysis of indicant. Biosci Biotech Biochem 60, 147-149

Mo, B., and Bewley, J.D. (2002). β-Mannosidase (EC 3.2.1.25) activity during and following germination of tomato (Lycopersicon esculentum Mill.) seeds. Purification, cloning and characterization. Planta 215, 141–152.

Nam, E.S., Kim, M.S., Lee, H.B., and Ahn, J.K. (2010). Beta-glycosidase of Thermus thermophilus KNOUC202: gene and biochemical properties of the enzyme expressed in Escherichia coli. Prikl. Biokhim. Mikrobiol. 46, 562–571.

Nesbø, C.L., Farrell, A.A., Zhaxybayeva, O., and L’Haridon, S. (2015). Thermotoga. In Bergey’s Manual of Systematics of Archaea and Bacteria.

Nielsen, P.E.R. V, Beuchat, L.R., and Frisvad, J.C. (1988). Growth of and Fumitremorgin Production by Neosartorya fischeri as Affected by Temperature , Light , and Water Activity. Appl. Environ. Microbiol. 54, 1504–1510.

Nieuwhof, M. (1976). The effect of temperature on growth and development of cultivars of radish under winter conditions. Sci Hortic-Amsterdam 5, 111-118

Nishizaki, Y., Yasunaga, M., Okamoto, E., Okamoto, M., Hirose, Y., Yamaguchi, M., Ozeki, Y. and Sasaki, N. (2013). P-hydroxybenzoyl-glucose is a zwitter donor for the biosynthesis of 7-polyacylated anthocyanin in Delphinium. Plant Cell 25: 4150–4165

Nomura, T., Quesada, A.L. and Kutchan, T.M. (2008). The new β-D-Glucosidase in terpenoid-isoquinoline alkaloid biosynthesis in Psychotria ipecacuanha. J Biol Chem 283, 34650-34659

Nong, H., Zhang, J.-M., Li, D.-Q., Wang, M., Sun, X.-P., Zhu, Y.J., Meijer, J. and Wang, Q.-H. (2010). Characterization of a novel β‐thioglucosidase CpTGG1 in Carica papaya and its substrate‐dependent and ascorbic acid‐independent O‐β‐glucosidase activity. J Integr Plant Biol 52, 879-890

Olsen, I. (2014). The family fusobacteriaceae. In The Prokaryotes: Firmicutes and Tenericutes.

Onyenwoke, R.U., and Wiegel, J. (2015a). Thermoanaerobacter. In Bergey’s Manual of Systematics of Archaea and Bacteria.

Opassiri, R., Ketudat-Cairns, J.R., Akiyama, T., Wara-Aswapati, O., Svasti, J. and Esen, A. (2003). Characterization of a riceb-glucosidase highly expressed in flowerand germinating shoot. Plant Sci 165, 627-638

Paavilainen, S., Hellman, J., and Korpela, T. (1993). Purification, characterization, gene cloning, and sequencing of a new beta-glucosidase from Bacillus circulans subsp. alkalophilus. Appl. Environ. Microbiol. 59, 927-932.

Pandey, S. (2017). Catharanthus roseus: Cultivation under stress conditions. In Catharanthus roseus: Current research and future prospects, pp. 383-397

Park, N.-Y., Cha, J., Kim, D.-O., and Park, C.-S. (2007). Enzymatic characterization and substrate specificity of thermostable beta-glycosidase from hyperthermophilic archaea, Sulfolobus shibatae, expressed in E. coli. J. Microbiol. Biotechnol. 17, 454–460.

Patel, B.K.C. (2015). Dictyoglomus. In Bergey’s Manual of Systematics of Archaea and Bacteria.

Pei, J., Pang, Q., Zhao, L., Fan, S., and Shi, H. (2012). Thermoanaerobacterium thermosaccharolyticum β-glucosidase: A glucose-tolerant enzyme with high specific activity for cellobiose. Biotechnol. Biofuels 5, 31.

Peng, X., Su, H., Mi, S., and Han, Y. (2016). A multifunctional thermophilic glycoside hydrolase from Caldicellulosiruptor owensensis with potential applications in production of biofuels and biochemicals. Biotechnol. Biofuels 9, 98.

Perez‐Pons, J.A., Cayetano, A., Rebordosa, X., Lloberas, J., Guasch, A., and Querol, E. (1994). A β‐glucosidase gene (bgl3) from Streptomyces sp. strain QM‐B814: Molecular cloning, nucleotide sequence, purification and characterization of the encoded enzyme, a new member of family 1 glycosyl hydrolases. Eur. J. Biochem. 223, 557–565.

Petitdemange, E., Caillet, F., Giallo, J., and Gaudin, C. (1984). Clostridium cellulolyticum sp. nov., a cellulolytic, mesophilic species from decayed grass. Int. J. Syst. Bacteriol. 34, 155–159.

Pikuta, E. V. (2015). Anoxybacillus. In Bergey’s Manual of Systematics of Archaea and Bacteria.

Priest, F.G., Goodfellow, M., Shute, L.A., and Berkeley3, R.C.W. (1987). Bacillus amyloliquefaciens sp. nov. norn. rev. Int. J. Syst. Bacteriol. 37, 69–71.

Prokofeva, M.I., Kostrikina, N.A., Kolganova, T. V., Tourova, T.P., Lysenko, A.M., Lebedinsky, A. V., and Bonch-Osmolovskaya, E.A. (2009). Isolation of the anaerobic thermoacidophilic crenarchaeote Acidilobus saccharovorans sp. nov. and proposal of Acidilobales ord. nov., including Acidilobaceae fam. nov. and Caldisphaeraceae fam. nov. Int. J. Syst. Evol. Microbiol. 59, 3116–3122.

Radhakrishna, S., Mohinder Singh, M., and John, C.K. (1988). Technology for Rural Development.

Rahmathulla, V.K. (2012). Management of climatic factors for successful silkworm (Bombyx mori L.) crop and higher silk production: A Review. Psyche J Ent 2012, 121234

Rainey, F.A., Hollen, B.J., and Small, A.M. (2015). Clostridium. In Bergey’s Manual of Systematics of Archaea and Bacteria.

Ramachandran, P., Tiwari, M.K., Singh, R.K., Haw, J.R., Jeya, M., and Lee, J.K. (2012). Cloning and characterization of a putative β-glucosidase (NfBGL595) from Neosartorya fischeri. Process Biochem. 47, 99–105.

Riou, C., Salmon, J.-M., Vallier, M.-J., Günata, Z. and Barre, P. (1998). Purification, characterization, and substrate specificity of a novel highly glucose-tolerant β-glucosidase from Aspergillus oryzae. Appl Environ Microbiol 64: 3607-3614

Rivero-Lepinckas, L., Crist, D., and Scholl, R. (2006). Growth of plants and preservation of seeds. Methods Mol. Biol. 323, 3–12.

Robb, F.T., Maeder, D.L., Brown, J.R., DiRuggiero, J., Stump, M.D., Yeh, R.K., Weiss, R.B., and Dunn, D.M. (2001). Genomic sequence of hyperthermophile, Pyrococcus furiosus: implications for physiology and enzymology. Methods Enzymol. 330, 134–157.

Rodrigues de Miranda, L. (1984). Bullera derx. In The Yeasts: A Taxonomic Study, pp. 581-584

Saeed, A., Hayat, K., Khan, A.A., and Iqbal, S. (2007). Heat Tolerance Studies in Tomato (Lycopersicon esculentum Mill.). Int. J. Agric. Biol. 9, 649–652.

SAIKI, T., KIMURA, R., and ARIMA, K. (1972). Isolation and Characterization of Extremely Thermophilic Bacteria from Hot Springs. Agric. Biol. Chem. 36, 2357–2366.

Sakurai, Y., Misawa, S. and Shiota, H. (2014). Growth and respiratory activity of Aspergillus oryzae grown on solid state medium. Agric Biol Chem 49, 745-750

Sang, K.K., Kwang, K.C., Jong, K.A., Seung, H.K., Seung, H.L., Hong, G.L., and Choi, Y.J. (2005). Cloning, expression, and enzyme characterization of thermostable β-glycosidase from Thermus flavus AT-62. Enzyme Microb. Technol. 37, 655–662.

Santos, C.A., Zanphorlin, L.M., Crucello, A., Tonoli, C.C.C., Ruller, R., Horta, M.A.C., Murakami, M.T., and De Souza, A.P. (2016). Crystal structure and biochemical characterization of the recombinant ThBgl, a GH1 Β-glucosidase overexpressed in Trichoderma harzianum under biomass degradation conditions. Biotechnol. Biofuels 9, 71.

Sasaki, K., Nishijima, T., Honda, K., Saga, K., and Sameshima, M. Effects of Difference between Day and Night Temperature and Short Time Temperature Fluctuations on the Growth of Delphinium grandiflorum L. (2007) Hort. Res. (Japan). 6, 577–583.

Schöniger, R., Lindemann, P., Grimm, R., Eckerskorn, C. and Luckner, M. (1998). Cardenolide 16′-O-glucohydrolase from Digitalis lanata. Purification and characterization. Planta 205, 477-482

Seshadri, S., Akiyama, T., Opassiri, R., Kuaprasert, B., and Cairns, J.K. (2009). Structural and enzymatic characterization of Os3BGlu6, a rice beta;-glucosidase hydrolyzing hydrophobic glycosides and (1→3)- and (1→2)-linked disaccharides. Plant Physiol. 151, 47–58.

Shaik, N.M., Misra, A., Singh, S., Fatangare, A.B., Ramakumar, S., Rawal, S.K., and Khan, B.M. (2013). Functional characterization, homology modeling and docking studies of β-glucosidase responsible for bioactivation of cyanogenic hydroxynitrile glucosides from Leucaena leucocephala (subabul). Mol. Biol. Rep. 40, 1351–1363.

Shanshan, Y.U., Yin, H., Wang, X., Feng, L., Chunchun, X.U., Li, J., Hongxiang, H., and Shuying, L. (2014). A Novel Thermostable β-Galactosidase from Geobacillus kaustophilus HTA42. Chem. Res. Chin. Univ 30, 778–784.

Sharma, K., Thakur, A., Kumar, R., and Goyal, A. (2019). Structure and biochemical characterization of glucose tolerant β-1,4 glucosidase (HtBgl) of family 1 glycoside hydrolase from Hungateiclostridium thermocellum. Carbohydr. Res. 483, 107750.

Shelton, H.M., and Brewbaker, J.L. (1994). Leucaena leucocephala - the most widely used forage tree legume. In Forage Tree Legumes in Tropical Agriculture.

Shin, K.C., and Oh, D.K. (2014). Characterization of a novel recombinant β-glucosidase from Sphingopyxis alaskensis that specifically hydrolyzes the outer glucose at the C-3 position in protopanaxadiol-type ginsenosides. J. Biotechnol. 172, 30–37.

Singh, A., Shahid, M., Srivastava, M., Pandey, S., Sharma, A., and Kumar, V. (2014). Optimal Physical Parameters for Growth of Trichoderma species at Varying Virology & Mycology Optimal Physical Parameters for Growth of Trichoderma Species at Varying pH , Temperature and Agitation. Virol. Mycol. 3, 127.

Skory, C.D. and Freer, S.N. (1994). Cloning and characterization of a gene encoding a cell-bound, extracellular β-glucosidase in the yeat Candida wickerhamii. Appl Environ Microb 61, 518-525

Song, X., Xue, Y., Wang, Q., and Wu, X. (2011). Comparison of three thermostable β-glucosidases for application in the hydrolysis of soybean isoflavone glycosides. J. Agric. Food Chem. 59, 1954–1961.

Spiridonov, N.A., and Wilson, D.B. (2001). Cloning and biochemical characterization of BglC, a β-glucosidase from the cellulolytic actinomycete thermobifida fusca. Curr. Microbiol. 42, 295–301.

Stackebrandt, E., and Schumann, P. (2015). Cellulomonas. In Bergey’s Manual of Systematics of Archaea and Bacteria.

Stetter, K.O., and Huber, H. (2015). Pyrococcus. In Bergey’s Manual of Systematics of Archaea and Bacteria.

Sue, M., Ishihara, A. and Iwamura, H. (2000). Purification and characterization of a beta-glucosidase from rye (Secale cereale L.) seedlings. Plant Sci 12, 67-74

Sue, M., Yamazaki, K., Yajima, S., Nomura, T., Matsuwaka, T., Iwamura, H. and Miyamoto, T. (2006). Molecular and structural characterization of hexameric β-D-glucosidases in wheat and rye. Plant Physiol 141, 1237-1247

Takami, H., Nishi, S., Lu, J., Shimamura, S., and Takaki, Y. (2004). Genomic characterization of thermophilic Geobacillus species isolated from the deepest sea mud of the Mariana Trench. Extremophiles 8, 351–356.

Takashima, S., Nakamura, A., Hidaka, M., Masaki, H., and Uozumi, T. (1999). Molecular cloning and expression of the novel fungal β-glucosidase genes from Humicola grisea and Trichoderma reesei. J. Biochem. 125, 728–736.

Takashima, S., Nakamura, A., Masaki, H., and Uozumi, T. (1996). Purification and characterization of cellulases from humicola grisea. Biosci. Biotechnol. Biochem. 60, 77–82.

Tamaru, Y., Miyake, H., Kuroda, K., Nakanishi, A., Matsushima, C., Doi, R.H., and Ueda, M. (2011). Comparison of the mesophilic cellulosome-producing Clostridium cellulovorans genome with other cellulosome-related clostridial genomes. Microb. Biotechnol. 4, 64–73.

Teuber, M. (2015). Lactococcus. In Bergey’s Manual of Systematics of Archaea and Bacteria.

Thompson, J., Robrish, S.A., Bouma, C.L., Freedberg, D.I., and Folk, J.E. (1997). Phospho-β-glucosidase from Fusobacterium mortiferum: Purification, cloning, and inactivation by 6-phosphoglucono-δ-lactone. J. Bacteriol. 179, 1636–1645.

Tiwari, R., Singh, P.K., Singh, S., Nain, P.K.S., Nain, L., and Shukla, P. (2017). Bioprospecting of novel thermostable β-glucosidase from Bacillus subtilis RA10 and its application in biomass hydrolysis. Biotechnol. Biofuels 10, 246.

Trujillo, A.I.U., Giraldo, C.B. and Gomez, E.J.N. (2017). Ex situ potential conservation of ipecacuanha (Psychotria ipecacuanha (Brot.) Stokes.), a critically endangered medicinal plant species. Acta Agron 66, 598-605

Trujillo, M.E., and Goodfellow, M. (2015). Thermobifida. In Bergey’s Manual of Systematics of Archaea and Bacteria.

Uchima, C.A., Tokuda, G., Watanabe, H., Kitamoto, K. and Arioka, M. (2012). Heterologous expression in pichia pastoris and characterization of an endogenous thermostable and high-glucose-tolerant β-glucosidase from the termite Nasutitermes takasagoensis. Appl Environ Microbiol 78, 4288-4293

Victor E. Del Bene. (1990). Temperature. In Clinical Methods: The History, Physical, and Laboratory Examinations.

Vishnivetskaya, T.A., Siletzky, R., Jefferies, N., Tiedje, J.M., and Kathariou, S. (2007). Effect of low temperature and culture media on the growth and freeze-thawing tolerance of Exiguobacterium strains. Cryobiology 54, 234–240.

Wakuta, S., Hamada, S., Ito, H., Matsuura, H., Nabeta, K. and Matsui, H. (2010). Identification of a β-glucosidase hydrolyzing tuberonic acid glucoside in rice (Oryza sativa L.). Phytochemistry 71, 1280-1288

Wan, X.L., Zhou, Q., Wang, Y.Y., Wang, W.E., Bao, M.Z., and Zhang, J.W. (2015). Identification of heat-responsive genes in carnation (Dianthus caryophyllus L.) by RNA-seq. Front. Plant Sci. 6, 519.

Wang Z. Y., Mo J. C., and Lu Y. J (2009). Biology and ecology of Macrotermes barneyi (Isoptera: Termitidae). Sociobiol. 54, 777–786.

Wang, L., Liu, Q.M., Sung, B.H., An, D.S., Lee, H.G., Kim, S.G., Kim, S.C., Lee, S.T., and Im, W.T. (2011). Bioconversion of ginsenosides Rb1, Rb2, Rc and Rd by novel β-glucosidase hydrolyzing outer 3-O glycoside from Sphingomonas sp. 2F2: Cloning, expression, and enzyme characterization. J. Biotechnol. 156, 125–133.

Wang, M., Li, D., Sun, X., Zhu, Y.J., Nong, H. and Zhang, J. (2009). Characterization of a root-specific β-thioglucoside glucohydrolase gene in Carica papaya and its recombinant protein expressed in Pichia pastoris. Plant Sci 177, 716-723

Wang, X., He, X., Yang, S., An, X., Chang, W., and Liang, D. (2003). Structural basis for thermostability of beta-glycosidase from the thermophilic eubacterium Thermus nonproteolyticus HG102. J. Bacteriol. 185, 4248–4255.

Watanabe, T., Sato, T., Yoshioka, S., Koshijima, T. and Kuwahara, M. (1992). Purification and properties of Aspergillus niger β-glucosidase. Eur J Biochem 209: 651-659

White, P.J., Cooper, H.D., Clarkson, D.T., Earnshaw, M.J. and Loughman, B.C. (1991). Effects of low temperature on growth and nutrient accumulation in rye (Secale cereal) and wheat (Triticum aestivum). Ann Bot 67, 23-31

Wiegel, J., and Ljungdahl, L.G. (1981). Thermoanaerobacter ethanolicus gen. nov., spec. nov., a new, extreme thermophilic, anaerobic bacterium. Arch. Microbiol. 128, 343–348.

Windberger, E., Huber, R., Trineone, A., Fricke, H., and Stetter, K.O. (1989). Thermotoga thermarum sp. nov. and Thermotoga neapolitana occurring in African continental solfataric springs. Arch Microbiol 151, 506–512.

Wright, R.M., Yablonsky, M.D., Shalita, Z.P., Goyal, A.K., and Eveleigh, D.E. (1992). Cloning, characterization, and nucleotide sequence of a gene encoding Microbispora bispora BglB, a thermostable β-glucosidase expressed in Escherichia coli. Appl. Environ. Microbiol. 58, 3455–3465.

Wu, Y., Chi, S., Yun, C., Shen, Y., Tokuda, G., and Ni, J. (2012). Molecular cloning and characterization of an endogenous digestive β-glucosidase from the midgut of the fungus-growing termite Macrotermes barneyi. Insect Mol. Biol. 21, 604–614.

Wu, Y., Yuan, S., Chen, S., Wu, D., Chen, J., and Wu, J. (2013). Enhancing the production of galacto-oligosaccharides by mutagenesis of Sulfolobus solfataricus β-galactosidase. Food Chem. 138, 1588–1595.

Xiangyuan, H., Shuzheng, Z., and Shoujun, Y. (2001). Cloning and expression of thermostable β-glycosidase gene from Thermus nonproteolyticus HG102 and characterization of recombinant enzyme. Appl. Biochem. Biotechnol. - Part A Enzym. Eng. Biotechnol. 94, 243–255.

Xoxiong, B., Chang, K.J., Ahn, C.H., Lim, Y.S., Kim, Y.B., Park, S.U., Park, B.J., Sung, I.J. and Park, C.H. (2011). Effect on temperature, deep sea water and seed quality on growth of buckwheat sprouts. Korean J Plant Res 24, 724-728

Xu, H., Xiong, A.S., Zhao, W., Tian, Y.S., Peng, R.H., Chen, J.M., and Yao, Q.H. (2011). Characterization of a glucose-, xylose-, sucrose-, and d-galactose- stimulated β-glucosidase from the alkalophilic bacterium Bacillus halodurans C-125. Curr. Microbiol. 62, 833–839.

Xue, Y.M., Xu, C.Y., Hou, J.J., Li, X.Q., and Cao, Z.G. (2015). Enhanced Soluble Expression of a Thermostble β glucosidase from Thermotoga maritima in Escherichia coli and its Applicaton in Immobilization. Appl. Biochem. Microbiol. 51, 306–315.

Yabuuchi, E., and Kosako, Y. (2015). Sphingomonas. In Bergey’s Manual of Systematics of Archaea and Bacteria.

Yang, F., Yang, X., Li, Z., Du, C., Wang, J., and Li, S. (2015). Overexpression and characterization of a glucose-tolerant β-glucosidase from T. aotearoense with high specific activity for cellobiose. Appl. Microbiol. Biotechnol. 99, 8903–8915.

Yang, S.J., Kataeva, I., Wiegel, J., Yin, Y., Dam, P., Xu, Y., Westpheling, J., and Adams, M.W.W. (2010). Classification of “Anaerocellum thermophilum” strain DSM 6725 as Caldicellulosiruptor bescii sp. nov. Int. J. Syst. Evol. Microbiol. 60, 2011–2015.

Yernool, D.A., McCarthy, J.K., Eveleigh, D.E., and Bok, J.D. (2000). Cloning and characterization of the glucooligosaccharide catabolic pathway β-glucan glucohydrolase and cellobiose phosphorylase in the marine hyperthermophile thermotoga neapolitana. J. Bacteriol. 182, 5172–5179.

Yoo, W., Lee, C.W., Kim, B. young, Huong Luu Le, L.T., Park, S.H., Kim, H.W., Shin, S.C., Kim, K.K., Lee, J.H., and Kim, T.D. (2019). Structural and functional analysis of a dimeric fumarylacetoacetate hydrolase (EaFAH) from psychrophilic Exiguobacterium antarcticum. Biochem. Biophys. Res. Commun. 509, 773–778.

Young, J.M., Kerr, A., and Sawada, H. (2015). Agrobacterium. In Bergey’s Manual of Systematics of Archaea and Bacteria.

Yu, H.L., Xu, J.H., Lu, W.Y. and Lin G.Q. (2007). Identification, purification and characterization of β -glucosidase fromapple seed as a novel catalyst for synthesis of O-glucosides. Enzyme Microb Tech 40, 354-361

Zahoor, S., Javed, M.M., Aftab, M.N., and Ikram-Ul-Haq (2011). Cloning and expression of β-glucosidase gene from Bacillus licheniformis into E. coli BL 21 (DE3). Biologia (Bratisl). 66, 213–220.

Zanphorlin, L.M., De Giuseppe, P.O., Honorato, R.V., Tonoli, C.C.C., Fattori, J., Crespim, E., De Oliveira, P.S.L., Ruller, R., and Murakami, M.T. (2016). Oligomerization as a strategy for cold adaptation: Structure and dynamics of the GH1 β-glucosidase from Exiguobacterium antarcticum B7. Sci. Rep. 6, 23776.

Zhang, D., Lax, A.R., Bland, J.M., Yu, J., Federova, N. and Nierman W.C. (2010). Hydrolysis of filter‐paper cellulose to glucose by two recombinant endogenous glycosyl hydrolases of Coptotermes formosanus. Insect Sci 17, 245-252

Zhao, L., Xie, J., Zhang, X., Cao, F., and Pei, J. (2013a). Overexpression and characterization of a glucose-tolerant β-glucosidase from Thermotoga thermarum DSM 5069T with high catalytic efficiency of ginsenoside Rb1 to Rd. J. Mol. Catal. B Enzym. 95, 62–69.

Zhao, W., Peng, R., Xiong, A., Fu, X., Tian, Y., and Yao, Q. (2012). Expression and characterization of a cold-active and xylose-stimulated β-glucosidase from Marinomonas MWYL1 in Escherichia coli. Mol. Biol. Rep. 39, 2937–2943.

Zhao, Z., Ramachandran, P., Kim, T.S., Chen, Z., Jeya, M. and Lee, J.K. (2013). Characterization of an acid-tolerant β-1,4-glucosidase from Fusarium oxysporum and its potential as an animal feed additive. Appl Microbiol Biotechnol 97, 10003-10011

Zhou, C., Tokuhisa, J.G., Bevan, D.R., and Esen, A. (2012). Properties of β-thioglucoside hydrolases (TGG1 and TGG2) from leaves of Arabidopsis thaliana. Plant Sci. 191–192, 82–92.

Zou, Z., Yu, H., Li, C., Zhou, X., Hayashi, C., and Sun, J. (2012). A new thermostable β-glucosidase mined from Dictyoglomus thermophilum : Properties and performance in octyl glucoside synthesis at high temperatures. 118, 425–430.

**Table S3.** Catalytic parameters for modern and ancestral glycosidases determined from the fits of the Michaelis-Menten equation to the rate versus substrate concentration profiles of Figure 5. The errors are standard deviations derived from the fits. All values correspond to 25 ºC and pH 7 (HEPES buffer 50 mM).

|  | 4-nitrophenyl β-D-  glucopyranoside | | | 4-nitrophenyl β-D-galactopyranoside | | |
| --- | --- | --- | --- | --- | --- | --- |
|  | *k_cat_* (s^-1^) | *K_M_* (mM) | *k_cat_/K_M_*  (s^-1^ mM^-1^) | *k_cat_* (s^-1^) | *K_M_* (mM) | *k_cat_/K_M_*  (s^-1^ mM^-1^) |
| Ancestral | 0.101 ± 0.005 | 2.7 ± 0.3 | 0.038 ± 0.003 | 0.07 ± 0.01 | 11.4 ± 3 | 0.0062 ±0.0006 |
| Ancestral with bound heme | 0.339 ± 0.001 | 9.11 ± 0.06 | 0.037 ± 0.001 | 0.210 ±0.008 | 15.1 ± 0.8 | 0.0139 ± 0.0003 |
| *Halothermothrix orenii* | 13.0 ± 0.5 | 0.37 ± 0.08 | 35 ± 6 | 52 ± 5 | 11 ± 2 | 4.9 ± 0.3 |
| *Thermotoga maritima* | 18.1 ± 0.5 | 0.27 ± 0.03 | 68 ± 8 | 28.6 ± 0.5 | 5.7 ± 0.2 | 5.0 ± 0.1 |
| *Marinomonas sp.* (strain MWYL1) | 21.9 ± 0.5 | 0.19 ± 0.02 | 114 ± 12 | 31 ± 7 | 14 ± 5 | 2.1 ± 0.3 |
| *Saccharophagus degradans* (strain 2-40_T_) | 27.6 ± 0.8 | 0.41 ± 0.06 | 66 ± 8 | 12 ± 1 | 28 ± 4 | 0.43 ± 0.01 |

**Table S4.** Amino acid residues at critical active site positions in modern and ancestral glycosidases. The catalytic carboxylic acids, as well as the positions involved in the binding of the glycone and aglycone moieties of the substrate (Marana et al., 2016) are shown. Residues at those positions in the ancestral glycosidase and the modern glycosidase from *Halothermothrix orenii* are given. The last column provides the residue statistics for the set of modern glycosidases used as starting point for ancestral sequence reconstruction. Glycosidases are known to be somewhat specific for the glycone moiety of the substrate and much less specific for the aglycone moiety, which is reflected in a lower residue conservation at the protein residues involved in aglycone binding. See Figure S6 for a graphical illustration.

|  | Ancestral glycosidase | Modern glycosidase from *Halothermothrix orenii* | Sequence statistics in the set of modern glycosidases used as a starting point for ancestral sequence reconstruction. The residue present in the ancestral protein is highlighted in bold |
| --- | --- | --- | --- |
| Catalytic residues | E171 | E166 | **E 98%** Q 0.7% R 0.7% S 0.7% |
|  | E358 | E354 | **E 100%** |
| Residues involved in binding the glycone part of the substrate | Q25 | Q20 | **Q 98.7%** H 0.7% P 0.7% |
|  | H126 | H121 | **H 97.3%** W 1.3% D 0.7% N 0.7% |
|  | N170 | N165 | **N 99.3%** C 0.7% |
|  | W404 | W401 | **W 98.7%** M 0.7% R 0.7% |
|  | E411 | E408 | **E 84.7%** S 14.7% R 0.7% |
|  | W412 | W409 | **W 86.7%** A 5.3% F 2.7% L 1.3% T 1.3% I 0.7% M 0.7% S 0.7% V 0.7% |
| Residues involved in binding the aglycone part of the substrate | W40 | W35 | **W 85.3%** S 2.7% V 2.7% A 2.0% I 2.0% F 1.3% L 1.3% M 1.3% C 0.7% T 0.7% |
|  | F43 | F38 | **F 36%** W 36% Y 6.7% L 5.3% A 4.7% M 2.7% E 1.3% G 1.3% K 1.3% Q 1.3% V 1.3% I 0.7% S 0.7% T 0.7% |
|  | W127 | W122 | **W 50.7%** F 39.3% Y 7.3% G 0.7% H 0.7% L 0.7% S 0.7% |
|  | F175 | V170 | Y 21.3% V 18.0% **F 17.3%** S 12.0% P 6.0% I 4.7% A 4.0% M 3.3% L 2.7% Q 2.7% T 2.7% Q 2.7% T 2.7% W 2.7% N 2% C 0.7% |
|  | L178 | E173 | **L 30.7%** M 10.0% G 8.0% N 8.0% A 6.7% Q 6.7% H 4.7% F 4.0% E 3.3% C 2.7% K 2.7% S 2.7% Y 2.0% D 1.3% P 1.3% R 1.3% T 1.3% V 1.3% W 0.7% |
|  | H185 | H180 | **H 26.7%** W 21.3% F 20.7% L 5.3% Y 4.0% K 3.3% M 2.7% S 2.7% G 2.0% I 2.0% Q 2.0% R 2.0% A 1.3% D 1.3% V 1.3% E 0.7% N 0.7% |
|  | N227 | N222 | **N 48.7%** A 16.0% H 8.0% S 6.0% D 4.7% L 3.3% I 2.0% Q 2.0% T 2.0% V 2.0% F 1.3% Y 1.3% C 0.7% E 0.7% G 0.7% R 0.7% |
|  | W332 | W327 | **W 86%** Y 6.7% E 1.3% L 1.3% F 0.7% G 0.7% H 0.7% I 0.7% R 0.7% S 0.7% V 0.7% |
|  | A413 | A410 | **A 40.7%** S 8.0% E 6.7% L 6.7% T 6.7% D 5.3% G 5.3% H 4.0% I 3.3% N 3.3% R 2.0% V 2.0% F 1.3% P 1.3% Q 1.3% C 0.7% K 0.7% M 0.7% |
|  | F420 | F417 | **F 80.0%** Y 18.0% L 1.3% R 0.7% |

**Table S5.** Atomic surface area values for heme bound to the ancestral glycosidase and the bound heme upon mutating to alanine in silico residues that block its access to the active sate (Pro172, Asn173, Ile224, Leu226, Asn227 and Pro 272). As reference, the values for free heme are given in the last column.

| Atom | Ancestral | Ancestral with mutations to alanine | Free heme |
| --- | --- | --- | --- |
| NB | 0 | 4.9 | 4.9 |
| ND | 0 | 1.9 | 4.8 |
| C1 | 0 | 4.2 | 6.6 |
| C1 | 0 | 3.7 | 6.3 |
| C1 | 0 | 3.8 | 6.5 |
| C1 | 0 | 3.2 | 6.7 |
| C2 | 0 | 2.8 | 3 |
| C2 | 0.1 | 2.6 | 5.5 |
| C2 | 0 | 2.6 | 6 |
| C2 | 0 | 1.7 | 4.7 |
| C3 | 0 | 2.7 | 4.8 |
| C3 | 0 | 0 | 4.5 |
| C3 | 0 | 2.6 | 5.5 |
| C3 | 0 | 0.2 | 3.2 |
| C4 | 0 | 3.3 | 6.5 |
| C4 | 0 | 2.9 | 6.4 |
| C4 | 0 | 5.9 | 6.6 |
| C4 | 0 | 2.5 | 6 |
| CA | 0 | 20.6 | 20.6 |
| CA | 0 | 2.6 | 24.5 |
| CA | 5.6 | 12.8 | 25.2 |
| CA | 0.1 | 0.1 | 16.8 |
| CB | 0.1 | 4.8 | 20.4 |
| CB | 0.2 | 0.2 | 58.6 |
| CB | 10.7 | 20.9 | 58.5 |
| CB | 0 | 17 | 20.6 |
| CG | 0 | 2.1 | 7.8 |
| CG | 0 | 5.5 | 8.2 |
| CH | 0 | 3.9 | 8.4 |
| CH | 0 | 11.3 | 15.1 |
| CH | 0 | 5.7 | 12 |
| CH | 0.4 | 6.2 | 13.6 |
| CM | 0 | 37.3 | 54.6 |
| CM | 0 | 2.6 | 54 |
| CM | 0 | 4 | 52.4 |
| CM | 2.5 | 3.1 | 52.8 |
| NA | 0 | 4.2 | 5.1 |
| NC | 0 | 4.4 | 5.7 |
| O1 | 10.8 | 39.9 | 52.1 |
| O1 | 4.8 | 21 | 43.8 |
| O2 | 4.6 | 15.7 | 37.3 |
| O2 | 3.1 | 16.9 | 50 |
| FE | 0 | 5.2 | 5.2 |

**Table S6.** Data collection and refinement statistics (values in parentheses are for highest-resolution shell).

| Protein | ancestral | ancestral-heme |
| --- | --- | --- |
| PDB ID | 6Z1H | 6Z1M |
| Space group | P 21 | P 21 |
| Unit cell  a, b, c (Å)  β (°) | 52.26 80.67 97.81  100.06 | 58.93 89.49 141.12  94.211 |
| ASU | 2 | 3 |
| Resolution (Å) ^*^ | 43.38 - 2.5 (2.59 - 2.5) | 52.87 - 2.45 (2.54 - 2.45) |
| R*_merge_* (%)^*^ | 8.90 (105.30) | 80.96 (80.85) |
| I/σ_I_ ^*^ | 10.9 (1.7) | 10.15 (1.44) |
| Completeness (%)^*^ | 99.92 (99.93) | 99.02 (99.57) |
| Unique reflections^*^ | 27829 (2775) | 53447 (5335) |
| Multiplicity | 7.4 (7.5) | 3.2 (3.2) |
| Wilson B-factor | 59.93 | 50.34 |
| CC(1/2) ^*^ | 0.999 (0.707) | 0.997 (0.718) |
| **Refinement** |  |  |
| R*_work_*/R*_free_* (%) | 19.59 / 24.10 | 17.55 / 22.09 |
| No. atoms | 6601 | 11026 |
| Protein | 6546 | 10676 |
| Ligands | 21 | 194 |
| Solvent | 34 | 156 |
| B-factor (Å^2^) | 76.63 | 62.81 |
| R.m.s deviations |  |  |
| Bond lengths (Å) | 0.003 | 0.005 |
| Bond angles (^0^) | 0.65 | 0.73 |
| Ramachandran (%) |  |  |
| Favored | 96.09 | 96.14 |
| Outliers | 0.13 | 0 |

^*^Statistics for the highest-resolution shell are shown in parentheses.

**Supplementary FIgures**

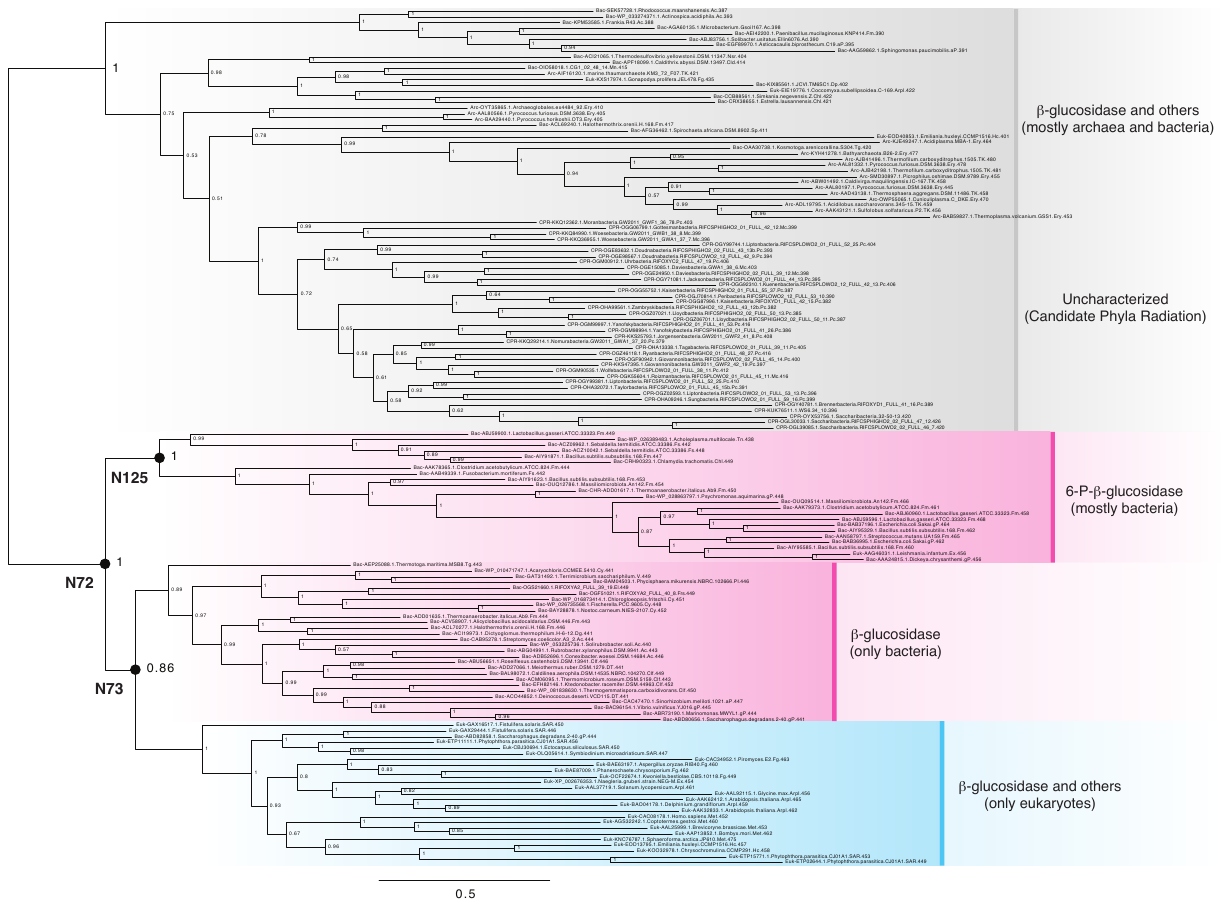

**Figure S1.** Bayesian analysis of family 1 glycosidases (GH1) protein sequences with sequence annotations. The annotation includes the accession number, the taxonomical information (domain name, phylum name and species name) and the sequence length. Three black dots indicate three ASR nodes (N72, N73 and N125). Scale bar represents 0.5 amino acid replacements per site per unit evolutionary time. Abbreviations: Ac = Actinobacteria; aP = α-Proteobacteria; Arc = Archaea; Arpl = Archaeplastida; Asg = Asgard group; Bac = Bacteria; Bc = Bacteroidetes; bP = β-Proteobacteria; Chl = Chlamydiae; Cld = Calditrichaeota (Caldithrix); Clf = Cloroflexi; CPR = Candidate Phyla Radiation; Cy = Cyanobacteria; Euk = Eukaryotes; Ex = Excavates; Fg = Fungi; Fm = Firmicutes; Frs = Fraserbacteria; Dg = Dictyoglomi; dP = δ-Proteobacteria; Dp = Dependentiae (TM6); DPN = DPANN group; DT = Deinococcus-Thermus; Ery = Euryarchaeota; Fs = Fusobacteria; gP = γ-Proteobacteria; Hc = Hacrobia; Mc = Microgenomates; Met = Metazoa; Mn = Marinimicrobia; Nsr = Nitrospirae; Pc = Parcubacteria; Pl = Planctomycetes; V = Verrucomicrobia; SAR = SAR group; Sp = Spirochaetes; Tg = Thermotogae; TK = TACK group; Tn = Tenericutes.

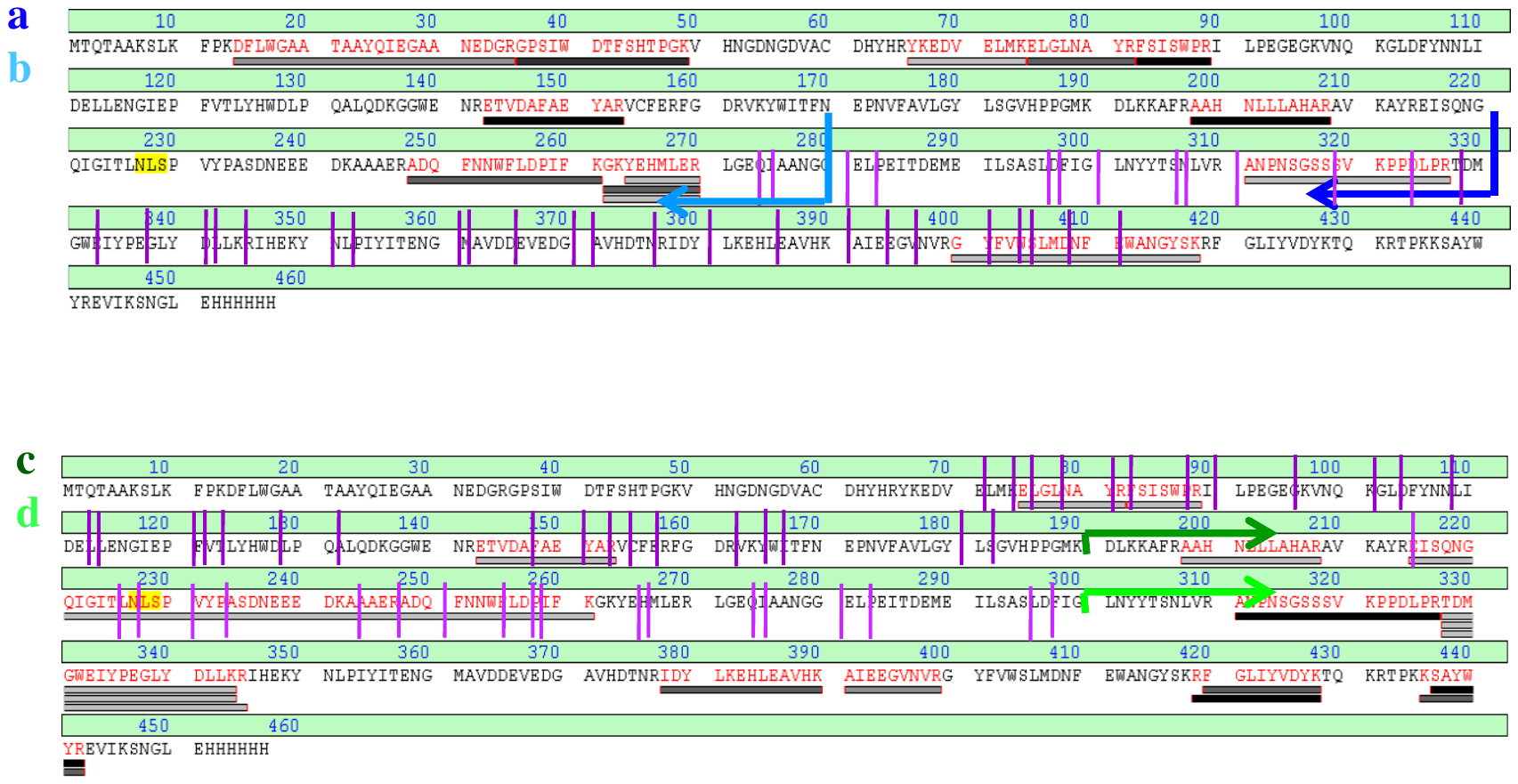

**Figure S2.** Estimation of the location of thermolysin cleavage sites from mass spectrometry and peptide-mapping fingerprinting. Potential thermolysin restriction sites are shown by vertical purple lines. Four thermolysin fragments were studied (a, b, c, and d: see left for color code). Fragment masses were determined by MALDI and their sequences were investigated using peptide mapping finger-printing and MALDI-TOF/TOF. Sequences for several sub-fragments (shown) could be thus determined and the length of the original fragments could be assessed. Fragments a and b extend approximately from the amino terminus to the dark and light blue arrows in the upper panel. Fragments c and d extend approximately from the dark and light green arrows in the lower panel to the carboxyl terminus. Comparison with the restriction sites allows a determination of the plausible thermolysin cleavage sites, as shown in Figure 2C of the main text.

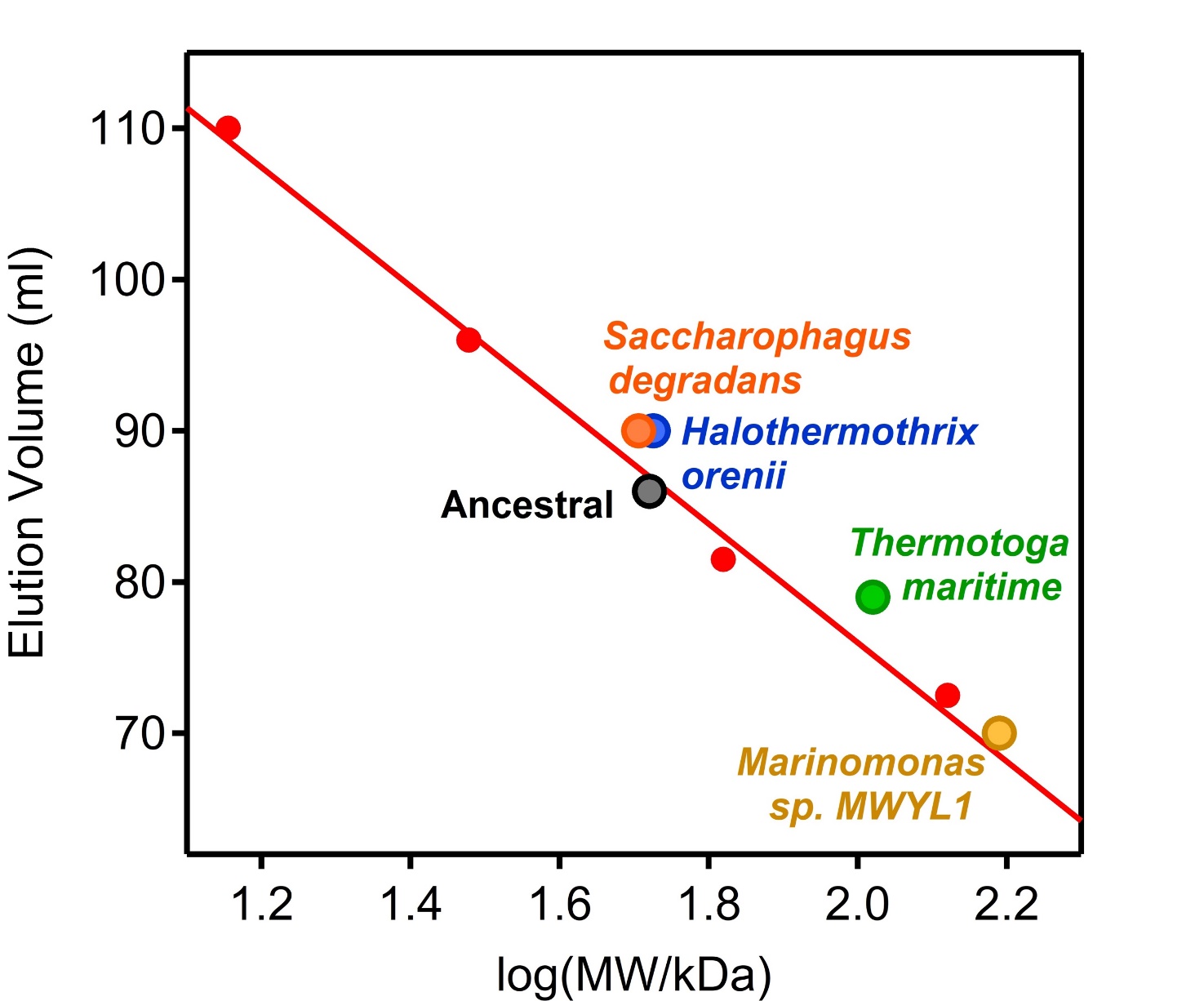

**Figure S3.** Assessment of association state of modern and ancestral glycosidases through gel filtration chromatography (HiLoad 16/600 Superdex 200 pg GE Healthcare). The molecular mass (MW) was estimated by the calibration curve of elution volume vs. log (MW). The protein markers used were: bovine serum albumin (monomer: 66 kDa, dimer: 132 kDa), deoxyribonuclease I from bovine pancreas (30.1 kDa) and lysozyme (14.3 kDa). The ancestral glycosidase and the modern glycosidases from *Saccarophagus degradans and Halothermothrix orenii* are monomers. The modern glycosidases from *Thermotoga maritima* is a dimer and the modern glycosidase from *Marinomonas sp. MWTL1* is a trimer.

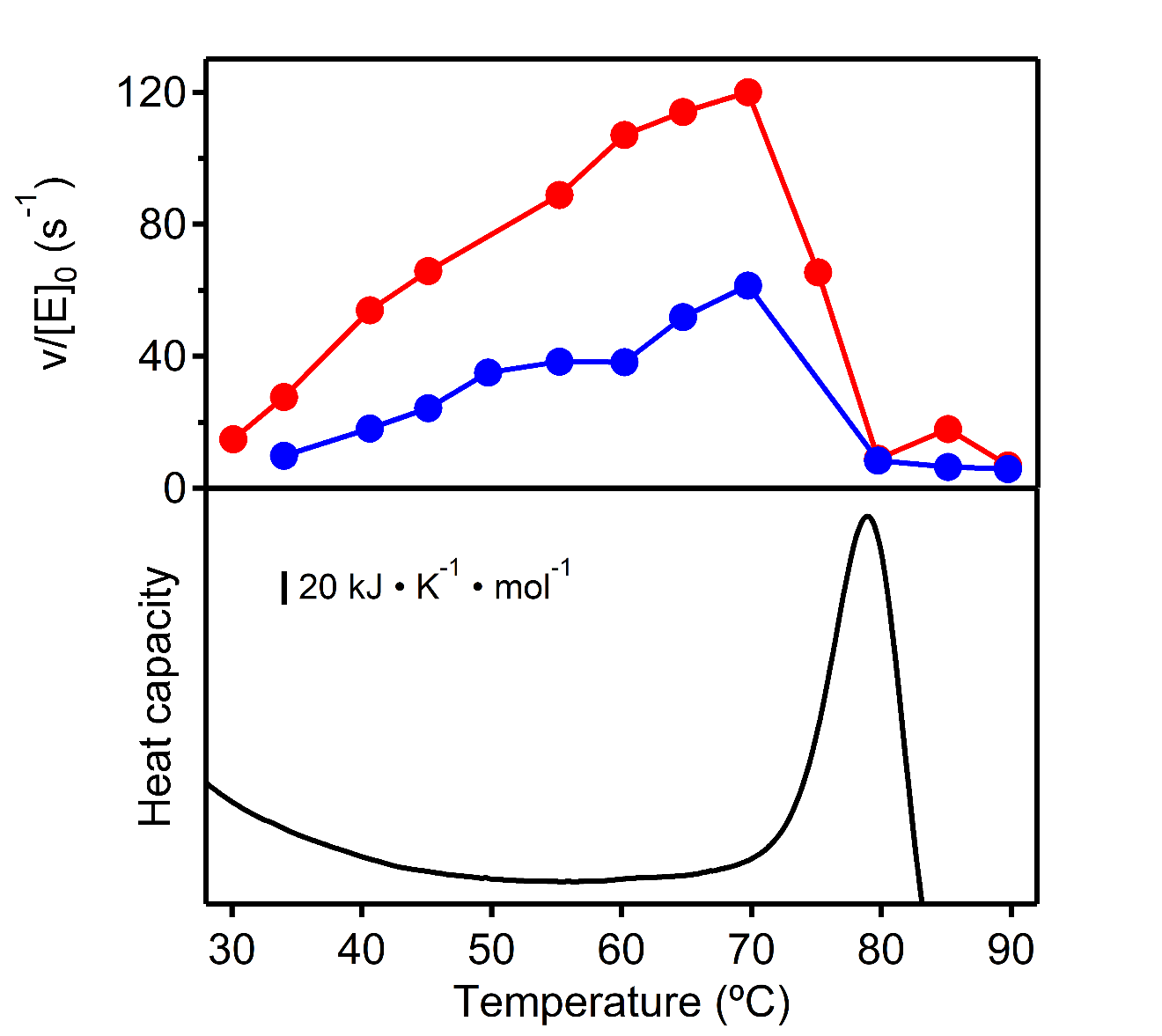

**Figure S4.** Determination of the optimum temperature for the modern glycosidase from *Halothermothrix orenii* using two different substrates 4-nitrophenyl-β-D-glucopyranoside (red) and 4-nitrophenyl-β-D-galactopyranoside (blue). The lower panel shows a differential scanning calorimetry profile for the enzyme under the same buffer conditions. Clearly, the activity drop observed at high temperature (upper panel) corresponds to the denaturation of the protein, as seen in the lower panel.

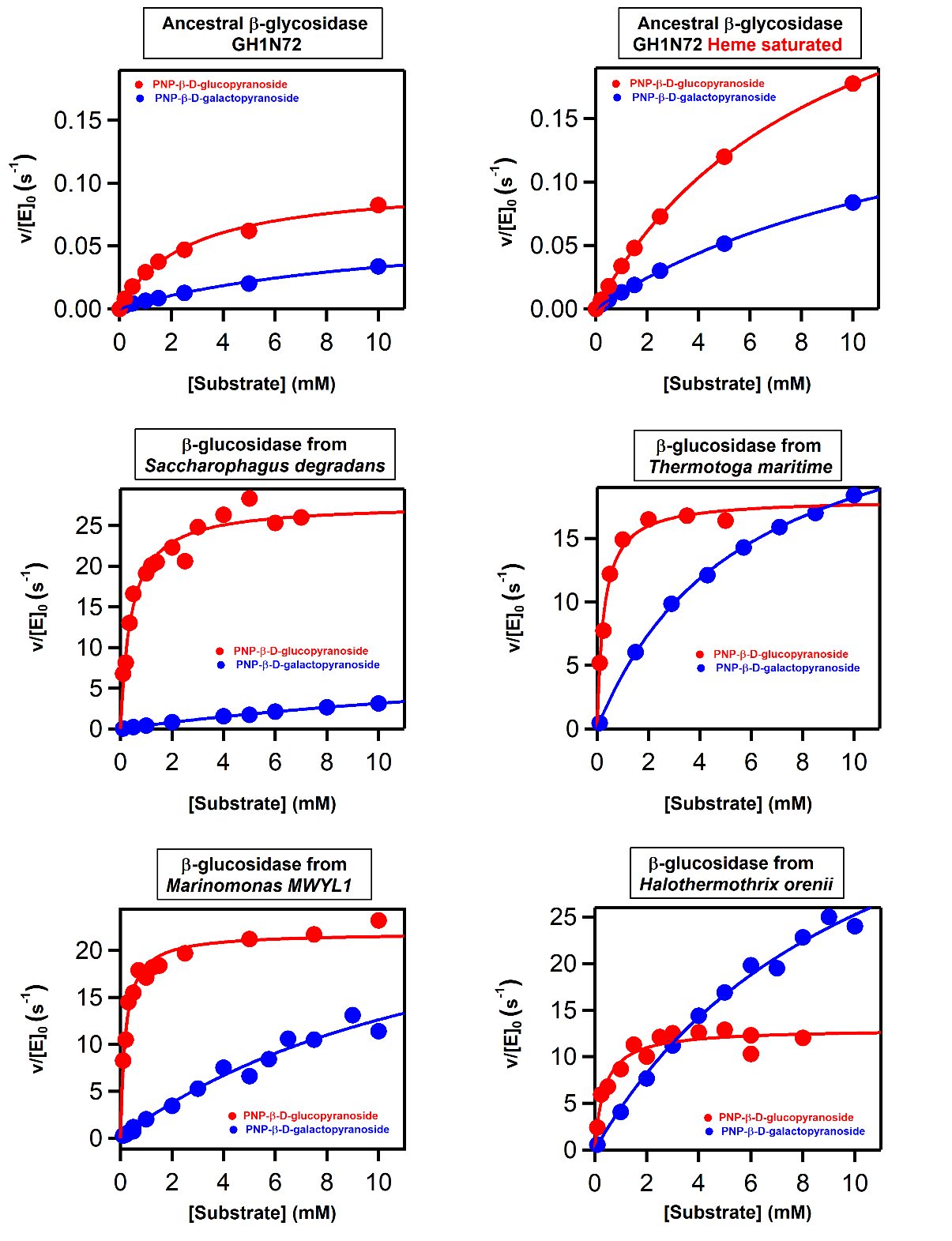

**Figure S5.** Michaelis plots for ancestral and modern glycosidases. Data is provided for glucopyranoside and galactopyranoside substrates. The continuous lines are the best fits of the Michaelis-Menten equation. The catalytic parameters derived from these fits are provided in Table S3 and Figure 4 of the main text. Note that y-axis range is the same in the plots for the ancestral protein without and with bound heme, to make visually clear that heme binding enhances catalysis by several-fold. Note also that the studied modern glycosidases bind the glucopyranoside substrate more tightly than the galactopyranoside substrates. This preference is commonly observed for family 1 glycosidases (Withers, 2012).

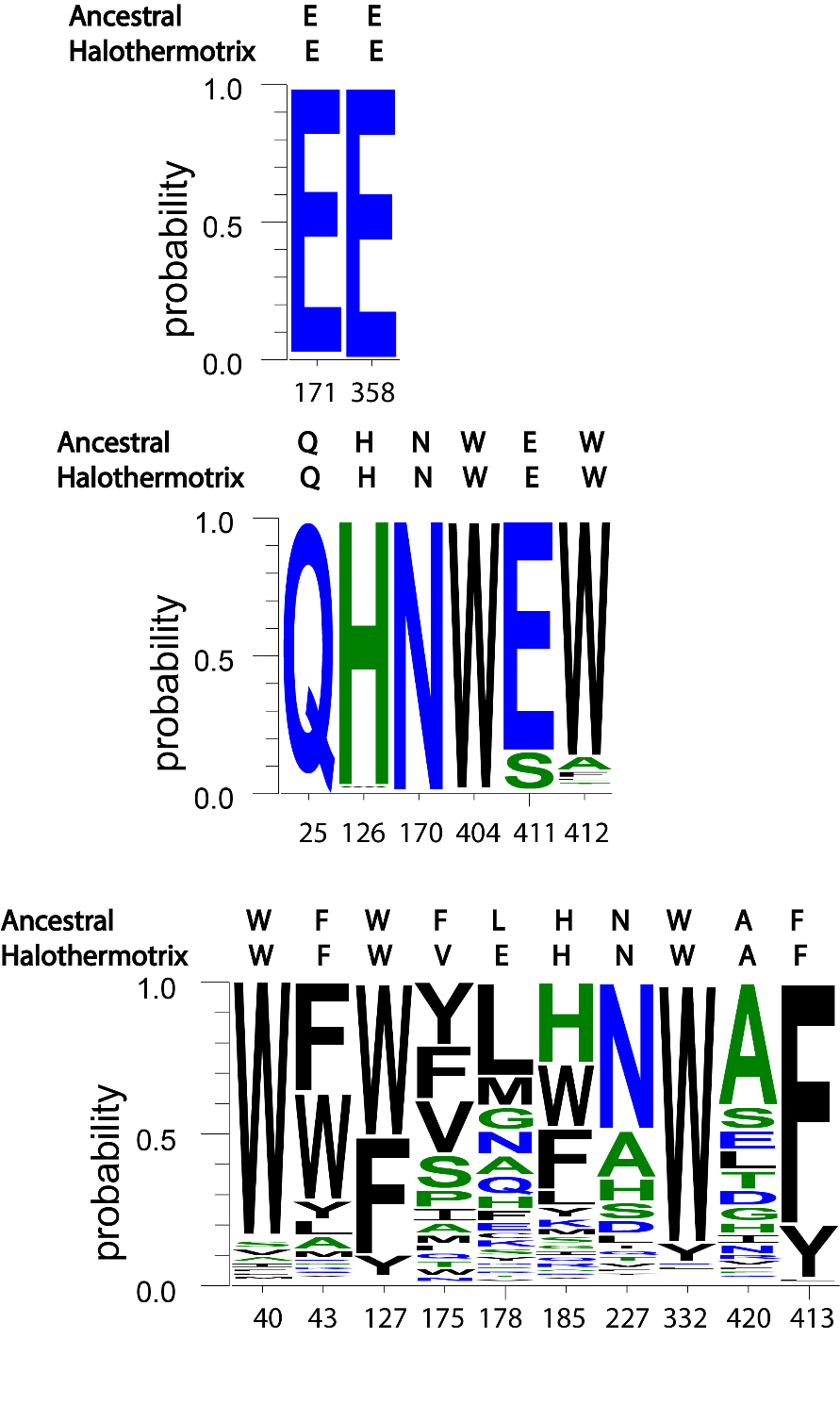

**Figure S6.** Statistics of residue occupancy at critical active site positions in the set of modern glycosidases used as starting point for ancestral sequence reconstruction (see also Table S4). The graphics shown refer to the catalytic carboxylic acids (upper), the positions involved in the binding of the glycone moiety of the substrate (middle) and the positions involved in the binding of the aglycone moiety of the substrate (lower). The sequences of the ancestral glycosidase and the modern glycosidase from *Halothermothrix orenii* are also given. Glycosidases are known to be somewhat specific for the glycone moiety of the substrate and much less specific for the aglycone moiety, which is reflected in lower residue conservation at the protein residues involved in aglycone binding.

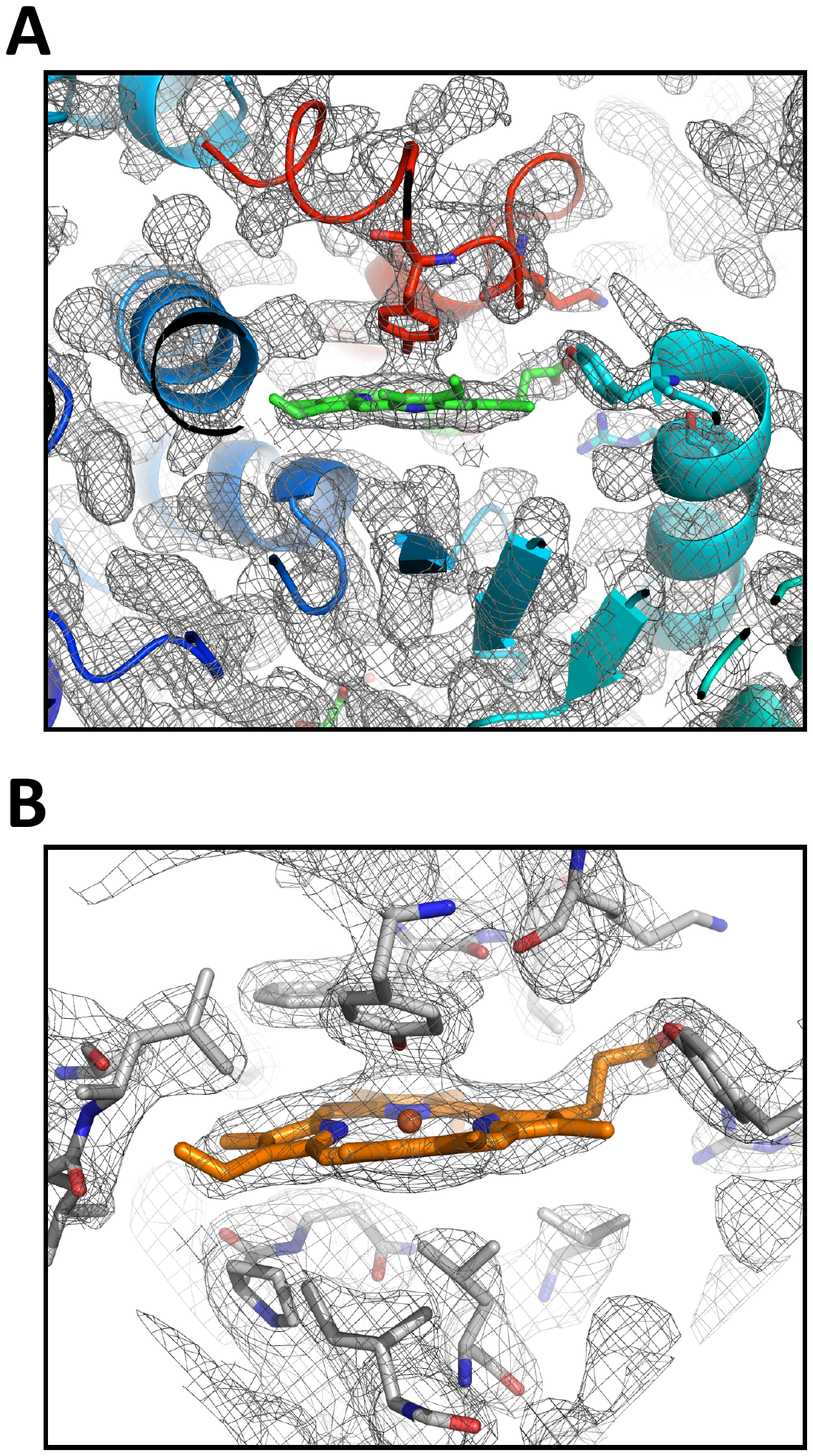

**Figure S7. (**A) Structure of the ancestral glycosidase showing the bound heme group into the well-defined |2Fo-Fc| electron density map contoured at 1 σ. The four amino acids involved in hydrogen bonds (See Figure 6A) are shown as sticks. B) Blow-up showing all residue participating in binding the heme group .

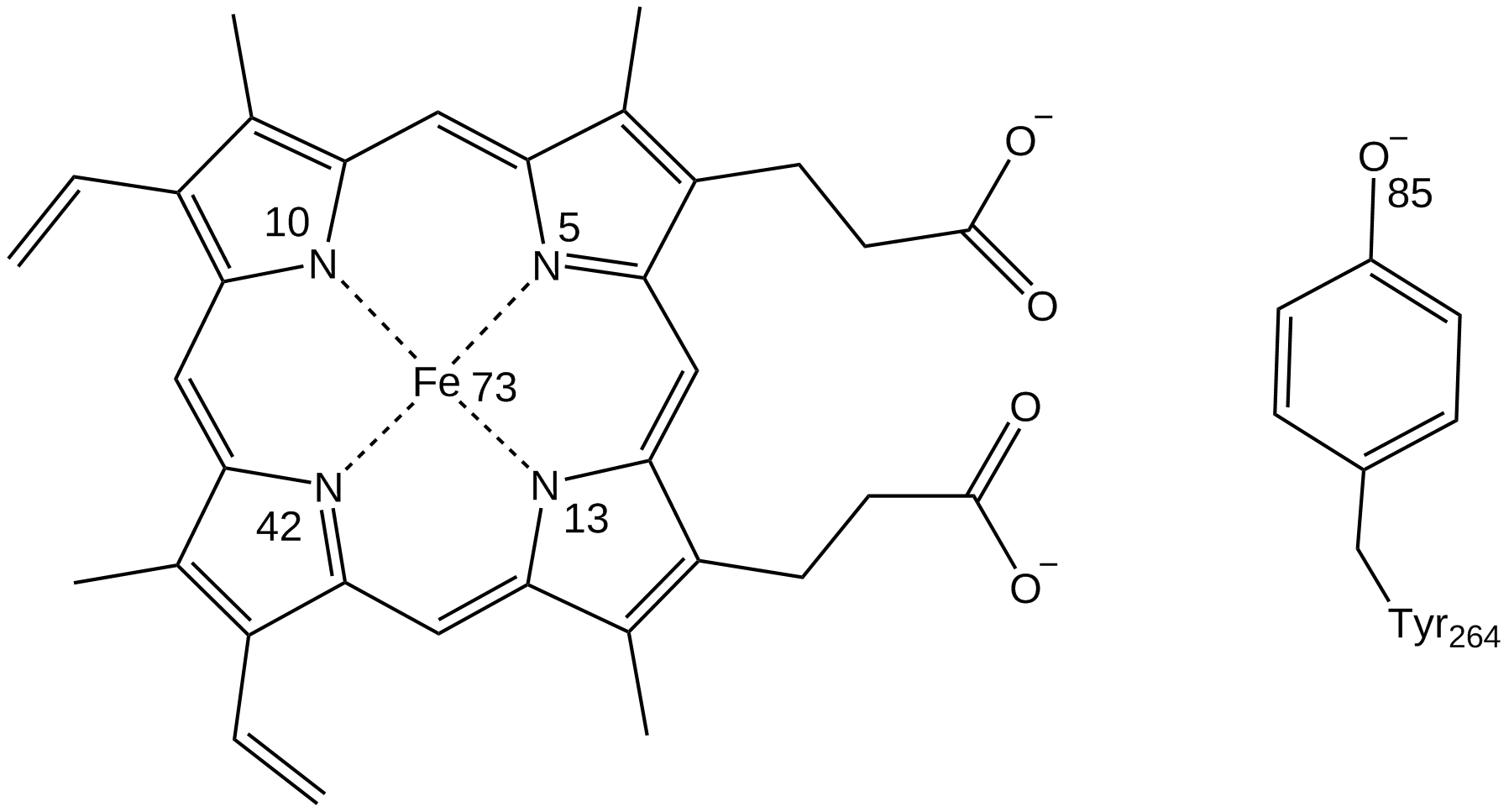

**Figure S8.** Atom numbering of the atoms considered in the bonded model between the Fe and the heme and Tyr264.

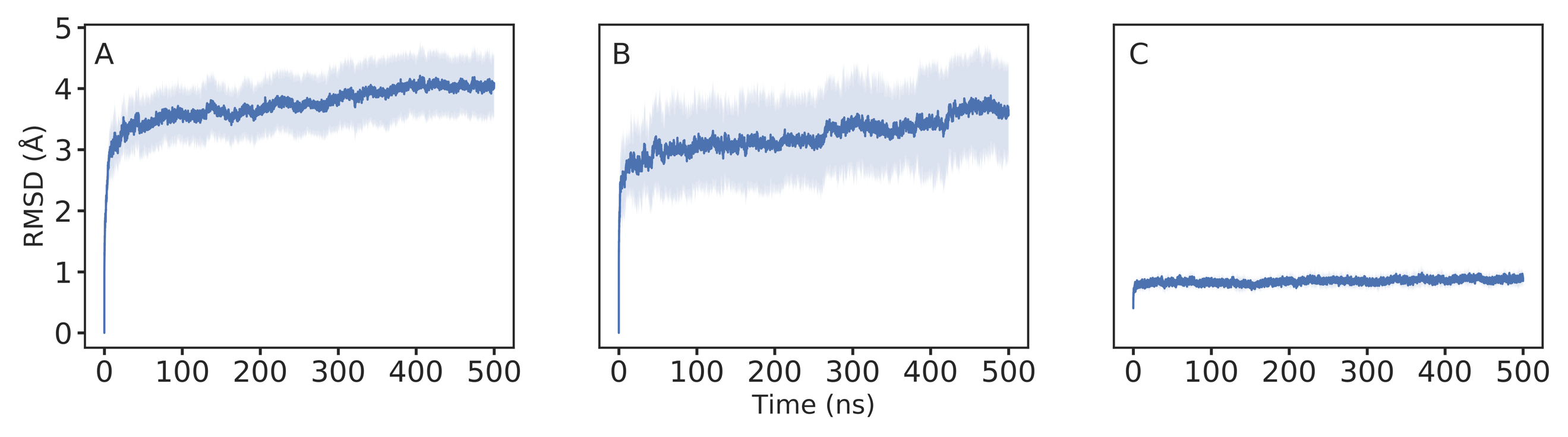

**Figure S9.** The root mean square deviations (RMSD, Å) of all backbone atoms of the (A) ancestral glycosidase without heme bound, (B) ancestral glycosidase with heme bound and (C) modern glycosidase from *Halothermothrix orenii* over ten individual 500 ns MD simulations per system (i.e. 5 μs cumulative simulation time per system). The average RMSD per system is denoted by solid blue lines, and the standard deviations per point over all trajectories are illustrated by the shaded area on each plot.
